# Small GTPases RhoA and RhoC coordinately regulate cardiomyocyte size in the mouse heart

**DOI:** 10.64898/2026.10.12.758623

**Authors:** Vikram Prasad, Naga Swathi Sree Kavuri, Max Toubin, J. Matthew Kofron, Yi Zheng, Jop H. van Berlo, Jeffery D. Molkentin, Davy Vanhoutte

**Affiliations:** Division of Molecular Cardiovascular Biology, Heart Institute, Cincinnati Children’s Hospital Medical Center, Cincinnati, OH 45229, USA; Department of Pediatrics, University of Cincinnati, Cincinnati Children’s Hospital Medical Center, Cincinnati, OH 45229, USA; Division of Developmental Biology, Cincinnati Children’s Hospital Medical Center, Cincinnati, OH 45229, USA; Division of Experimental Hematology and Cancer Biology, Cancer and Blood Diseases Institute, Cincinnati Children’s Hospital Medical Center, Cincinnati, OH 45229, USA; Lillehei Heart Institute, Cardiovascular Division, University of Minnesota, Minneapolis, MN 55455, USA

**Keywords:** RhoA, RhoC, cardiomyocyte hypertrophy, cardiac remodeling, small GTPase, pressure overload

## Abstract

Rho family GTPases regulate cardiomyocyte growth and cytoskeletal organization, yet the overlapping and individual function of the two primary isoforms RhoA and RhoC, remain largely unknown in the heart. Here, we examined cardiac structure and function at baseline and in response to mechanical and neurohumoral stress in mice with cardiomyocyte-specific deletion of *Rhoa*, global loss of *Rhoc*, or combined deficiency of both isoforms. Deletion of either *Rhoa* or *Rhoc* increased myocardial expression of the other isoform, showing reciprocal compensation. At baseline, *Rhoc* deficiency produced elongated, narrower cardiomyocytes, whereas additional loss of *Rhoa* restored cell width but length remained longer. Combined *Rhoa/Rhoc* deficiency also disrupted intercalated disc protein expression and organization and caused age-dependent cardiac hypertrophy with mild pulmonary congestion despite preserved systolic function. Following pressure overload, loss of either isoform increased hypertrophy and fibrosis compared with control groups, while combining *Rhoa/Rhoc* deficiency significantly worsened cardiac remodeling and systolic dysfunction. Neurohumoral stimulation with angiotensin II and phenylephrine similarly revealed cooperative control of hypertrophic growth in *Rhoa/Rhoc* double deleted cardiomyocytes versus either single deletion. These findings identify RhoA and RhoC as partially compensatory effectors of cardiomyocyte growth and ventricular remodeling with stress stimulation, although their individual contributions to hypertrophic growth, structural integrity, and cardiac function depend on both the nature and duration of cardiac stress.

**KEY POINTS SUMMARY:**

- Cardiac hypertrophy is regulated by small GTPase proteins including RhoA, but the role of the closely related protein RhoC in the heart remains poorly defined.
- Using mice with genetic deletion of *Rhoa* and *Rhoc*, we show that they compensate for one another, and that loss of *Rhoc* alone alters the size and shape of cardiomyocytes even without added stress.
- Loss of *Rhoc*, but not *Rhoa*, reduced heart function within two weeks of a sudden increase in workload, whereas *Rhoa* was important with prolonged workload.
- Combined loss of *Rhoa* and *Rhoc* in cardiomyocytes produces significantly greater hypertrophy to neurohumoral signals compared with deletion of either gene alone.
- These findings suggest that RhoA and RhoC play both overlapping and non-redundant roles in the heart’s response to stress, with implications for understanding and treating cardiac hypertrophy and failure.

## INTRODUCTION

Cardiac hypertrophy is a compensatory adaptive response to increased mechanical load, neurohumoral stimulation, or myocardial injury that serves to limit wall stress and preserve cardiac function (van Berlo *et al*., 2013; Nakamura & Sadoshima, 2018). However, sustained pathological stimulation can drive the transition from compensatory hypertrophy to maladaptive remodeling and ultimately heart failure (Nakamura & Sadoshima, 2018). At the cellular level, pressure overload classically drives concentric hypertrophy with preferential cardiomyocyte thickening, whereas volume overload favors eccentric hypertrophy and cardiomyocyte elongation (Kehat *et al*., 2011; Nakamura & Sadoshima, 2018; Grimes *et al*., 2024). Cardiomyocyte elongation and eccentric remodeling are also recognized features of ventricular dilation and decompensation. These distinct growth patterns reflect coordinated remodeling of the sarcomere and cytoskeleton that shape both the magnitude and the geometric type of cardiomyocyte growth in length versus width (Grimes *et al*., 2024). Small Rho GTPases are central regulators of actin cytoskeletal dynamics and are therefore well positioned to influence these processes (Heasman & Ridley, 2008). Among these, RhoA is the best characterized in the heart, acting through downstream effectors, including Rho-associated kinase (ROCK) and the formin mDia, to regulate actin organization (Heasman & Ridley, 2008; Miyamoto, 2024) and myocyte hypertrophy (Lauriol *et al*., 2014).

In cardiomyocytes, RhoA is activated downstream of hypertrophic agonist receptors that engage signaling pathways involved in cytoskeletal remodeling and transcriptional regulation, including myocardin-related transcription factor (MRTF)/serum response factor (SRF) and Yes-associated protein (YAP) signaling (Appert-Collin *et al*., 2007; Mayers *et al*., 2010; Sakabe *et al*., 2022; Miyamoto, 2024). Angiotensin II activates RhoA in cultured cardiomyocytes to promote premyofibril formation and sarcomere assembly (Aoki *et al*., 1998), while Rho signaling more broadly regulates hypertrophic gene expression downstream of G protein-coupled receptor (GPCR) agonists (Aikawa *et al*., 2000). However, the consequences of RhoA signaling in the myocardium depend on the magnitude, duration, and context of its activation (Lauriol *et al*., 2014; Miyamoto, 2024). In vivo, high-level cardiac-specific expression of wild-type or constitutively active RhoA produces sinus and atrioventricular nodal dysfunction, chamber dilation, and contractile failure (Sah *et al*., 1999). In contrast, modest cardiomyocyte-specific expression of activated RhoA protects against acute ischemia-reperfusion injury through a protein kinase D-dependent pathway (Xiang *et al*., 2011). Similarly, cardiomyocyte-specific *Rhoa* deletion does not impair the initial hypertrophic response to pressure overload but accelerates subsequent ventricular dilation and contractile dysfunction during chronic pressure overload, in part through diminished extracellular signal-regulated kinase 1/2 (ERK1/2) activation and altered calcium handling (Lauriol *et al*., 2014). Notably, *Rhoa* deletion in cardiomyocytes also reduces interstitial fibrosis, indicating both protective and detrimental aspects of myocardial remodeling by RhoA signaling in the heart (Lauriol *et al*., 2014).

In contrast to RhoA, the role of the closely related isoform RhoC in cardiomyocytes remains poorly defined. RhoA and RhoC share approximately 85% amino acid sequence identity and overlap substantially in their upstream regulators and downstream effectors, suggesting that one isoform could compensate for the other (Wheeler & Ridley, 2004; Eckenstaler *et al*., 2022). Evidence for such compensation comes from a genetic mouse model of oncogenic K-Ras-driven lung tumorigenesis, in which conditional *Rhoa* deletion exacerbated tumor formation through a mechanism partly dependent on RhoC, whereas combined *Rhoa/Rhoc* deletion produced a distinct, non-additive outcome (Zandvakili *et al*., 2015). Nevertheless, differences in their hypervariable C-terminal regions contribute to distinct subcellular localization and signaling properties for RhoA and RhoC (Wheeler & Ridley, 2004; Eckenstaler *et al*., 2022). Furthermore, studies in migrating and invasive cells have demonstrated that RhoA and RhoC can engage distinct downstream effectors with sometimes opposing effects on cytoskeletal organization and cell morphology (Vega *et al*., 2011; Ridley, 2013). Whether similar functional overlap and isoform-specific regulation occur in cardiomyocytes remains unknown.

Here, we investigated the individual and combined requirements for RhoA and RhoC in cardiac homeostasis and stress-induced remodeling by combining cardiomyocyte-specific deletion of *Rhoa* with global loss of *Rhoc*. We hypothesized that RhoC makes distinct, non-redundant contributions to cardiomyocyte geometry and cardiac stress adaptation beyond the established protective role of RhoA, and that combined loss of both GTPases produces hypertrophic and functional deficits exceeding either single deletion alone. By examining the individual and combined consequences of *Rhoa* and *Rhoc* deficiency, we sought to better understand the potential therapeutic benefit behind selective versus combined inhibitor strategies.

## METHODS

### Animals

All animal procedures were approved by the Institutional Animal Care and Use Committee (IACUC) of Cincinnati Children’s Hospital Medical Center under protocols IACUC2016-0069, IACUC2018-0047 and IACUC2024-0043. Sample sizes were based on prior experience with these models and on power calculations to detect biologically relevant differences in the primary outcomes while using the minimum number of mice. Whenever feasible, animals were randomized to experimental groups, and investigators were blinded during data acquisition and analysis. Male and female mice were included in approximately equal proportions; data from both sexes were pooled, and sex was not analyzed as a biological variable to reduce total usage. Mice were maintained under controlled environmental conditions (21-22°C, 40-60% humidity, 12-h light/12-h dark cycle) with *ad libitum* access to food and water.

*Rhoa*-LoxP (fl) targeted mice (*Rhoa*^fl/fl^; RRID:MGI:5004976) and constitutive *Rhoc* gene deleted mice (*Rhoc*^-/-^; no RRID available for this in-house-derived line) were provided by Dr. Yi Zheng (Hakem *et al*., 2005; Melendez *et al*., 2011). Transgenic mice expressing Cre recombinase under the control of the β-myosin heavy chain (βMHC, *Myh7* gene) promoter (βMHC-cre; RRID:MGI:3054171) were described previously (Oka *et al*., 2006). *Rhoa*^fl/fl^ mice were crossed with *βMHC*-cre transgenic mice, which had been independently generated on the same mixed *C57BL/6-129/Sv* genetic background, with or without the *Rhoc*-null allele, to generate the five genotypes used in this study: WT^βMHC-cre^ (Cre-only controls), *Rhoa*^fl/fl^ (controls), *Rhoa*^βMHC-cre^ (cardiomyocyte-specific *Rhoa* deletion), *Rhoa*^fl/fl^;*Rhoc*^-/-^ (global *Rhoc* deletion), and *Rhoa*^βMHC-cre^;*Rhoc*^-/-^ (combined *Rhoa*/*Rhoc* deletion). No human subjects or human tissue samples were used in this study.

### Cardiac Hypertrophy Models and Echocardiography

Eight- to ten-week-old mice underwent transverse aortic constriction (TAC) or sham surgery as previously described (Kehat *et al*., 2011; Schips *et al*., 2019). Briefly, mice were anesthetized with 3% isoflurane and intubated using an 18-gauge catheter. Anesthesia was maintained at 1.7% isoflurane using a rodent ventilator throughout the procedure. Following a median sternotomy, the transverse aortic arch was exposed and constricted by tying a 7-0 silk suture around the aorta together with a 27-gauge blunted-needle positioned between the brachiocephalic artery and the left common carotid artery. The needle was then removed, producing a reproducible reduction in aortic lumen diameter. Doppler echocardiography was performed after surgery to verify the pressure gradient across the constriction. Extended-release buprenorphine (Ethiqa XR, Fidelis Animal Health; North Brunswick Township, NJ, 3.25 mg/kg, 1.3 mg/mL) was administered subcutaneously as a single dose following extubation, providing postoperative analgesia for 72 h. Sham-operated mice underwent the identical surgical procedure except that the suture was not tied around the aorta.

Cardiac hypertrophy was also induced by continuous infusion of angiotensin II (Ang II; 432 µg/kg/day; Sigma-Aldrich, St Louis, MO, #A9525) and phenylephrine (PE; 100 mg/kg/day; Sigma-Aldrich, #P6126) using Alzet micro-osmotic pumps (Durect Corp., Cupertino CA, model 1002, #0004317) (Maillet *et al*., 2009). Pumps were filled with either the Ang II/PE solution or vehicle (150 mM NaCl containing 0.01 N acetic acid) using a blunted filling needle and primed overnight at 37°C in vehicle solution before implantation. The following day, eight- to ten-week-old mice were anesthetized with 3% isoflurane, and osmotic pumps were implanted subcutaneously over the thoracic region of the back. Skin incisions were closed using 4-0 silk sutures and extended-release buprenorphine (Ethiqa XR; 3.25 mg/kg, 1.3 mg/mL) was administered subcutaneously as a single dose for postoperative analgesia (72 h). Following both TAC and pump implantation surgeries, mice recovered in a 30°C incubator and were closely monitored until fully ambulatory. Animals exhibiting signs of distress were euthanized as described below (see Histopathology). Surgical incisions were treated with topical antibiotic ointment for two consecutive days following surgery.

Cardiac function was evaluated by transthoracic echocardiography using a Sonos 5500 ultrasound system (Philips Ultrasound, Reedsville PA) equipped with a 15-MHz microprobe. Mice were lightly anesthetized with isoflurane (approximately 2%, to effect), and M-mode measurements were acquired in triplicate and averaged for each animal. All image acquisition and subsequent analysis were performed in a blinded manner (Maillet *et al*., 2009).

### Isolation of Adult Cardiomyocytes

Adult ventricular cardiomyocytes were isolated from 2-month-old mice by Langendorff perfusion as previously described (Kehat *et al*., 2011; Goonasekera *et al*., 2012). Briefly, mice were injected intraperitoneally with heparin (30–50 U; Sigma-Aldrich, #H3149) and anesthetized with isoflurane (1–3% in oxygen). Once anesthesia was confirmed by loss of the pedal withdrawal reflex, hearts were rapidly excised, which served as the method of euthanasia, and were cannulated and perfused with oxygenated Tyrode solution (buffer A) containing 120 mM NaCl, 5.4 mM KCl, 1.2 mM NaH_2_PO_4_, 5.6 mM glucose, 20 mM NaHCO_3_, 1.6 mM MgCl_2_, 10 mM 2,3-butanedione monoxime (BDM), and 5 mM taurine. Buffer A was continuously gassed with 95% O_2_ and 5% CO_2_. Hearts were subsequently digested by perfusion with buffer A supplemented with collagenase type II (1 mg/mL; Worthington Biochemical, Lakewood NJ, #LS004174) and protease type XIV (0.08 mg/mL; Sigma-Aldrich, #P5147-1G) at 37°C for 10-14 min. Following enzymatic digestion, the ventricles were gently dissociated into single-cell suspensions and filtered, after which extracellular Ca^2+^ was gradually restored to a final concentration of 1.8 mM. Cardiomyocytes were then plated onto laminin-coated culture dishes and maintained in serum-free HyClone Medium 199/Earle’s Balanced Salt Solution (EBSS) (Thermo Fisher Scientific, Waltham MA, #MT10060CV) supplemented with 1x penicillin-streptomycin (Sigma-Aldrich, #P0781-100ML) until further analysis.

Cardiomyocyte morphology was evaluated by phase-contrast microscopy using a Nikon Eclipse Ti-S inverted microscope equipped with NIS-Elements Advanced Research (AR) software (Nikon Instruments Inc., Melville, NY; RRID:SCR_014329). Cell length and width were measured in a blinded manner in at least 150 cardiomyocytes per heart from three hearts per genotype (Kehat *et al*., 2011); per-heart means were used for statistical analysis.

### Histopathology

At the indicated experimental endpoints, mice were anesthetized with isoflurane (1–3% in oxygen). Once anesthesia was confirmed by loss of the pedal withdrawal reflex, hearts were excised, followed by cervical dislocation, and heart and lung weights were recorded. Heart weight-to-body weight (HW/BW) and lung weight-to-body weight (LW/BW) ratios were calculated and expressed as mg/g. Hearts were fixed overnight in 4% paraformaldehyde (Electron Microscopy Sciences, Morgantown PA, #15714), dehydrated through graded ethanol, embedded in paraffin, and sectioned at a thickness of 5 µm. Tissue sections were stained with Masson’s trichrome using standard procedures (Khalil *et al*., 2017).

Masson’s trichrome-stained heart sections were imaged using an Olympus BX51 microscope equipped with a Nikon DS-F*i*3 camera and NIS-Elements software (Nikon Instruments; RRID:SCR_014329). For fibrosis quantification and the representative images in Figures 3–5, individual fields covering the entire heart section were acquired with a 20× objective without stitching; the low-magnification images in Figure 1L were acquired with a 2× objective. Before imaging, white balance was set automatically on a tissue-free area of each slide adjacent to the section; no further image adjustments were applied. Fibrosis was quantified from the 20× images using Fiji/ImageJ software (National Institutes of Health; RRID:SCR_003070) (Khalil *et al*., 2017). For each field, tissue area was defined by converting the image to binary and excluding background pixels containing no tissue. The same image was then separated into stain-specific channels using the Color Deconvolution plugin with the built-in Masson trichrome vectors, and the Colour_1 (methyl blue, collagen) channel had a threshold that separates fibrotic from non-fibrotic tissue, using a single fixed threshold applied to all images. For each heart, tissue and non-fibrotic areas were summed across all fields, and fibrosis was calculated as (total tissue area − non-fibrotic area) / total tissue area × 100. Because entire sections were analyzed, including perivascular and epicardial regions, values represent total cardiac fibrosis. Image analysis was performed by an operator blinded to genotype and treatment group. Representative images shown in the figures were selected from experiments that were representative of the overall findings.

**Figure 1.**
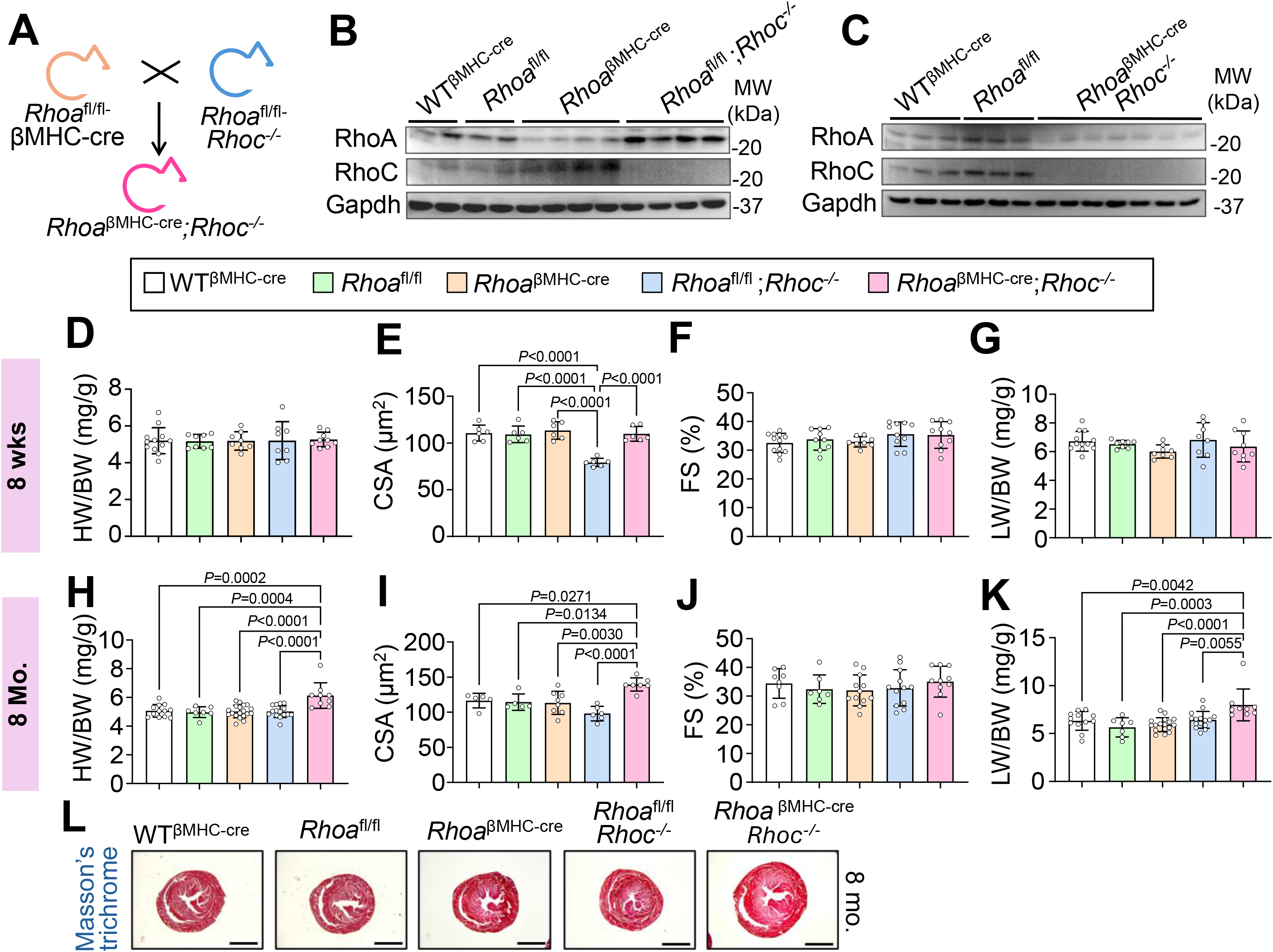
Reciprocal regulation of RhoA and RhoC in the heart and age-dependent remodeling after their combined deletion. (**A**) Breeding scheme to generate mice with cardiomyocyte-specific deletion of *Rhoa* (referred to as *Rhoa*^βMHC-cre^) in a *Rhoc* gene-deleted background (*Rhoc*^-/-^), using *Rhoa*-LoxP (fl)-targeted (*Rhoa*^fl/fl^) mice crossed with *βMHC*-cre transgenic mice. (**B-C**) Western blot analysis of RhoA and RhoC protein levels in 8-week-old WT^βMHC-cre^, *Rhoa*^fl/fl^, *Rhoa*^βMHC-cre^, *Rhoa*^fl/fl^;*Rhoc*^-/-^ and *Rhoa*^βMHC-cre^;*Rhoc*^-/-^ hearts. GAPDH was used as a loading control (n = 5 WT^βMHC-cre^, 5 *Rhoa*^fl/fl^, 4 *Rhoa*^βMHC-cre^, 4 *Rhoa*^fl/fl^;*Rhoc*^-/-^ and 7 *Rhoa*^βMHC-cre^;*Rhoc*^-/-^ hearts across panels B and C). (**D-K**) Heart weight-to-body weight (HW/BW), cardiomyocyte cross-sectional area (CSA), echocardiography measured fractional shortening (FS) percentage (%), and lung weight-to-body weight (LW/BW) in the indicated groups of mice at 8 weeks (**D-G**) and 8 months (**H-K**) of age. The legend shown refers to panels "D-K" in this figure. Data are shown as mean ± SD. Number of mice used in each experiment are shown in the graphs. For CSA, n = hearts, with ≥1,000 cardiomyocytes quantified per heart. Exact P values for significant comparisons are shown directly on the graphs by one-way ANOVA with Tukey’s multiple comparisons test; exact P values for all comparisons, including non-significant ones, are provided in Supplementary Table 1. (**L**) Low magnification cardiac histological images from indicated genotypes stained with Masson’s trichrome at 8 months of age. Images of entire heart sections were acquired using an Olympus BX51 microscope equipped with a Nikon DS-Fi3 camera and a 2× objective. White balance was set automatically on a tissue-free area of each slide adjacent to the section before acquisition; no further image adjustments were applied. Scale bar, 2 mm.

### Immunofluorescence

For cardiomyocyte cross-sectional area measurements, 5-µm paraffin heart sections were deparaffinized, washed 3 times with PBS and incubated with Alexa Fluor 647-conjugated wheat germ agglutinin (WGA; Molecular Probes, Eugene OR, #W32466; 1:100 in PBS) for 1 h at room temperature to label cardiomyocyte membranes. Sections were mounted using ProLong Diamond Antifade Mountant containing DAPI (Thermo Fisher Scientific, #P36971) to counterstain nuclei and preserve fluorescence.

Adult ventricular cardiomyocytes were isolated as described above and plated onto laminin-coated Ibidi µ-Slide 8-well chambers (Ibidi USA, Fitchburg WI, #80826). Cells were washed with PBS, fixed in 4% paraformaldehyde for 10 min, and rinsed three times with PBS. Following permeabilization and blocking with PBS containing 3% normal goat serum and 0.1% Triton X-100 for 20 min at room temperature or with 3% BSA in PBS, cells were incubated overnight at 4°C with either mouse monoclonal anti-α-actinin (Sigma-Aldrich, #A7811; 1:1000; RRID:AB_476766) or anti-Nav1.5 (Alomone Labs, Jerusalem, Israel, #ASC-005; 1:100; RRID: AB_2040001), respectively. After washing, cells were incubated for 45 min at room temperature with Alexa Fluor 568- or Alexa Fluor 488-conjugated secondary antibodies (Invitrogen; 1:400 in blocking buffer). Nuclei were counterstained with DAPI (Thermo Fisher Scientific, #D1306; 1:10,000).

Fluorescence imaging was performed under identical acquisition settings using a Nikon A1+ laser-scanning confocal microscope (Nikon Instruments, Melville, NY) equipped with a 40x water-immersion objective (NA 1.15) and NIS-Elements Advanced Research (AR) software (Nikon Instruments; RRID:SCR_014329). Representative images shown in the figures were selected from experiments with comparable results.

### Cardiomyocyte Cross-Sectional Area Measurements

Cardiomyocyte cross-sectional area (CSA) was quantified from WGA-stained heart sections using Nikon NIS-Elements software (version 5.42.06; Nikon Instruments; RRID:SCR_014329). WGA staining delineated the sarcolemma of individual cardiomyocytes and served as the basis for automated cell segmentation. Ground-truth binary masks were manually generated from representative heart sections and used to train the deep-learning-based 2D Segment Objects AI (OAI) algorithm. The initial model was trained for 500 iterations until a training loss of 0.01 was achieved and was subsequently retrained using an independent set of manually segmented images to further optimize segmentation accuracy while maintaining a training loss of 0.01. Following AI-based segmentation, a standardized image analysis workflow was applied that included: (i) identification of cardiomyocytes by OAI segmentation, (ii) removal of small objects using a clean filter (radius = 11 pixels; ∼3.57 µm), (iii) exclusion of elongated non-cardiomyocyte structures, such as blood vessels, using a circularity filter (0.3-1.0), (iv) filling of internal segmentation gaps, and (v) exclusion of objects intersecting the image border. Automated segmentations were visually inspected, and incorrectly segmented cells were manually corrected using the Binary Segmentation tool by separating fused cells or merging cells that had been inappropriately divided because of weak WGA staining. WGA-stained sections were imaged on the Nikon A1+ laser-scanning confocal microscope described above using a 20× objective and identical acquisition settings for all groups, and the same trained segmentation model and analysis pipeline were applied uniformly to all experimental groups. At least 1,000 cardiomyocytes were quantified per heart, and all analyses were performed in a blinded manner.

### Western Blotting

Hearts were excised, snap-frozen in liquid nitrogen, and stored at −80°C until analysis. Tissue was homogenized in ice-cold modified radioimmunoprecipitation assay (RIPA) buffer containing 1% Triton X-100, 1% sodium deoxycholate, 0.1% sodium dodecyl sulfate (SDS), 50 mM Tris-HCl (pH 7.4), and 150 mM NaCl, supplemented with Halt Protease and Phosphatase Inhibitor Cocktail (Thermo Fisher Scientific, #78442). Samples were sonicated using a VirSonic 60 sonicator (SP Scientific, Warminster, PA) at power setting 3 for three 10-s cycles. Lysates were clarified by centrifugation at 20,500 x g for 14 min at 4°C and stored at −80°C until use. Protein concentrations were measured using a Direct Detect infrared spectrometer (MilliporeSigma, Burlington, MA).

Protein lysates were combined with 5x Laemmli sample buffer and heated at 100°C for 10 min. Equal amounts of protein were separated by SDS-PAGE and transferred onto polyvinylidene fluoride membranes (PVDF; EMD Millipore, #IPFL00010). Membranes were incubated with the primary antibodies listed below, followed by the appropriate secondary antibodies. For alkaline phosphatase-based detection, membranes were incubated with IgG-AP-conjugated secondary antibodies (Santa Cruz Biotechnology, Dallas TX; 1:2,500), developed using ECF substrate (GE Healthcare, Buckinghamshire, United Kingdom, #RPN5785), and imaged with a Gel Doc XR+ system using Image Lab software (Bio-Rad Laboratories, Hercules, CA; RRID:SCR_014210). Alternatively, membranes were incubated with IRDye 800CW- or IRDye 680RD-conjugated secondary antibodies (LI-COR Biosciences, Lincoln, NE; 1:3,000) and imaged using an Odyssey CLx Imaging System (LI-COR Biosciences). Band intensities were quantified using LI-COR Image Studio software (RRID:SCR_015795).

Primary antibodies included connexin 43 (Cell Signaling Technology, Danvers, MA; #3512; 1:1,000; RRID:AB_2294590), FBXO44 (Sigma-Aldrich, #HPA003363; 1:1,000; RRID:AB_1078831), glyceraldehyde-3-phosphate dehydrogenase (GAPDH; Fitzgerald Industries International, Gardner, MA; #10R-G109A; 1:10,000; RRID:AB_1285808), Na+/K+-ATPase α1 (Developmental Studies Hybridoma Bank, Iowa City, IA, #a6F; 1:250; RRID: AB_528092), Na+/K+-ATPase α2 (EMD Millipore, #AB9094; 1:500; RRID: AB_347648), Na+/K+-ATPase β1 (Developmental Studies Hybridoma Bank, #24; 1:250; RRID: AB_528094), anti-Nav1.5 (Alomone Labs, #ASC-005; 1:500; RRID: AB_2040001), NCX1 (Swant, Brugdorf, Switzerland; Cat# R3F1; 1:1,000; RRID:AB_2716744), RhoA 67B9 (Cell Signaling Technology, #2117; 1:500; RRID:AB_10693922), RhoC D40E4 (Cell Signaling Technology, #3430; 1:1,000; RRID:AB_2179246), and YAP1 (Cell Signaling Technology, #14074; 1:1,000; RRID:AB_2650491). Uncropped immunoblots corresponding to the data presented in this study are provided as supporting information, together with the positions of the molecular weight standards.

### RhoA Activation Assay

RhoA activity was determined using a RhoA Activation Assay Kit (Cell Biolabs, San Diego, CA; #STA-403-A) according to the manufacturer’s instructions. Briefly, snap-frozen heart tissue was homogenized in ice-cold assay lysis buffer supplemented with protease inhibitors, and lysates were clarified by centrifugation at 14,000 x g for 10 min at 4°C. Protein concentrations were determined as described above, prior to the assay, and equal amounts of total protein were incubated with Rhotekin Rho-binding domain (RBD) agarose beads for 1 h at 4°C with gentle end-over-end mixing to pull down GTP-bound (active) Rho proteins. Following incubation, the beads were collected by centrifugation, washed three times with assay buffer, and bound proteins were eluted by boiling in reducing SDS-PAGE sample buffer. Active RhoA was then detected by SDS-PAGE and immunoblotting with a RhoA-specific antibody (RhoA 67B9; see above), as described above.

### Statistics and Reproducibility

Data are presented as mean ± standard deviation (SD). Statistical analyses were performed using GraphPad Prism 9 software (GraphPad Software, Boston, MA; RRID:SCR_002798). Details of the statistical tests used, sample sizes (n), and the definition of n for each experiment are provided in the corresponding figure legends. Comparisons between two groups were performed using an unpaired, two-tailed Student’s T-test. For comparisons involving more than two groups, statistical significance was determined by one-way analysis of variance (ANOVA) followed by Tukey’s multiple-comparisons test. Multiplicity-adjusted P values are reported for ANOVA analyses. A two-sided P value <0.05 was considered statistically significant. No animals or data points were excluded from the analyses.

## RESULTS

### Compensatory regulation of RhoA and RhoC in cardiomyocytes

RhoA has previously been implicated in cardiomyocyte hypertrophy and protective signaling in the myocardium (Sah *et al*., 1999; Brown *et al*., 2006; Lauriol *et al*., 2014; Miyamoto, 2024) whereas the role of RhoC in cardiomyocytes has remained largely undefined. Because loss of one Rho family member can induce compensatory upregulation of related isoforms (Wheeler & Ridley, 2004), we sought to define the individual and combined functions of RhoA and RhoC in the heart. To this end, we generated mice with cardiomyocyte-specific deletion of *Rhoa* on a global *Rhoc*-null background (*Rhoa*^βMHC-cre^;*Rhoc*^-/-^) by crossing *Rhoa^fl/fl^* and *βMHC*-cre transgenic mice on the *Rhoc*-null background (**Fig. 1A**). Immunoblot analysis confirmed efficient deletion of RhoA in *Rhoa*^βMHC-cre^ hearts and absence of RhoC in *Rhoc*^-/-^ and compound mutant hearts (**Fig. 1B-C**). Notably, loss of *Rhoa* gene in cardiomyocytes increased RhoC protein expression, whereas deletion of *Rhoc* led to a reciprocal increase in RhoA protein, indicating compensatory cross-regulation (**Fig. 1B-C**). By comparison, RhoB, the third closely related isoform, was previously reported to be undetectable in cardiomyocytes (Aoki *et al*., 1998).

We next assessed cardiac structure and function under basal conditions. At 8 weeks of age, deletion of *Rhoa*, *Rhoc*, or both did not significantly alter heart weight-to-body weight ratio (HW/BW, one-way ANOVA, P = 0.999), fractional shortening (FS%, P = 0.252), or lung weight-to-body weight ratio (LW/BW, P = 0.279) across genotypes (**Fig. 1D, F-G**). However, cardiomyocyte cross-sectional area (CSA) was reduced in *Rhoa*^fl/fl^;*Rhoc*^-/-^ hearts compared with WT^βMHC-cre^ controls (P < 0.001; **Fig. 1E**). This reduction was not observed in *Rhoa*^βMHC-cre^;*Rhoc*^-/-^ hearts, which did not differ from WT^βMHC-cre^ (P = 0.9997) but did differ from *Rhoa*^fl/fl^;*Rhoc*^-/-^ (P < 0.0001), consistent with compensatory and partially opposing functions of RhoA and RhoC in regulating basal cardiomyocyte size (**Fig. 1E**). By 8 months of age, *Rhoa*^βMHC-cre^;*Rhoc*^-/-^ mice developed a spontaneous increase in HW/BW, increased cardiomyocyte cross-sectional area, and elevated LW/BW compared with all other genotypes (all P < 0.05) (**Fig. 1H-I, K**), consistent with age-dependent hypertrophic remodeling and early pulmonary congestion. Importantly, FS% remained preserved at this stage (one-way ANOVA, P = 0.695) (**Fig. 1J**), and neither single-mutant nor double-deleted hearts exhibited overt fibrosis or gross histopathologic abnormalities by Masson’s trichrome staining (**Fig. 1L**). Exact P values for all comparisons shown in figure 1 are provided in Supplementary Table 1.

Together, these findings support the existence of a compensatory RhoA-RhoC signaling axis in cardiomyocytes and indicate that combined loss of both GTPases is sufficient to drive age-dependent maladaptive remodeling that precedes overt systolic dysfunction or fibrosis.

### RhoA and RhoC coordinately regulate cardiomyocyte geometry and intercalated disc organization

We next performed morphometric analysis of isolated adult cardiomyocytes from 8-week-old hearts, prior to the development of overt cardiac pathology. Cardiomyocyte length differed significantly across genotypes (one-way ANOVA, P = 0.00390) (**Fig. 2A-B**), with *Rhoa*^fl/fl^;*Rhoc*^-/-^ and *Rhoa*^βMHC-cre^;*Rhoc*^-/-^ cardiomyocytes being significantly longer than *Rhoa*^fl/fl^ controls (P = 0.0266 and P = 0.0030, respectively), and double-deleted cells also significantly longer than *Rhoa*^βMHC-cre^ alone (P = 0.0481). Length did not differ between *Rhoa*^fl/fl^;*Rhoc*^-/-^ and double-deleted cardiomyocytes (P = 0.379), suggesting that elongation was mostly driven by *Rhoc* deficiency. Cardiomyocyte width also differed significantly across genotypes (one-way ANOVA, P < 0.001) (**Fig. 2 A,C**); *Rhoa*^fl/fl^;*Rhoc*^-/-^ cardiomyocytes were significantly narrower than both *Rhoa*^fl/fl^ (P < 0.001) and *Rhoa*^βMHC-cre^ (P = 0.0041) controls, whereas width of double-deleted cardiomyocytes did not differ significantly from either *Rhoa*^fl/fl^ (P = 0.248) or *Rhoa*^βMHC-cre^ (P = 0.9994) but was significantly greater than in *Rhoa*^fl/fl^;*Rhoc*^-/-^ (P = 0.0047), suggesting that concurrent *Rhoa* deletion restored cell width in *Rhoc*-deficient cardiomyocytes. Consequently, the length-to-width ratio differed significantly across genotypes (one-way ANOVA, P < 0.0001; **Fig. 2D**). Both *Rhoa*^fl/fl^;*Rhoc*^-/-^ and *Rhoa*^βMHC-cre^;*Rhoc*^-/-^ cardiomyocytes exhibited significantly greater elongation than either control genotype (all P < 0.01), with no significant difference between the *Rhoc*-deficient and double-deleted cardiomyocytes (P = 0.482). Together, these findings indicate that RhoA and RhoC regulate distinct aspects of cardiomyocyte geometry. Loss of *Rhoc* elongated and narrowed cardiomyocytes, whereas *Rhoa* deletion alone had no detectable effect on either dimension. However, concurrent *Rhoa* deletion restored cell width toward control values without affecting elongation, suggesting that in the absence of *Rhoc*, *Rhoa* contributes to cardiomyocyte narrowing (exact P values for all comparisons are provided in Supplementary Table 2).

**Figure 2.**
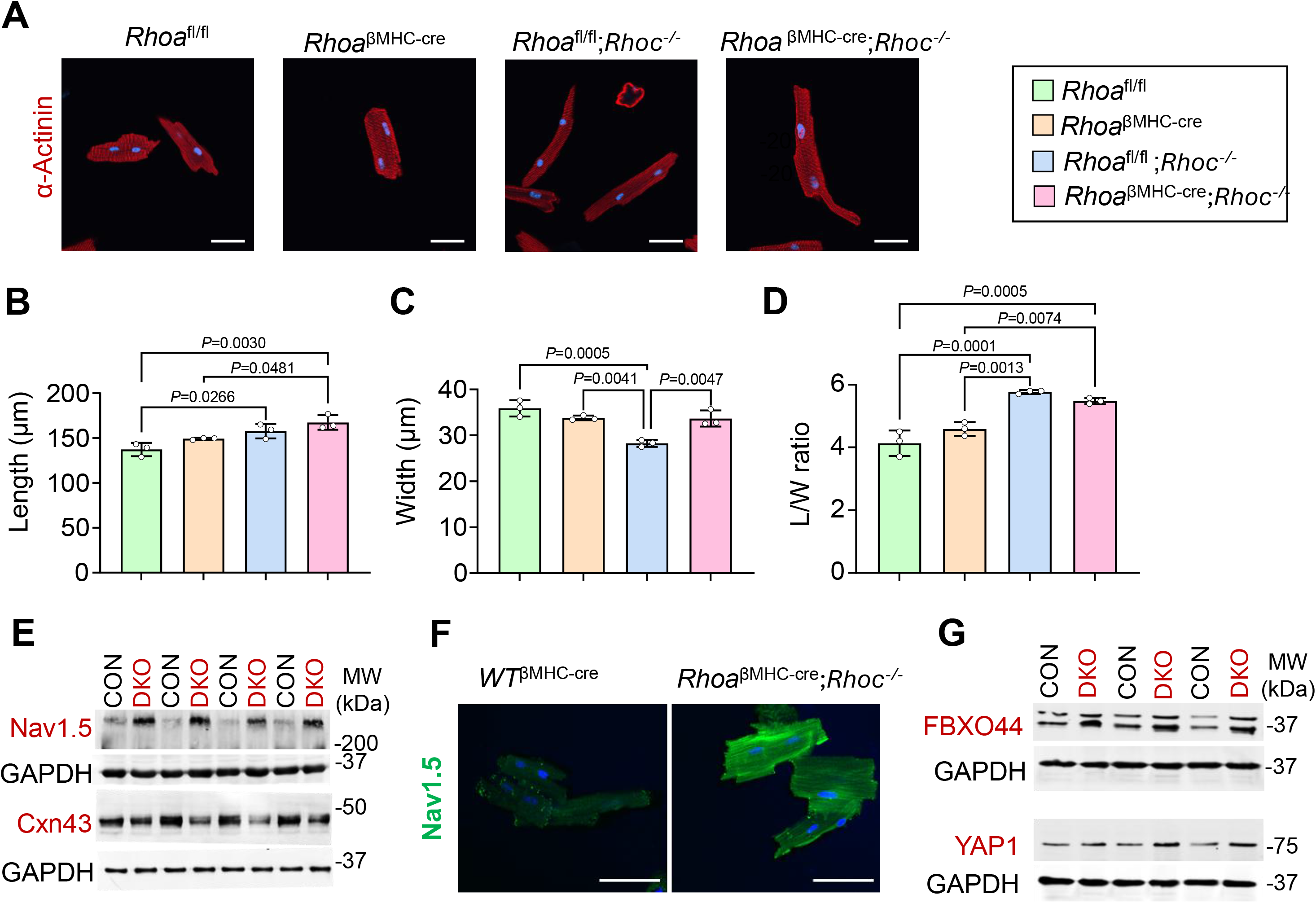
RhoA and RhoC coordinately regulate cardiomyocyte growth geometry and intercalated disc organization. (**A**) Representative images of isolated cardiomyocytes from the indicated genotypes at 8 weeks of age, stained for sarcomeric α-actinin (red), with nuclei counterstained with DAPI (blue). Images are single optical sections acquired on a Nikon A1+ confocal microscope with a 40× water-immersion objective; no post-acquisition processing was applied. Scale bar = 50 µm. (**B-D**) Quantitative analysis of isolated myocyte length (**B**), width (**C**), and length/width (L/W) ratio (**D**) from hearts of the indicated mice (n = 3 hearts per genotype, ≥150 cardiomyocytes measured per heart; data points represent per-heart means, which were used for statistical analysis). The legend shown refers to panels "B-D" in this figure. Data are shown as mean ± SD. Number of mice used in each experiment are shown in the graphs. Exact P values for significant comparisons are shown directly on the graphs by one-way ANOVA with Tukey’s multiple comparisons test. Exact P values for all comparisons, including non-significant ones, are provided in Supplementary Table 2. (**E**) Western blot analysis of Nav1.5 and Cxn43 protein levels in 8-week-old WT^βMHC-cre^ (CON) and *Rhoa*^βMHC-cre^;*Rhoc*^-/-^ hearts (DKO). GAPDH was used as a loading control (n = 4 hearts per genotype). (**F**) Representative images of isolated cardiomyocytes from WT^βMHC-cre^ and *Rhoa*^βMHC-cre^;*Rhoc*^-/-^ at 8 weeks of age, stained for Nav1.5 (green), with nuclei counterstained with DAPI (blue). Images are single optical sections acquired on a Nikon A1+ confocal microscope with a 40× water-immersion objective; no post-acquisition processing was applied. Scale bar = 30 µm. (**G**) Western blot analysis of FBXO44 and YAP1 protein levels in 8-week-old WT^βMHC-cre^ (CON) and *Rhoa*^βMHC-cre^;*Rhoc*^-/-^ hearts (DKO). GAPDH was used as a loading control (n = 3 hearts per genotype).

Because cardiomyocyte geometry is closely linked to cytoskeletal organization, membrane domain integrity, and mechanosensitive signaling (Nielsen *et al*., 2023), we next examined whether combined loss of *Rhoa* and *Rhoc* in cardiomyocytes alters proteins involved in intercalated disc organization and engages downstream remodeling pathways. Immunoblot analysis revealed a marked increase in total Nav1.5 protein in *Rhoa*^βMHC-cre^;*Rhoc*^-/-^ hearts, which is a key sodium channel at the cardiomyocyte sarcolemma and intercalated disc (P < 0.0001; **Fig. 2E, Supplemental Fig. 1A**). In contrast, protein levels of the Na+/K+-ATPase α1, α2, and β1 subunits, as well as NCX1, appeared unchanged, suggesting that this effect was not part of a broader alteration in major sodium-handling proteins (**Supplemental Fig. 1B**). Because Nav1.5 and connexin 43 (Cxn43) are structurally and functionally linked within the intercalated disc (Nielsen *et al*., 2023), we also examined Cxn43 and found that its abundance was reduced by approximately 50% in the *Rhoa*^βMHC-cre^;*Rhoc*^-/-^ hearts (P = 0.0045; **Fig. 2E, Supplemental Fig. 1C**). Interestingly, immunostaining for Nav1.5 in isolated *Rhoa*^βMHC-cre^;*Rhoc*^-/-^ cardiomyocytes showed extension of expression across membrane domains versus control myocytes (**Fig. 2F**).

Given these structural abnormalities, we also examined signaling pathways associated with mechanosensing and protein remodeling. *Rhoa*^βMHC-cre^;*Rhoc*^-/-^ hearts exhibited increased protein levels of YAP1 (Dupont *et al*., 2011), a central effector of mechanosensitive signaling, and FBXO44 (Sjogren *et al*., 2015; Liu *et al*., 2024), an F-box protein linked to ubiquitin-dependent protein turnover (P = 0.0361 for YAP1 and P = 0.0144 for FBXO44; **Fig. 2G, Supplemental Fig. 1D-E**). Collectively, these findings indicate that combined loss of RhoA and RhoC protein produces early abnormalities in cardiomyocyte geometry and expression of intercalated disc proteins before overt cardiac pathology becomes evident.

### RhoA and RhoC are induced by pressure overload and cooperatively restrain pathological remodeling

Given that RhoA activity is regulated by cycling between inactive GDP-bound and active GTP-bound states (Eckenstaler *et al*., 2022), we examined both total RhoA protein abundance and its active, GTP-bound fraction following transverse aortic constriction (TAC). Consistent with previous reports (Aoki *et al*., 1998; Lauriol *et al*., 2014), total and active GTP-bound RhoA levels were increased in *Rhoa*^fl/fl^ control hearts at 2, 7, and 14 days after TAC compared with sham controls (**Fig. 3A**). RhoC protein abundance was similarly elevated at 7 and 14 days, indicating that both GTPases are induced in response to pressure overload (**Fig. 3A**).

**Figure 3.**
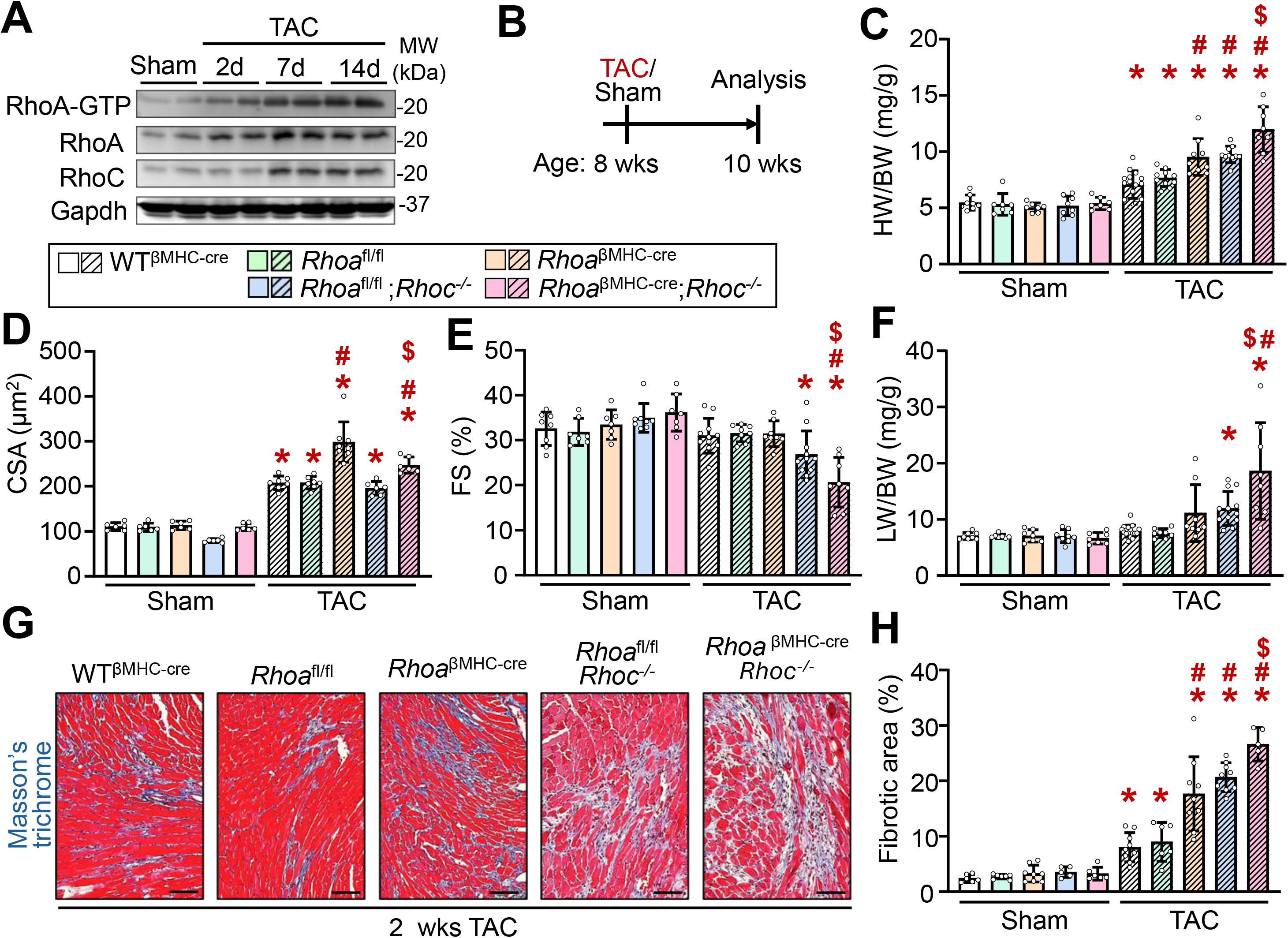
Deletion of *Rhoa* and *Rhoc* exacerbate pressure overload hypertrophy and failure. (**A**) Western blot for active RhoA (RhoA-GTP) and total levels of RhoA and RhoC from *Rhoa*^fl/fl^ hearts subjected to sham surgery or 2, 7, or 14 days (d) of transverse aortic constriction (TAC). GAPDH was used as a loading control (n = 2 hearts per time point). (**B**) Schematic diagram depicting the experimental outline. Eight-week-old mice were subjected to TAC or sham surgery and harvested 2 weeks later. (**C-F**) Quantification of HW/BW, cardiomyocyte cross-sectional area (CSA), echocardiography measured FS% and LW/BW in the indicated groups of mice subjected to 2 weeks of TAC or sham surgery (**G**) Representative histological images of cardiac fibrosis stained in blue by Masson’s trichrome in the indicated genotypes subjected to TAC for 2 weeks. Images were acquired using an Olympus BX51 microscope equipped with a Nikon DS-Fi3 camera and a 20× objective. White balance was set automatically on a tissue-free area of each slide adjacent to the section before acquisition; no further image adjustments were applied. Scale bars, 50 µm. (**H**) Quantitative analysis of cardiac fibrosis following 2 weeks of TAC compared with sham-operated controls across the indicated genotypes. The legend shown refers to all graphs in the figure. Data are shown as mean ± SD. Number of mice used in each experiment are shown in the graphs. For CSA, n = hearts shown in graphs, with ≥1,000 cardiomyocytes quantified per heart. *P < 0.05 vs. sham of the same genotype; #P < 0.05 vs. both WT^βMHC-cre^-TAC and *Rhoa*^fl/fl^-TAC; $P < 0.05 vs. both *Rhoa*^βMHC-cre^-TAC and *Rhoa*^fl/fl^;*Rhoc*^-/-^-TAC (one-way ANOVA with Tukey’s multiple comparisons test). Exact P values for all comparisons are provided in Supplementary Table 3.

To directly assess their functional roles, 8-week-old WT^βMHC-cre^, *Rhoa*^fl/fl^, *Rhoa*^βMHC-cre^, *Rhoa*^fl/fl^;*Rhoc*^-/-^, and *Rhoa*^βMHC-cre^;*Rhoc*^-/-^ mice were subjected to TAC and analyzed 2 weeks later (**Fig. 3B**). HW/BW increased significantly following TAC in all genotypes (all P < 0.05; **Fig. 3C**). The two TAC-treated control groups did not differ significantly (P = 0.957), whereas *Rhoa*^βMHC-cre^, *Rhoa*^fl/fl^;*Rhoc*^-/-^, and *Rhoa*^βMHC-cre^;*Rhoc*^-/-^ hearts exhibited significantly greater hypertrophy than both WT^βMHC-cre^ (all P < 0.001) and *Rhoa*^fl/fl^ (P = 0.0151, P = 0.00230, and P < 0.001, respectively) controls. Although the two single-mutant groups showed comparable hypertrophy (P > 0.999), the increase in HW/BW was significantly greater in *Rhoa*^βMHC-cre^;*Rhoc*^-/-^ hearts than in either *Rhoa*^βMHC-cre^ (P < 0.001) or *Rhoa*^fl/fl^;*Rhoc*^-/-^ (P = 0.00260) hearts. These findings indicate that combined loss of *Rhoa* and *Rhoc* exacerbates the hypertrophic response to pressure overload beyond either single deletion alone. Indeed, cardiomyocyte cross-sectional area also increased significantly following TAC in all genotypes (all P < 0.001; **Fig. 3D**), although the genotype-dependent differences in cardiomyocyte cross-sectional area did not mirror those observed in whole-heart hypertrophy. *Rhoa*^βMHC-cre^ hearts exhibited the greatest increase, significantly exceeding both control genotypes and the two other mutant genotypes (all P < 0.001). In contrast, cardiomyocyte cross-sectional area in *Rhoa*^fl/fl^;*Rhoc*^-/-^ hearts did not differ significantly from either WT^βMHC-cre^ (P = 0.978) or *Rhoa*^fl/fl^ (P = 0.976) controls, despite a significant increase relative to their own sham group (P < 0.001). *Rhoa*^βMHC-cre^;*Rhoc*^-/-^ hearts exhibited an intermediate response, with cross-sectional area greater than in both WT^βMHC-cre^ (P = 0.0130) and *Rhoa*^fl/fl^ (P = 0.0137) controls and *Rhoa*^fl/fl^;*Rhoc*^-/-^ hearts (P < 0.001), but lower than in *Rhoa*^βMHC-cre^ hearts (P < 0.001). Thus, the exaggerated increase in whole-heart mass following loss of *Rhoc* was not accompanied by a proportional increase in cardiomyocyte cross-sectional area, suggesting greater dilatory growth.

We next assessed whether these differences in hypertrophic remodeling were accompanied by changes in cardiac function. FS% remained preserved after 2 weeks of TAC in WT^βMHC-cre^, *Rhoa*^fl/fl^, and *Rhoa*^βMHC-cre^ hearts (all P > 0.99 versus respective sham controls; **Fig. 3E**). In contrast, FS% declined significantly in *Rhoa*^fl/fl^;*Rhoc*^-/-^ (P = 0.0010) and *Rhoa*^βMHC-cre^;*Rhoc*^-/-^ hearts (P < 0.0001). Although FS% in *Rhoa*^fl/fl^;*Rhoc*^-/-^ hearts did not differ significantly from either TAC-treated WT^βMHC-cre^ (P = 0.275) or *Rhoa*^fl/fl^ (P = 0.228) controls, *Rhoa*^βMHC-cre^;*Rhoc*^-/-^ hearts exhibited significantly greater functional impairment than both control genotypes (both P < 0.0001). Moreover, FS% in *Rhoa*^βMHC-cre^;*Rhoc*^-/-^ hearts was significantly lower than in either *Rhoa*^βMHC-cre^ (P < 0.0001) or *Rhoa*^fl/fl^;*Rhoc*^-/-^ hearts (P = 0.0346). These findings indicate that loss of *Rhoc* compromises systolic function early during pressure overload, with further deterioration following concurrent *Rhoa* deletion.

Consistent with the functional measurements, LW/BW remained unchanged after TAC in WT^βMHC-cre^, *Rhoa*^fl/fl^, and *Rhoa*^βMHC-cre^ mice (all P > 0.33 versus respective sham controls; **Fig. 3F**). *Rhoa*^fl/fl^;*Rhoc*^-/-^ mice exhibited a modest but significant increase relative to sham (P = 0.0438), although this did not reach significance compared with TAC-treated WT^βMHC-cre^ (P = 0.052) or *Rhoa*^fl/fl^ (P = 0.109) controls. In contrast, *Rhoa*^βMHC-cre^;*Rhoc*^-/-^ mice developed a marked increase in LW/BW (P < 0.0001 versus sham), significantly exceeding both control genotypes (both P < 0.0001), as well as *Rhoa*^βMHC-cre^ (P = 0.0007) and *Rhoa*^fl/fl^;*Rhoc*^-/-^ (P = 0.0010) mice following TAC. Together with the pronounced reduction in FS%, these findings indicate that combined loss of *Rhoa* and *Rhoc* accelerates the development of cardiac decompensation during pressure overload suggesting that these 2 proteins mostly overlap in function during cardiac stress.

Finally, cardiac fibrosis, quantified from Masson’s trichrome-stained sections, increased significantly following TAC in all genotypes (all P < 0.05 versus respective sham controls; **Fig. 3G–H**). Fibrosis was comparable between the two TAC-treated control groups (P = 0.9998), but was significantly greater in *Rhoa*^βMHC-cre^, *Rhoa*^fl/fl^;*Rhoc*^-/-^, and *Rhoa*^βMHC-cre^;*Rhoc*^-/-^ hearts compared with both controls (all P < 0.001). Although the two single-mutant groups did not differ significantly (P = 0.589), fibrosis was further increased in *Rhoa*^βMHC-cre^;*Rhoc*^-/-^ hearts compared with either *Rhoa*^βMHC-cre^ (P = 0.0002) or *Rhoa*^fl/fl^;*Rhoc*^-/-^ (P = 0.0384) hearts. Thus, combined loss of *Rhoa* and *Rhoc* exacerbates fibrotic remodeling beyond either single deletion alone. Exact P values for all comparisons shown in figure 3 are provided in Supplementary Table 3.

### Cardiomyocyte RhoA protects against progression to heart failure during sustained pressure overload

To confirm previous reports that cardiomyocyte-specific deletion of *Rhoa* predisposes to progressive cardiac dysfunction during chronic hemodynamic stress (Lauriol *et al*., 2014), we subjected WT^βMHC-cre^, *Rhoa*^fl/fl^, and *Rhoa*^βMHC-cre^ mice to 12 weeks of TAC (**Fig. 4A**). Chronic pressure overload significantly increased HW/BW in all three genotypes compared with their respective sham controls (WT^βMHC-cre^, P = 0.0167; *Rhoa*^fl/fl^ P = 0.0049; *Rhoa*^βMHC-cre^, P < 0.0001; **Fig. 4B**). While the two TAC-treated control groups did not differ (P = 0.983), hypertrophy was significantly greater in *Rhoa*^βMHC-cre^ hearts compared with both WT^βMHC-cre^ (P = 0.0034) and *Rhoa*^fl/fl^ (P = 0.0147) controls. Similarly, cardiomyocyte cross-sectional area increased significantly in all three genotypes following 12 weeks of TAC (all P < 0.0001), with a further increase in *Rhoa*^βMHC-cre^ hearts compared with WT^βMHC-cre^ (P = 0.0134) and *Rhoa*^fl/fl^ (P = 0.0022) controls (**Fig. 4C**). Similarly, FS% declined significantly following 12 weeks of TAC in all three genotypes (**Fig. 4D**) but the decline was significantly more pronounced in *Rhoa*^βMHC-cre^ mice, as was pulmonary edema (**Fig. 4E**). These findings indicate that loss of cardiomyocyte *Rhoa* accelerates the transition from compensated hypertrophy to decompensated heart failure during sustained pressure overload. However, fibrosis increased similarly in all three genotypes after 12 weeks of pressure overload (WT^βMHC-cre^, P = 0.0007; *Rhoa*^fl/fl^, P = 0.0005; *Rhoa*^βMHC-cre^, P = 0.0007), without significant differences between *Rhoa*^βMHC-cre^ hearts and either WT^βMHC-cre^ (P > 0.9999) or *Rhoa*^fl/fl^ (P = 0.9972) controls (**Fig. 4F, G**). Thus, the exaggerated functional deterioration observed in *Rhoa*-deficient hearts is unlikely to be explained by additional late-stage fibrotic remodeling (exact P values for all comparisons are provided in Supplementary Table 4), although RhoA functions to delay the transition from adaptive hypertrophy to heart failure, consistent with prior studies (Lauriol *et al*., 2014).

**Figure 4.**
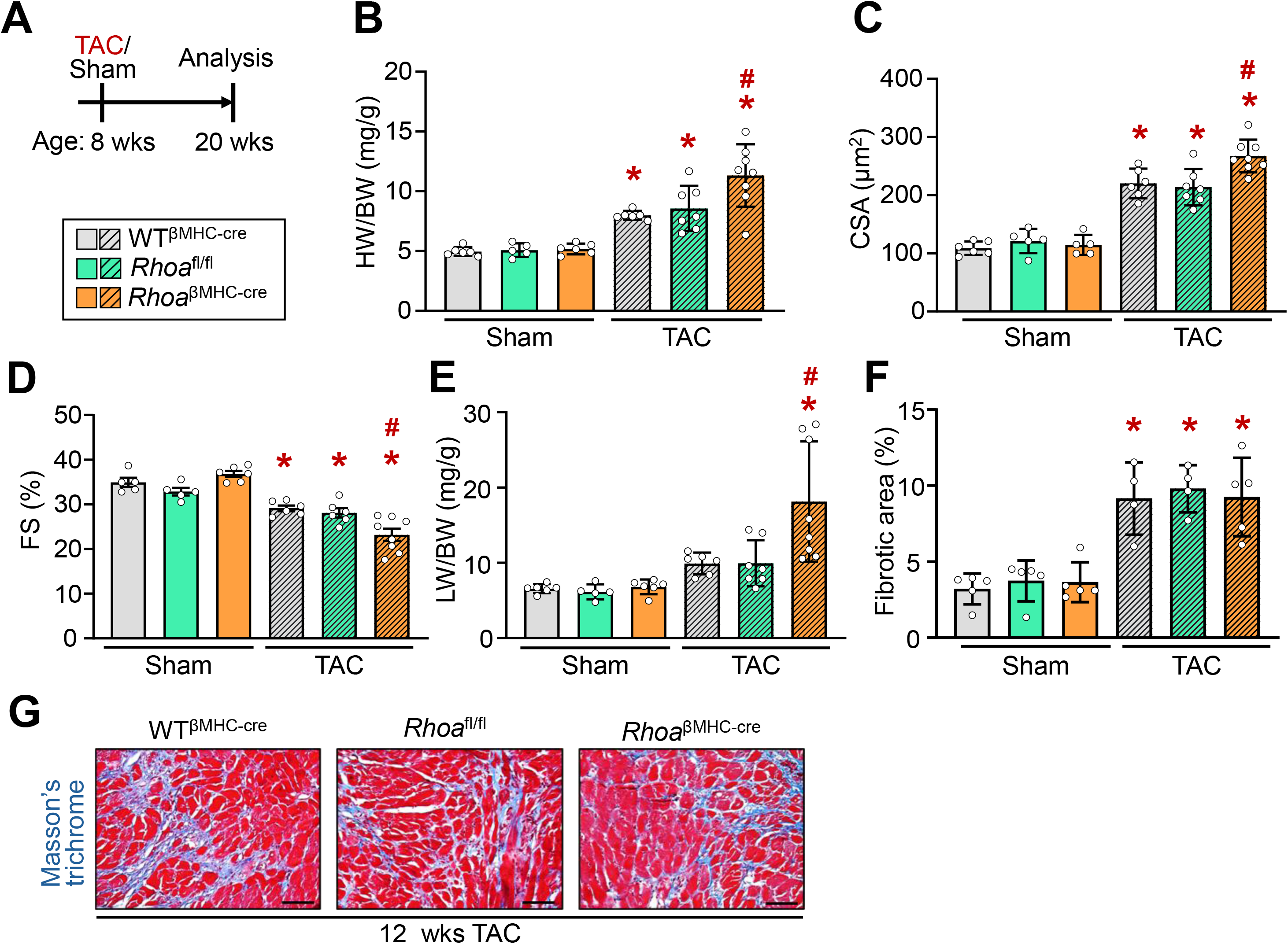
*Rhoa*^βMHC-cre^ hearts are more susceptible to failure after long-term pressure overload. (**A**) Schematic diagram depicting the experimental outline. Eight-week-old mice were subjected to TAC or sham surgery and harvested 12 weeks later. The legend shown refers to all other panels in the figure. (**B-E**) Quantification of HW/BW, cardiomyocyte CSA, echocardiography measured FS%, and LW/BW in the indicated groups of mice subjected to 12 weeks of TAC or sham surgery. (**F**) Quantitative analysis of cardiac fibrosis in hearts following 12 weeks of TAC compared with sham-operated controls across the indicated genotypes. (**G**) Representative histological images of cardiac fibrosis stained in blue by Masson’s trichrome in indicated genotypes subjected to TAC for 12 weeks. Images were acquired using an Olympus BX51 microscope equipped with a Nikon DS-Fi3 camera and a 20× objective. White balance was set automatically on a tissue-free area of each slide adjacent to the section before acquisition; no further image adjustments were applied. Scale bars, 50 µm. Data are shown as mean ± SD. Number of mice used in each experiment are shown in the graphs. For CSA, n = hearts shown by dots, with ≥1,000 cardiomyocytes quantified per heart. *P < 0.05 vs. sham of the same genotype; #P < 0.05 vs. both WT^βMHC-cre^-TAC and *Rhoa*^fl/fl^-TAC (one-way ANOVA with Tukey’s multiple comparisons test). Exact P values for all comparisons are provided in Supplementary Table 4.

### Loss of *Rhoa* and *Rhoc* exacerbates neurohumoral stress–induced cardiac dysfunction

Since Rho signaling is typically GPCR coupled (Aoki *et al*., 1998; Aikawa *et al*., 2000; Brown *et al*., 2006; Appert-Collin *et al*., 2007), we next examined the response to neurohumoral stimulation using combined angiotensin II and phenylephrine (Ang II/PE) infusion for 2 weeks (**Fig. 5A**). Ang II/PE infusion did not significantly increase HW/BW in WT^βMHC-cre^ (P = 0.189) or *Rhoa*^fl/fl^ (P = 0.303) hearts. In contrast, both *Rhoa*^βMHC-cre^ (P = 0.0036) and *Rhoa*^fl/fl^;*Rhoc*^-/-^ (P = 0.0001) hearts exhibited a significant increase relative to their respective vehicle-treated controls, although neither differed significantly from either control genotype following Ang II/PE infusion (all P > 0.06; **Fig. 5B**). Notably, *Rhoa*^βMHC-cre^;*Rhoc*^-/-^ hearts exhibited a markedly greater hypertrophic response (P < 0.0001 versus vehicle), significantly exceeding both control genotypes (both P < 0.0001) and both *Rhoa*^βMHC-cre^ (P < 0.0001) and *Rhoa*^fl/fl^;*Rhoc*^-/-^ (P < 0.0001) hearts following Ang II/PE infusion. These findings indicate that combined loss of *Rhoa* and *Rhoc* enhances the whole-heart hypertrophic response to neurohumoral stimulation. Cardiomyocyte cross-sectional area increased significantly following Ang II/PE infusion in all genotypes, including WT^βMHC-cre^ (P = 0.0183) and *Rhoa*^fl/fl^ (P = 0.0312) controls (**Fig. 5C**). Despite this increase, neither *Rhoa*^βMHC-cre^ nor *Rhoa*^fl/fl^;*Rhoc*^-/-^ hearts differed significantly from either control genotype following Ang II/PE infusion (all P > 0.9999). In contrast, *Rhoa*^βMHC-cre^;*Rhoc*^-/-^ hearts exhibited significantly greater cardiomyocyte cross-sectional area than both controls (both P < 0.0001), as well as *Rhoa*^βMHC-cre^ (P < 0.0001) and *Rhoa*^fl/fl^;*Rhoc*^-/-^ (P = 0.0001) hearts (**Fig. 5C**). Thus, combined loss of both GTPases exacerbates the increase in cardiomyocyte cross-sectional area in response to neurohumoral stress beyond either single deletion alone, similar to long-term TAC stimulation.

**Figure 5.**
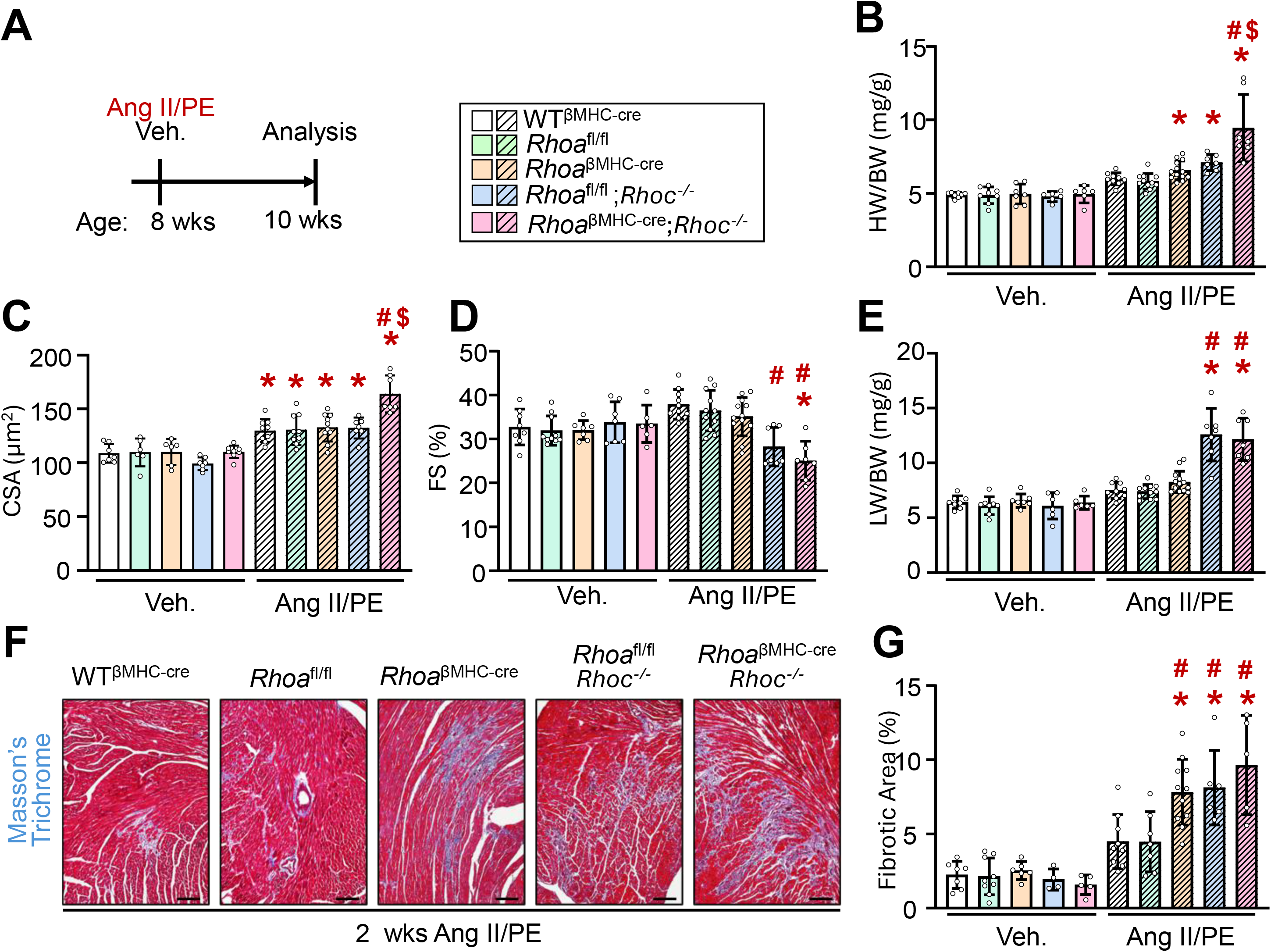
Deletion of *Rhoa* and *Rhoc* enhances Ang II/PE induced cardiac hypertrophy and dysfunction. (**A**) Schematic diagram depicting the experimental outline. Here, 8-week-old mice were subjected to 2 weeks of angiotensin II (Ang II)/phenylephrine (PE) stimulation or vehicle control before analysis. The legend shown refers to all other panels in the figure. (**B-E**) Quantification of HW/BW, cardiomyocyte CSA, echocardiography measured FS%, and LW/BW in the indicated groups of mice subjected to 2 weeks of Ang II/PE stimulation or vehicle control. (**F**) Representative histological images of cardiac fibrosis stained in blue by Masson’s trichrome in indicated genotypes subjected to 2 weeks of Ang II/PE stimulation. Images were acquired using an Olympus BX51 microscope equipped with a Nikon DS-Fi3 camera and a 20× objective. White balance was set automatically on a tissue-free area of each slide adjacent to the section before acquisition; no further image adjustments were applied. Scale bars, 50 µm. (**G**) Quantitative analysis of cardiac fibrosis following 2 weeks of Ang II/PE stimulation or vehicle control across the indicated genotypes. All data shown are mean ± SD. Number of mice used in each experiment are shown in the graphs. For CSA, n = hearts shown in the graphs, with ≥1,000 cardiomyocytes quantified per heart. *P < 0.05 vs. vehicle of the same genotype; #P < 0.05 vs. both WT^βMHC-cre^-Ang II/PE and *Rhoa*^fl/fl^-Ang II/PE; $P < 0.05 vs. both *Rhoa*^βMHC-cre^-Ang II/PE and *Rhoa*^fl/fl^;*Rhoc*^-/-^-Ang II/PE (one-way ANOVA with Tukey’s multiple comparisons test). Exact P values for all comparisons are provided in Supplementary Table 5.

We next assessed whether these differences in hypertrophic remodeling were accompanied by changes in cardiac function. FS% did not decline significantly following Ang II/PE infusion in WT^βMHC-cre^ (P = 0.190), *Rhoa*^fl/fl^ (P = 0.248), *Rhoa*^βMHC-cre^ (P = 0.842), or *Rhoa*^fl/fl^;*Rhoc*^-/-^ (P = 0.260) hearts versus respective vehicle-treated controls (**Fig. 5D**). However, *Rhoa*^fl/fl^;*Rhoc*^-/-^ hearts exhibited significantly lower FS% than both WT^βMHC-cre^ (P = 0.0003) and *Rhoa*^fl/fl^ (P = 0.0029) controls, as well as *Rhoa*^βMHC-cre^ hearts (P = 0.0226), with Ang II/PE infusion. *Rhoa*^βMHC-cre^;*Rhoc*^-/-^ hearts exhibited a significant decline in FS% relative to vehicle-treated mice (P = 0.0125), and significantly lower fractional shortening than both control genotypes (both P < 0.0001) and *Rhoa*^βMHC-cre^ hearts (P < 0.0001). Notably, FS% did not differ significantly between *Rhoa*^fl/fl^;*Rhoc*^-/-^ and *Rhoa*^βMHC-cre^;*Rhoc*^-/-^ hearts (P = 0.891; **Fig. 5D**). These findings suggest that RhoC plays a particularly important role in preserving systolic function during neurohumoral stress, with concurrent deletion of *Rhoa* producing no detectable additional functional impairment under these experimental conditions. Consistent with these functional findings, LW/BW increased significantly following Ang II/PE infusion in *Rhoa*^fl/fl^;*Rhoc*^-/-^ and *Rhoa*^βMHC-cre^;*Rhoc*^-/-^ mice (both P < 0.0001 versus respective vehicle-treated controls), but remained unchanged in WT^βMHC-cre^ (P = 0.655), *Rhoa*^fl/fl^ (P = 0.331), and *Rhoa*^βMHC-cre^ (P = 0.0624) mice (**Fig. 5E**). LW/BW did not differ significantly between *Rhoa*^fl/fl^;*Rhoc*^-/-^ and *Rhoa*^βMHC-cre^;*Rhoc*^-/-^ mice following Ang II/PE infusion (P = 0.9994), although it was significantly greater in *Rhoa*^βMHC-cre^;*Rhoc*^-/-^ mice than in *Rhoa*^βMHC-cre^ mice (P < 0.0001). Together, these observations further support an important role for RhoC in limiting functional deterioration and pulmonary congestion during neurohumoral stress.

Finally, cardiac fibrosis increased significantly following Ang II/PE infusion in *Rhoa*^βMHC-cre^ (P < 0.0001), *Rhoa*^fl/fl^;*Rhoc*^-/-^ (P = 0.0001), and *Rhoa*^βMHC-cre^;*Rhoc*^-/-^ hearts (P < 0.0001), but not in WT^βMHC-cre^ (P = 0.330) or *Rhoa*^fl/fl^ (P = 0.282) controls (**Fig. 5F, G**). All three mutant genotypes exhibited significantly greater fibrosis than both control genotypes (all P < 0.03), without significant differences between the mutant genotypes themselves (all pairwise P > 0.70). Thus, deletion of either *Rhoa* or *Rhoc* in cardiomyocytes is sufficient to exacerbate fibrotic remodeling during neurohumoral stress, with no additional effect with combined deletion. Exact P values for all comparisons are provided in Supplementary Table 5.

Collectively, these findings demonstrate that RhoA and RhoC cooperatively limit hypertrophic remodeling in response to neurohumoral stress, with combined loss of both GTPases producing the greatest increase in heart mass and cardiomyocyte cross-sectional area. In contrast, the functional deterioration and pulmonary congestion observed in absence of *Rhoc* were not further exacerbated by concurrent *Rhoa* deletion, highlighting a particularly important role for RhoC in preserving cardiac function during neurohumoral stress.

## DISCUSSION

In this study, we demonstrate that RhoA and RhoC undergo reciprocal compensatory regulation in the myocardium but make distinct, non-redundant contributions to cardiomyocyte geometry and the cardiac response to stress. Combined loss of *Rhoa* and *Rhoc* produced age-dependent hypertrophic remodeling under baseline conditions, exacerbated the hypertrophic response to both hemodynamic and neurohumoral stress beyond either single deletion, and further increased fibrosis during pressure overload. However, their relative contributions to hypertrophic growth and systolic function varied with the nature and duration of the stimulus, indicating that these closely related GTPases are not fully interchangeable regulators of cardiac remodeling.

The reciprocal increase in RhoA and RhoC following deletion of the other isoform is consistent with a broader pattern of cross-regulation described among Rho family members in other tissues (Wheeler & Ridley, 2004; Zandvakili *et al*., 2015). A plausible mechanistic basis for this compensation is competition for RhoGDIα, a chaperone that stabilizes Rho GTPases. In cancer cells, RhoA and RhoC compete for a shared, limiting pool of RhoGDIα, such that reducing one isoform frees RhoGDIα to stabilize the other and increase its steady-state protein abundance (Giang Ho *et al*., 2011). However, this compensatory regulation was insufficient to preserve normal cardiomyocyte geometry, since *Rhoc* deficiency produced elongated and narrowed cardiomyocytes, whereas concurrent *Rhoa* deletion restored cell width toward control values without correcting elongation. Such divergent effects may reflect isoform-specific effector coupling. In other cell types, RhoC signals preferentially through formin-like 2 and 3 (FMNL2 and FMNL3), whereas RhoA signals prominently through ROCK (Vega *et al*., 2011). FMNL family members are expressed in the postnatal heart, and FMNL2 contributes to myofibril organization in cardiomyocytes (Rosado *et al*., 2014). Cardiomyocyte lengthening and widening reflect the preferential addition of sarcomeres in series or in parallel, respectively (Russell *et al*., 2010), and the balance between them is influenced by mitogen-activated protein kinase kinase 1 (MEK1)-ERK1/2 signaling. Activated MEK1 promotes widening and ERK1/2 deletion promotes lengthening, potentially through remodeling of the non-sarcomeric actin cytoskeleton (Kehat *et al*., 2011; Grimes *et al*., 2024). Because cardiomyocyte *Rhoa* deletion reduces ERK1/2 activation during pressure overload (Lauriol *et al*., 2014), Rho GTPase signaling may intersect with this pathway. Previous work also showed that manipulation of RhoA activity did not alter cardiomyocyte length-to-width ratio *in vitro*, in contrast to the small GTPase cell division cycle 42 (Cdc42) (Nagai *et al*., 2003). Together, these observations suggest that RhoC contributes to aspects of cardiomyocyte geometry that are not fully compensated by increased RhoA, although the underlying signaling mechanisms remain to be defined.

Combined *Rhoa*/*Rhoc* deficiency also altered proteins involved in intercalated disc organization and mechanosensitive signaling before the development of overt cardiac pathology, including altered Nav1.5 localization, reduced Cxn43 abundance, and increased YAP1 and FBXO44. Given the dependence of intercalated disc organization on the actin cytoskeleton (Noorman *et al*., 2009; Pruna & Ehler, 2020; Nielsen *et al*., 2023), these changes suggest that RhoA and RhoC may contribute to the maintenance of cardiomyocyte structural integrity and mechanical coupling. Indeed, RhoA-dependent actin organization has been implicated in Cxn43 gap junction regulation in a heterologous expression system (Jara *et al*., 2024), and Nav1.5 membrane retention depends in part on the cortical actin cytoskeleton (Ziane *et al*., 2010), whereas YAP1 is sensitive to cytoskeletal tension and contributes to cardiac hypertrophic signaling (Dupont *et al*., 2011; Byun *et al*., 2019). Increased FBXO44 may also influence the hypertrophic response through degradation of RGS2, a negative regulator of Gq-coupled receptor signaling (Takimoto *et al*., 2009; Sjogren *et al*., 2015). Although these pathways provide potential links between *Rhoa*/*Rhoc* deficiency and the observed structural and hypertrophic abnormalities, their causal involvement remains to be established.

The distinct contributions of RhoA and RhoC became particularly apparent during TAC-induced pressure overload. FS% declined relative to sham within two weeks in *Rhoc*-deficient hearts, whereas *Rhoa*-deficient hearts remained functionally compensated despite displaying the greatest increase in cardiomyocyte cross-sectional area. This early hypertrophic response contrasts with a previous report in which cardiomyocyte-specific *Rhoa* deletion produced hypertrophy comparable to control hearts after acute pressure overload (Lauriol *et al*., 2014). That prior model used a different Cre driver than we employed (Oka *et al*., 2006). Nevertheless, prolonged pressure overload accelerated the transition to decompensated heart failure following *Rhoa* deletion in our model, consistent with the previously reported protective role of RhoA during chronic hemodynamic stress (Lauriol *et al*., 2014). A broadly similar pattern has been described for the related small GTPase Cdc42, whose cardiomyocyte-specific deletion also accelerates the transition to heart failure during pressure overload, albeit through a distinct, JNK-dependent signaling pathway (Maillet *et al*., 2009). Interestingly, IQGAP1, a shared interaction partner of RhoA and RhoC, supports adaptive cardiac remodeling through sustained ERK1/2 and AKT signaling during pressure overload (Sbroggio *et al*., 2011; Casteel *et al*., 2012). Whether RhoA, RhoC, or both contribute to this sustained adaptive response through IQGAP1 remains to be determined. Nevertheless, our findings suggest distinct temporal requirements for the two isoforms, with *Rhoc* deficiency associated with early functional vulnerability and the *Rhoa* gene becoming particularly important for preserving cardiac function during prolonged pressure overload.

An important finding across both stress models was the dissociation between hypertrophic growth and functional impairment. Following TAC, FS% declined relative to sham in *Rhoc*-deficient but not *Rhoa*-deficient hearts, despite smaller cardiomyocyte cross-sectional area in *Rhoc*-deficient hearts, although FS% did not differ significantly between the two genotypes after TAC. Similarly, under neurohumoral stimulation, FS% in *Rhoc*-deficient hearts was lower than in Ang II/PE-treated control and *Rhoa*-deficient hearts, although it did not decline significantly relative to vehicle-treated *Rhoc*-deficient mice, and *Rhoc* deficiency was sufficient to promote pulmonary congestion, whereas concurrent *Rhoa* deletion further exacerbated hypertrophic growth without additional functional deterioration. Thus, the magnitude of hypertrophy does not fully account for the functional abnormalities observed following *Rhoc* deficiency. Notably, *Rhoc*-deficient cardiomyocytes already exhibited an eccentric geometry (elongated and narrowed) at baseline, indicating that pre-existing geometric abnormalities may contribute to their subsequent functional vulnerability. Nevertheless, these findings suggest that RhoA and RhoC differentially regulate hypertrophic growth and the preservation of cardiac function.

Our findings also have implications for therapeutic strategies targeting RhoA signaling in heart failure. Recent evidence suggests that pharmacological inhibition of RhoA, including through RhoGDIα-dependent stabilization of its inactive GDP-bound state, may have therapeutic potential to treat heart failure (Xue *et al*., 2026). However, given the reciprocal regulation of RhoA and RhoC observed here, different approaches to RhoA inhibition may have distinct consequences for compensatory RhoC abundance and activity. Determining how pharmacological RhoA inhibition affects RhoC regulation may therefore be important for understanding its consequences in the diseased heart.

Several limitations warrant consideration. *Rhoc* deficiency was germline based and results in all cells lacking this protein, whereas *Rhoa* deletion was restricted to cardiomyocytes, and contributions from *Rhoc* loss in non-cardiomyocyte populations cannot be excluded. Furthermore, the proposed mechanistic relationships between *Rhoa*/*Rhoc* deficiency, cytoskeletal organization, and downstream signaling remain correlative. In addition, although male and female mice were included, the study was not powered to detect sex-specific effects. Finally, genetic deletion early in life may not fully recapitulate the consequences of acute pharmacological modulation in the adult heart. Future studies using cardiomyocyte-restricted *Rhoc* deletion and direct assessment of isoform-specific signaling will be needed to establish the mechanisms underlying these phenotypes.

In summary, RhoA and RhoC constitute a compensatory but functionally divergent signaling modules in the myocardium. Their distinct contributions to cardiomyocyte geometry, hypertrophic growth, and contractile function highlight the importance of isoform-specific regulation in the cardiac response to hemodynamic and neurohumoral stress.

## ADDITIONAL INFORMATION

### Data Availability Statement

Data supporting the findings of this study are included within the manuscript figures, with uncropped immunoblot images provided as Supporting Information. All other raw experimental data are available from the corresponding author upon reasonable request.

### Competing Interests

The authors declare no competing interests.

### Author Contributions

All experiments were performed in the laboratories of J.D.M. and D.V. at Cincinnati Children’s Hospital Medical Center. Conceptualization: V.P., J.D.M., D.V. Methodology: V.P., N.S.S.K., M.T., J.H.V.B., Y.Z., J.M.K., D.V. Investigation: V.P., N.S.S.K., M.T., J.H.V.B., D.V. Formal analysis: V.P., M.T., J.D.M., D.V. Resources: Y.Z. Data curation: V.P., N.S.S.K., M.T., D.V. Software: J.M.K. Visualization: V.P., J.D.M., D.V. Supervision: J.D.M., D.V. Writing – original draft: V.P., J.D.M., D.V. Writing – review & editing: all authors. All authors approved the final version of this manuscript and agree to be accountable for all aspects of the work in ensuring that questions related to the accuracy or integrity of any part of the work are appropriately investigated and resolved. All people designated as authors qualify for authorship, and all those who qualify for authorship are listed.

### Funding

This work was supported by a grant from the National Heart, Lung, and Blood Institute of the NIH [R01 HL105924 and R01 HL142217] to J.D.M., as well as by a grant by the FWO-Research Foundation Flanders [1208910N and V4.332.11N] to D.V.

### Generative AI Statement

During the preparation of this manuscript, the authors used Claude (Anthropic; Claude Opus 5.5, accessed via claude.ai) to improve grammar, language, and readability. The authors reviewed and edited the AI-assisted text and take full responsibility for the final content of the manuscript. No generative AI tools were used to generate, analyze or modify experimental data or images. Separately, cardiomyocyte cross-sectional area was quantified with a non-generative deep-learning segmentation tool (Segment Objects AI, Nikon NIS-Elements), as described in the Methods.

## Acknowledgements

None.

## AUTHORS’ TRANSLATIONAL PERSPECTIVE

Cardiac hypertrophy, the enlargement of heart muscle cells in response to increased workload or hormonal stress, can progress to heart failure when this adaptation fails. The signaling protein RhoA has been proposed as a target for new heart failure treatments, and recent evidence suggests that drugs inhibiting RhoA may offer therapeutic benefit. However, we show in mice that RhoA does not act alone in the heart. A closely related protein, RhoC, whose role in heart muscle remains poorly understood, works alongside RhoA to control the size and shape of heart muscle cells and to protect the heart during stress. Removing either RhoA or RhoC individually caused only a modest, and in some cases delayed, decline in heart function. Removing both together, however, produced substantially greater heart muscle enlargement and dysfunction than removing either alone, indicating that RhoA and RhoC compensate for one another. We also found that removing RhoC, but not RhoA, reduced heart function soon after a sudden increase in workload or during hormonal stress, whereas RhoA became more important during prolonged periods of increased workload. These findings indicate that RhoA and RhoC play distinct but complementary roles in protecting the heart, and that any future therapy targeting RhoA should consider this compensatory relationship with RhoC. Drugs that inhibit RhoA without addressing RhoC compensation could have unanticipated effects on heart muscle structure and function, and further research is needed to determine how RhoA-targeted treatments affect RhoC activity in the diseased human heart.

**Supplemental Figure 1. Selective alterations in ion-handling, gap junction, and growth-signaling proteins in pre-pathogenic *Rhoa*^βMHC-cre^;*Rhoc*^-/-^ hearts.** (**A**) Quantitation of Western blots for Nav1.5 shown in Fig. 2E. The legend shown refers to all other panels in the figure. (**B**) Western blot analysis of Na/K-ATPase α1, Na/K-ATPase α2, Na/K-ATPase β1, and NCX protein levels in 8-week-old WT^βMHC-cre^ (CON; N=4 hearts) and *Rhoa*^βMHC-cre^;*Rhoc*^-/-^ hearts (DKO; N=4 hearts). GAPDH was used as a loading control. (**C**) Quantitation of Western blots for Cxn43 shown in Fig. 2E. (**D-E**) Quantitation of Western blots for FBXO44 and YAP1 shown in Fig. 2G. All data shown are mean ± SD. Number of mice used in each experiment are shown in the graphs. Exact P values for significant comparisons are shown directly on the graphs by unpaired, two-tailed Student’s t-test.

## Notes

### Competing Interest Statement

The authors have declared no competing interest.

