## Supplementary material for "Small GTPases RhoA and RhoC coordinately regulate cardiomyocyte size in the mouse heart": Data Supplements

### SUPPLEMENTARY FIGURE 1

### SUPPLEMENTARY TABLES 1-5

**Genotype key used in Supplementary Tables 1-5:**

WT$^{\beta MHC\text{-}cre}$ (Cre-only control);

*Rhoa*$^{fl/fl}$ (*Rhoa* floxed control);

*Rhoa*$^{\beta MHC\text{-}cre}$ (cardiomyocyte-specific *Rhoa* deletion);

*Rhoa*$^{fl/fl}$;*Rhoc*$^{-/-}$ (global *Rhoc* deletion);

*Rhoa*$^{\beta MHC\text{-}cre}$;*Rhoc*$^{-/-}$ (combined *Rhoa/Rhoc* deletion).

Sham/Veh = Sham-operated or vehicle-treated controls;

TAC = transverse aortic constriction;

Ang/PE = angiotensin II/phenylephrine.

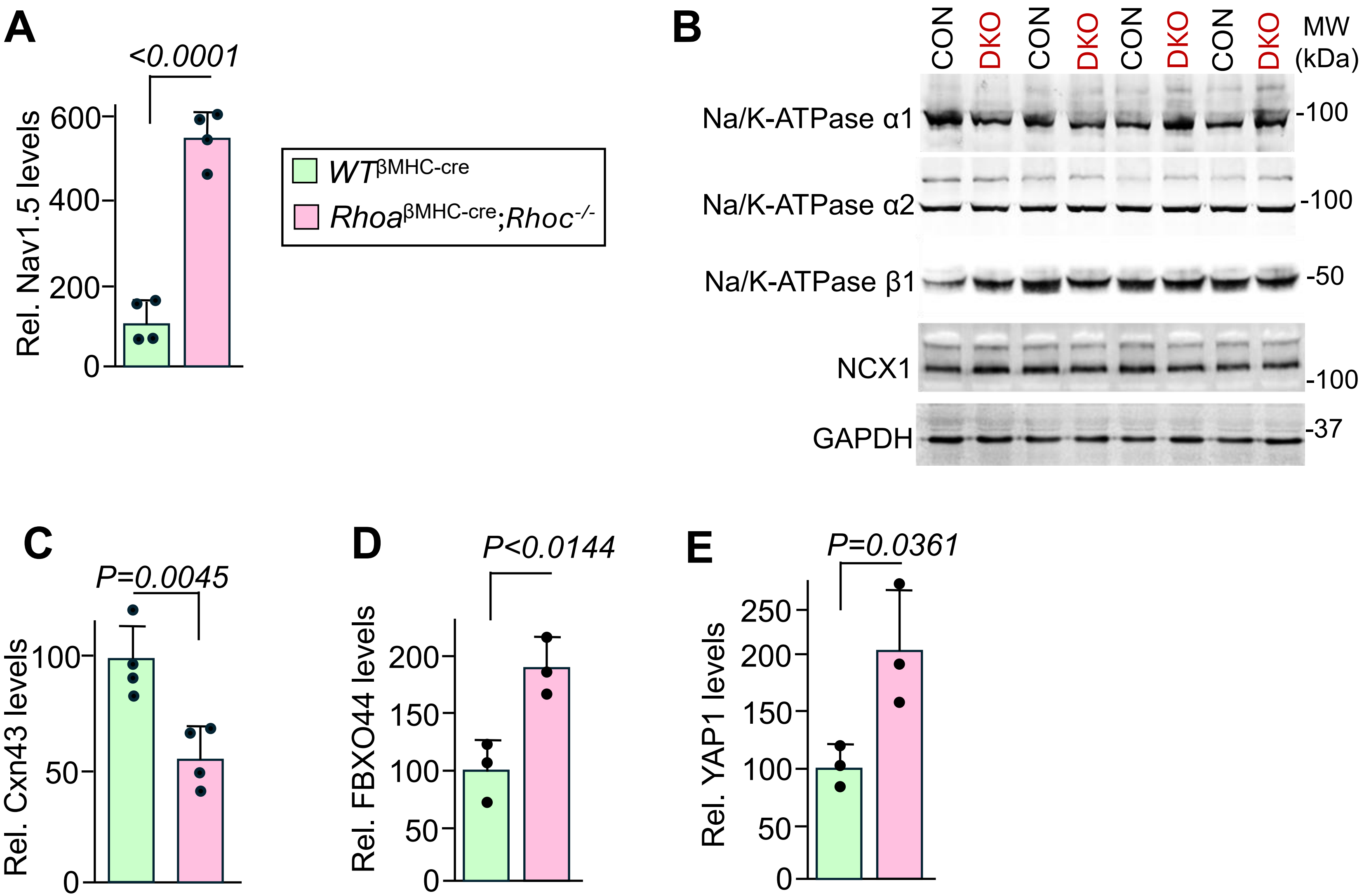


**Supplemental Figure 1. Selective alterations in ion-handling, gap junction, and growth-signaling proteins in pre-pathogenic *Rhoa*$^{\beta MHC\text{-}cre}$;*Rhoc*$^{-/-}$ hearts.** (**A**) Quantitation of Western blots for Nav1.5 shown in Fig. 2E. The legend shown refers to all other panels in the figure. (**B**) Western blot analysis of Na/K-ATPase α1, Na/K-ATPase α2, Na/K-ATPase β1, and NCX protein levels in 8-week-old WT$^{\beta MHC\text{-}cre}$ (CON; N=4 hearts) and *Rhoa*$^{\beta MHC\text{-}cre}$;*Rhoc*$^{-/-}$ hearts (DKO; N=4 hearts). GAPDH was used as a loading control. (**C**) Quantitation of Western blots for Cxn43 shown in Fig. 2E. (**D-E**) Quantitation of Western blots for FBXO44 and YAP1 shown in Fig. 2G. All data shown are mean ± SD. Number of mice used in each experiment are shown in the graphs. Exact P values for significant comparisons are shown directly on the graphs by unpaired, two-tailed Student's t-test.

**Supplementary Table 1. Statistical comparisons for Figure 1.**

| Comparison | Test | Mean Diff. | 95% CI | Adjusted P |
|---|---|---|---|---|
| **Figure 1, Panel D — HW/BW (8 wks)** | | | | |
| **Comparison** | **Test** | **Mean Diff.** | **95% CI** | **Adjusted P** |
| Omnibus one-way ANOVA | One-way ANOVA | F(4,40)=0.0276 | — | 0.999 |
| WT$^{\beta MHC\text{-}cre}$ vs *Rhoa*$^{fl/fl}$ | Tukey one-way ANOVA | 0.03492 | -0.8127 to 0.8825 | >0.9999 |
| WT$^{\beta MHC\text{-}cre}$ vs *Rhoa*$^{\beta MHC\text{-}cre}$ | Tukey one-way ANOVA | 0.01297 | -0.8346 to 0.8606 | >0.9999 |
| WT$^{\beta MHC\text{-}cre}$ vs *Rhoa*$^{fl/fl}$;*Rhoc*$^{-/-}$ | Tukey one-way ANOVA | -0.004743 | -0.8524 to 0.8429 | >0.9999 |
| WT$^{\beta MHC\text{-}cre}$ vs *Rhoa*$^{\beta MHC\text{-}cre}$;*Rhoc*$^{-/-}$ | Tukey one-way ANOVA | -0.06379 | -0.8827 to 0.7551 | 0.9994 |
| *Rhoa*$^{fl/fl}$ vs *Rhoa*$^{\beta MHC\text{-}cre}$ | Tukey one-way ANOVA | -0.02195 | -0.9505 to 0.9066 | >0.9999 |
| *Rhoa*$^{fl/fl}$ vs *Rhoa*$^{fl/fl}$;*Rhoc*$^{-/-}$ | Tukey one-way ANOVA | -0.03967 | -0.9682 to 0.8888 | >0.9999 |
| *Rhoa*$^{fl/fl}$ vs *Rhoa*$^{\beta MHC\text{-}cre}$;*Rhoc*$^{-/-}$ | Tukey one-way ANOVA | -0.09871 | -1.001 to 0.8036 | 0.9978 |
| *Rhoa*$^{\beta MHC\text{-}cre}$ vs *Rhoa*$^{fl/fl}$;*Rhoc*$^{-/-}$ | Tukey one-way ANOVA | -0.01771 | -0.9462 to 0.9108 | >0.9999 |
| *Rhoa*$^{\beta MHC\text{-}cre}$ vs *Rhoa*$^{\beta MHC\text{-}cre}$;*Rhoc*$^{-/-}$ | Tukey one-way ANOVA | -0.07676 | -0.9791 to 0.8256 | 0.9992 |
| *Rhoa*$^{fl/fl}$;*Rhoc*$^{-/-}$ vs *Rhoa*$^{\beta MHC\text{-}cre}$;*Rhoc*$^{-/-}$ | Tukey one-way ANOVA | -0.05904 | -0.9614 to 0.8433 | 0.9997 |
| **Figure 1, Panel E — CSA (8 wks)** | | | | |
| **Comparison** | **Test** | **Mean Diff.** | **95% CI** | **Adjusted P** |
| Omnibus one-way ANOVA | One-way ANOVA | F(4,25)=18.53 | — | **<0.0001** |
| WT$^{\beta MHC\text{-}cre}$ vs *Rhoa*$^{fl/fl}$ | Tukey one-way ANOVA | 1.364 | -12.33 to 15.05 | 0.9983 |
| WT$^{\beta MHC\text{-}cre}$ vs *Rhoa*$^{\beta MHC\text{-}cre}$ | Tukey one-way ANOVA | -2.811 | -16.50 to 10.88 | 0.9733 |

| | | | | |
|---|---|---|---|---|
| $WT^{\beta MHC\text{-}cre}$ vs $Rhoa^{fl/fl};Rhoc^{-/-}$ | Tukey one-way ANOVA | 31.38 | 17.69 to 45.07 | **<0.0001** |
| $WT^{\beta MHC\text{-}cre}$ vs $Rhoa^{\beta MHC\text{-}cre};Rhoc^{-/-}$ | Tukey one-way ANOVA | 0.8719 | -12.82 to 14.56 | 0.9997 |
| $Rhoa^{fl/fl}$ vs $Rhoa^{\beta MHC\text{-}cre}$ | Tukey one-way ANOVA | -4.175 | -17.87 to 9.517 | 0.8959 |
| $Rhoa^{fl/fl}$ vs $Rhoa^{fl/fl};Rhoc^{-/-}$ | Tukey one-way ANOVA | 30.01 | 16.32 to 43.70 | **<0.0001** |
| $Rhoa^{fl/fl}$ vs $Rhoa^{\beta MHC\text{-}cre};Rhoc^{-/-}$ | Tukey one-way ANOVA | -0.4918 | -14.18 to 13.20 | >0.9999 |
| $Rhoa^{\beta MHC\text{-}cre}$ vs $Rhoa^{fl/fl};Rhoc^{-/-}$ | Tukey one-way ANOVA | 34.19 | 20.50 to 47.88 | **<0.0001** |
| $Rhoa^{\beta MHC\text{-}cre}$ vs $Rhoa^{\beta MHC\text{-}cre};Rhoc^{-/-}$ | Tukey one-way ANOVA | 3.683 | -10.01 to 17.37 | 0.9311 |
| $Rhoa^{fl/fl};Rhoc^{-/-}$ vs $Rhoa^{\beta MHC\text{-}cre};Rhoc^{-/-}$ | Tukey one-way ANOVA | -30.51 | -44.20 to -16.81 | **<0.0001** |
| **Figure 1, Panel F — FS (8 wks)** | | | | |
| **Comparison** | **Test** | **Mean Diff.** | **95% CI** | **Adjusted P** |
| Omnibus one-way ANOVA | One-way ANOVA | F(4,44)=1.394 | — | 0.252 |
| $WT^{\beta MHC\text{-}cre}$ vs $Rhoa^{fl/fl}$ | Tukey one-way ANOVA | -1.236 | -5.912 to 3.440 | 0.9428 |
| $WT^{\beta MHC\text{-}cre}$ vs $Rhoa^{\beta MHC\text{-}cre}$ | Tukey one-way ANOVA | -0.3869 | -5.227 to 4.453 | 0.9994 |
| $WT^{\beta MHC\text{-}cre}$ vs $Rhoa^{fl/fl};Rhoc^{-/-}$ | Tukey one-way ANOVA | -3.038 | -7.579 to 1.502 | 0.3308 |
| $WT^{\beta MHC\text{-}cre}$ vs $Rhoa^{\beta MHC\text{-}cre};Rhoc^{-/-}$ | Tukey one-way ANOVA | -2.778 | -7.319 to 1.762 | 0.4206 |
| $Rhoa^{fl/fl}$ vs $Rhoa^{\beta MHC\text{-}cre}$ | Tukey one-way ANOVA | 0.8492 | -4.303 to 6.002 | 0.9898 |
| $Rhoa^{fl/fl}$ vs $Rhoa^{fl/fl};Rhoc^{-/-}$ | Tukey one-way ANOVA | -1.802 | -6.674 to 3.070 | 0.8296 |

| | | | | |
|---|---|---|---|---|
| *Rhoa*$^{fl/fl}$ vs *Rhoa*$^{\beta MHC\text{-}cre}$;*Rhoc*$^{-/-}$ | Tukey one-way ANOVA | -1.542 | -6.414 to 3.330 | 0.8952 |
| *Rhoa*$^{\beta MHC\text{-}cre}$ vs *Rhoa*$^{fl/fl}$;*Rhoc*$^{-/-}$ | Tukey one-way ANOVA | -2.651 | -7.681 to 2.378 | 0.5684 |
| *Rhoa*$^{\beta MHC\text{-}cre}$ vs *Rhoa*$^{\beta MHC\text{-}cre}$;*Rhoc*$^{-/-}$ | Tukey one-way ANOVA | -2.391 | -7.421 to 2.638 | 0.6606 |
| *Rhoa*$^{fl/fl}$;*Rhoc*$^{-/-}$ vs *Rhoa*$^{\beta MHC\text{-}cre}$;*Rhoc*$^{-/-}$ | Tukey one-way ANOVA | 0.2600 | -4.482 to 5.002 | 0.9999 |
| **Figure 1, Panel G — LW/BW (8 wks)** | | | | |
| **Comparison** | **Test** | **Mean Diff.** | **95% CI** | **Adjusted P** |
| Omnibus one-way ANOVA | One-way ANOVA | F(4,40)=1.322 | — | 0.279 |
| WT$^{\beta MHC\text{-}cre}$ vs *Rhoa*$^{fl/fl}$ | Tukey one-way ANOVA | 0.2045 | -0.8587 to 1.268 | 0.9814 |
| WT$^{\beta MHC\text{-}cre}$ vs *Rhoa*$^{\beta MHC\text{-}cre}$ | Tukey one-way ANOVA | 0.7150 | -0.3482 to 1.778 | 0.3233 |
| WT$^{\beta MHC\text{-}cre}$ vs *Rhoa*$^{fl/fl}$;*Rhoc*$^{-/-}$ | Tukey one-way ANOVA | -0.09249 | -1.156 to 0.9707 | 0.9991 |
| WT$^{\beta MHC\text{-}cre}$ vs *Rhoa*$^{\beta MHC\text{-}cre}$;*Rhoc*$^{-/-}$ | Tukey one-way ANOVA | 0.3661 | -0.6611 to 1.393 | 0.8456 |
| *Rhoa*$^{fl/fl}$ vs *Rhoa*$^{\beta MHC\text{-}cre}$ | Tukey one-way ANOVA | 0.5105 | -0.6542 to 1.675 | 0.7214 |
| *Rhoa*$^{fl/fl}$ vs *Rhoa*$^{fl/fl}$;*Rhoc*$^{-/-}$ | Tukey one-way ANOVA | -0.2970 | -1.462 to 0.8677 | 0.9486 |
| *Rhoa*$^{fl/fl}$ vs *Rhoa*$^{\beta MHC\text{-}cre}$;*Rhoc*$^{-/-}$ | Tukey one-way ANOVA | 0.1615 | -0.9704 to 1.293 | 0.9940 |
| *Rhoa*$^{\beta MHC\text{-}cre}$ vs *Rhoa*$^{fl/fl}$;*Rhoc*$^{-/-}$ | Tukey one-way ANOVA | -0.8075 | -1.972 to 0.3572 | 0.2940 |
| *Rhoa*$^{\beta MHC\text{-}cre}$ vs *Rhoa*$^{\beta MHC\text{-}cre}$;*Rhoc*$^{-/-}$ | Tukey one-way ANOVA | -0.3490 | -1.481 to 0.7829 | 0.9024 |
| *Rhoa*$^{fl/fl}$;*Rhoc*$^{-/-}$ vs *Rhoa*$^{\beta MHC\text{-}cre}$;*Rhoc*$^{-/-}$ | Tukey one-way ANOVA | 0.4586 | -0.6733 to 1.590 | 0.7752 |

**Figure 1, Panel H — HW/BW (8 mo)**

| Comparison | Test | Mean Diff. | 95% CI | Adjusted P |
|---|---|---|---|---|
| Omnibus one-way ANOVA | One-way ANOVA | F(4,57)=8.973 | — | **<0.0001** |
| $\mathrm{WT}^{\beta\mathrm{MHC\text{-}cre}}$ vs $Rhoa^{fl/fl}$ | Tukey one-way ANOVA | 0.0934 | -0.5957 to 0.7826 | 0.9954 |
| $\mathrm{WT}^{\beta\mathrm{MHC\text{-}cre}}$ vs $Rhoa^{\beta\mathrm{MHC\text{-}cre}}$ | Tukey one-way ANOVA | 0.0470 | -0.4873 to 0.5813 | 0.9991 |
| $\mathrm{WT}^{\beta\mathrm{MHC\text{-}cre}}$ vs $Rhoa^{fl/fl};Rhoc^{-/-}$ | Tukey one-way ANOVA | 0.0483 | -0.5129 to 0.6095 | 0.9992 |
| $\mathrm{WT}^{\beta\mathrm{MHC\text{-}cre}}$ vs $Rhoa^{\beta\mathrm{MHC\text{-}cre}};Rhoc^{-/-}$ | Tukey one-way ANOVA | -1.066 | -1.705 to -0.4268 | **<0.001** |
| $Rhoa^{fl/fl}$ vs $Rhoa^{\beta\mathrm{MHC\text{-}cre}}$ | Tukey one-way ANOVA | -0.0464 | -0.6871 to 0.5942 | 0.9996 |
| $Rhoa^{fl/fl}$ vs $Rhoa^{fl/fl};Rhoc^{-/-}$ | Tukey one-way ANOVA | -0.0452 | -0.7084 to 0.6181 | 0.9997 |
| $Rhoa^{fl/fl}$ vs $Rhoa^{\beta\mathrm{MHC\text{-}cre}};Rhoc^{-/-}$ | Tukey one-way ANOVA | -1.159 | -1.889 to -0.4289 | **<0.001** |
| $Rhoa^{\beta\mathrm{MHC\text{-}cre}}$ vs $Rhoa^{fl/fl};Rhoc^{-/-}$ | Tukey one-way ANOVA | 0.0013 | -0.4992 to 0.5017 | >0.9999 |
| $Rhoa^{\beta\mathrm{MHC\text{-}cre}}$ vs $Rhoa^{\beta\mathrm{MHC\text{-}cre}};Rhoc^{-/-}$ | Tukey one-way ANOVA | -1.113 | -1.699 to -0.5264 | **<0.0001** |
| $Rhoa^{fl/fl};Rhoc^{-/-}$ vs $Rhoa^{\beta\mathrm{MHC\text{-}cre}};Rhoc^{-/-}$ | Tukey one-way ANOVA | -1.114 | -1.725 to -0.5031 | **<0.0001** |

**Figure 1, Panel I — CSA (8 mo)**

| Comparison | Test | Mean Diff. | 95% CI | Adjusted P |
|---|---|---|---|---|
| Omnibus one-way ANOVA | One-way ANOVA | F(4,26)=9.622 | — | **<0.0001** |
| $\mathrm{WT}^{\beta\mathrm{MHC\text{-}cre}}$ vs $Rhoa^{fl/fl}$ | Tukey one-way ANOVA | 2.159 | -20.62 to 24.94 | 0.9986 |
| $\mathrm{WT}^{\beta\mathrm{MHC\text{-}cre}}$ vs $Rhoa^{\beta\mathrm{MHC\text{-}cre}}$ | Tukey one-way ANOVA | 3.117 | -17.41 to 23.65 | 0.9914 |
| $\mathrm{WT}^{\beta\mathrm{MHC\text{-}cre}}$ vs $Rhoa^{fl/fl};Rhoc^{-/-}$ | Tukey one-way ANOVA | 18.30 | -3.506 to 40.11 | 0.132 |

| | | | | |
|---|---|---|---|---|
| $WT^{βMHC\text{-}cre}$ vs *Rhoa*$^{βMHC\text{-}cre}$;*Rhoc*$^{-/-}$ | Tukey one-way ANOVA | -23.05 | -44.13 to -1.959 | **0.0271** |
| *Rhoa*$^{fl/fl}$ vs *Rhoa*$^{βMHC\text{-}cre}$ | Tukey one-way ANOVA | 0.9576 | -19.57 to 21.49 | >0.9999 |
| *Rhoa*$^{fl/fl}$ vs *Rhoa*$^{fl/fl}$;*Rhoc*$^{-/-}$ | Tukey one-way ANOVA | 16.14 | -5.665 to 37.95 | 0.223 |
| *Rhoa*$^{fl/fl}$ vs *Rhoa*$^{βMHC\text{-}cre}$;*Rhoc*$^{-/-}$ | Tukey one-way ANOVA | -25.21 | -46.29 to -4.118 | **0.0134** |
| *Rhoa*$^{βMHC\text{-}cre}$ vs *Rhoa*$^{fl/fl}$;*Rhoc*$^{-/-}$ | Tukey one-way ANOVA | 15.19 | -4.265 to 34.63 | 0.181 |
| *Rhoa*$^{βMHC\text{-}cre}$ vs *Rhoa*$^{βMHC\text{-}cre}$;*Rhoc*$^{-/-}$ | Tukey one-way ANOVA | -26.16 | -44.80 to -7.524 | **0.0030** |
| *Rhoa*$^{fl/fl}$;*Rhoc*$^{-/-}$ vs *Rhoa*$^{βMHC\text{-}cre}$;*Rhoc*$^{-/-}$ | Tukey one-way ANOVA | -41.35 | -61.38 to -21.31 | **<0.0001** |
| **Figure 1, Panel J — FS (8 mo)** | | | | |
| **Comparison** | **Test** | **Mean Diff.** | **95% CI** | **Adjusted P** |
| Omnibus one-way ANOVA | One-way ANOVA | F(4,44)=0.5564 | — | 0.695 |
| $WT^{βMHC\text{-}cre}$ vs *Rhoa*$^{fl/fl}$ | Tukey one-way ANOVA | 1.961 | -6.197 to 10.12 | 0.959 |
| $WT^{βMHC\text{-}cre}$ vs *Rhoa*$^{βMHC\text{-}cre}$ | Tukey one-way ANOVA | 2.408 | -4.786 to 9.603 | 0.875 |
| $WT^{βMHC\text{-}cre}$ vs *Rhoa*$^{fl/fl}$;*Rhoc*$^{-/-}$ | Tukey one-way ANOVA | 1.558 | -5.636 to 8.753 | 0.972 |
| $WT^{βMHC\text{-}cre}$ vs *Rhoa*$^{βMHC\text{-}cre}$;*Rhoc*$^{-/-}$ | Tukey one-way ANOVA | -0.6450 | -8.122 to 6.832 | 0.9992 |
| *Rhoa*$^{fl/fl}$ vs *Rhoa*$^{βMHC\text{-}cre}$ | Tukey one-way ANOVA | 0.4476 | -7.049 to 7.944 | 0.9998 |
| *Rhoa*$^{fl/fl}$ vs *Rhoa*$^{fl/fl}$;*Rhoc*$^{-/-}$ | Tukey one-way ANOVA | -0.4024 | -7.899 to 7.094 | 0.9999 |
| *Rhoa*$^{fl/fl}$ vs *Rhoa*$^{βMHC\text{-}cre}$;*Rhoc*$^{-/-}$ | Tukey one-way ANOVA | -2.606 | -10.37 to 5.162 | 0.874 |

| | | | | |
|---|---|---|---|---|
| $Rhoa^{\beta MHC\text{-}cre}$ vs $Rhoa^{fl/fl}$;$Rhoc^{-/-}$ | Tukey one-way ANOVA | -0.8500 | -7.285 to 5.585 | 0.9956 |
| $Rhoa^{\beta MHC\text{-}cre}$ vs $Rhoa^{\beta MHC\text{-}cre}$;$Rhoc^{-/-}$ | Tukey one-way ANOVA | -3.053 | -9.802 to 3.696 | 0.701 |
| $Rhoa^{fl/fl}$;$Rhoc^{-/-}$ vs $Rhoa^{\beta MHC\text{-}cre}$;$Rhoc^{-/-}$ | Tukey one-way ANOVA | -2.203 | -8.952 to 4.546 | 0.884 |
| **Figure 1, Panel K — LW/BW (8 mo)** | | | | |
| **Comparison** | **Test** | **Mean Diff.** | **95% CI** | **Adjusted P** |
| Omnibus one-way ANOVA | One-way ANOVA | F(4,55)=7.328 | — | **<0.0001** |
| $WT^{\beta MHC\text{-}cre}$ vs $Rhoa^{fl/fl}$ | Tukey one-way ANOVA | 0.6735 | -0.7030 to 2.050 | 0.643 |
| $WT^{\beta MHC\text{-}cre}$ vs $Rhoa^{\beta MHC\text{-}cre}$ | Tukey one-way ANOVA | 0.3901 | -0.7011 to 1.481 | 0.851 |
| $WT^{\beta MHC\text{-}cre}$ vs $Rhoa^{fl/fl}$;$Rhoc^{-/-}$ | Tukey one-way ANOVA | -0.1130 | -1.234 to 1.008 | 0.999 |
| $WT^{\beta MHC\text{-}cre}$ vs $Rhoa^{\beta MHC\text{-}cre}$;$Rhoc^{-/-}$ | Tukey one-way ANOVA | -1.680 | -2.957 to -0.4042 | **0.0042** |
| $Rhoa^{fl/fl}$ vs $Rhoa^{\beta MHC\text{-}cre}$ | Tukey one-way ANOVA | -0.2834 | -1.583 to 1.016 | 0.972 |
| $Rhoa^{fl/fl}$ vs $Rhoa^{fl/fl}$;$Rhoc^{-/-}$ | Tukey one-way ANOVA | -0.7865 | -2.111 to 0.5383 | 0.458 |
| $Rhoa^{fl/fl}$ vs $Rhoa^{\beta MHC\text{-}cre}$;$Rhoc^{-/-}$ | Tukey one-way ANOVA | -2.354 | -3.813 to -0.8954 | **<0.001** |
| $Rhoa^{\beta MHC\text{-}cre}$ vs $Rhoa^{fl/fl}$;$Rhoc^{-/-}$ | Tukey one-way ANOVA | -0.5031 | -1.528 to 0.5222 | 0.640 |
| $Rhoa^{\beta MHC\text{-}cre}$ vs $Rhoa^{\beta MHC\text{-}cre}$;$Rhoc^{-/-}$ | Tukey one-way ANOVA | -2.071 | -3.264 to -0.8775 | **<0.0001** |
| $Rhoa^{fl/fl}$;$Rhoc^{-/-}$ vs $Rhoa^{\beta MHC\text{-}cre}$;$Rhoc^{-/-}$ | Tukey one-way ANOVA | -1.567 | -2.788 to -0.3471 | **0.0055** |

### Supplementary Table 2. Statistical comparisons for Figure 2.

| **Panel B — Length um** | | | | |
|---|---|---|---|---|
| **Comparison** | **Test** | **Mean Diff.** | **95% CI** | **Adjusted P** |
| Omnibus one-way ANOVA | One-way ANOVA | F(3,8)=10.39 | — | **0.00390** |
| *Rhoa*$^{fl/fl}$ vs *Rhoa*$^{\beta MHC\text{-}cre}$ | Tukey one-way ANOVA | -12.04 | -30.12 to 6.044 | 0.222 |
| *Rhoa*$^{fl/fl}$ vs *Rhoa*$^{fl/fl}$;*Rhoc*$^{-/-}$ | Tukey one-way ANOVA | -20.63 | -38.71 to -2.551 | **0.0266** |
| *Rhoa*$^{fl/fl}$ vs *Rhoa*$^{\beta MHC\text{-}cre}$;*Rhoc*$^{-/-}$ | Tukey one-way ANOVA | -30.28 | -48.36 to -12.20 | **0.0030** |
| *Rhoa*$^{\beta MHC\text{-}cre}$ vs *Rhoa*$^{fl/fl}$;*Rhoc*$^{-/-}$ | Tukey one-way ANOVA | -8.595 | -26.67 to 9.485 | 0.469 |
| *Rhoa*$^{\beta MHC\text{-}cre}$ vs *Rhoa*$^{\beta MHC\text{-}cre}$;*Rhoc*$^{-/-}$ | Tukey one-way ANOVA | -18.24 | -36.32 to -0.1594 | **0.0481** |
| *Rhoa*$^{fl/fl}$;*Rhoc*$^{-/-}$ vs *Rhoa*$^{\beta MHC\text{-}cre}$;*Rhoc*$^{-/-}$ | Tukey one-way ANOVA | -9.645 | -27.72 to 8.435 | 0.379 |
| **Panel C — Width um** | | | | |
| **Comparison** | **Test** | **Mean Diff.** | **95% CI** | **Adjusted P** |
| Omnibus one-way ANOVA | One-way ANOVA | F(3,8)=18.05 | — | **0.0006** |
| *Rhoa*$^{fl/fl}$ vs *Rhoa*$^{\beta MHC\text{-}cre}$ | Tukey one-way ANOVA | 2.095 | -1.372 to 5.561 | 0.287 |
| *Rhoa*$^{fl/fl}$ vs *Rhoa*$^{fl/fl}$;*Rhoc*$^{-/-}$ | Tukey one-way ANOVA | 7.615 | 4.148 to 11.08 | **0.0005** |
| *Rhoa*$^{fl/fl}$ vs *Rhoa*$^{\beta MHC\text{-}cre}$;*Rhoc*$^{-/-}$ | Tukey one-way ANOVA | 2.218 | -1.249 to 5.684 | 0.248 |
| *Rhoa*$^{\beta MHC\text{-}cre}$ vs *Rhoa*$^{fl/fl}$;*Rhoc*$^{-/-}$ | Tukey one-way ANOVA | 5.520 | 2.054 to 8.987 | **0.0041** |
| *Rhoa*$^{\beta MHC\text{-}cre}$ vs *Rhoa*$^{\beta MHC\text{-}cre}$;*Rhoc*$^{-/-}$ | Tukey one-way ANOVA | 0.1233 | -3.343 to 3.590 | 0.9994 |
| *Rhoa*$^{fl/fl}$;*Rhoc*$^{-/-}$ vs *Rhoa*$^{\beta MHC\text{-}cre}$;*Rhoc*$^{-/-}$ | Tukey one-way ANOVA | -5.397 | -8.863 to -1.931 | **0.0047** |
| **Panel D — L/W ratio** | | | | |
| **Comparison** | **Test** | **Mean Diff.** | **95% CI** | **Adjusted P** |

| | | | | |
|---|---|---|---|---|
| Omnibus one-way ANOVA | One-way ANOVA | F(3,8)=31.24 | — | **<0.0001** |
| *Rhoa*$^{fl/fl}$ vs *Rhoa*$^{βMHC-cre}$ | Tukey one-way ANOVA | -0.4557 | -1.073 to 0.1614 | 0.162 |
| *Rhoa*$^{fl/fl}$ vs *Rhoa*$^{fl/fl}$;*Rhoc*$^{-/-}$ | Tukey one-way ANOVA | -1.635 | -2.252 to -1.018 | **<0.001** |
| *Rhoa*$^{fl/fl}$ vs *Rhoa*$^{βMHC-cre}$;*Rhoc*$^{-/-}$ | Tukey one-way ANOVA | -1.347 | -1.964 to -0.7296 | **<0.001** |
| *Rhoa*$^{βMHC-cre}$ vs *Rhoa*$^{fl/fl}$;*Rhoc*$^{-/-}$ | Tukey one-way ANOVA | -1.179 | -1.796 to -0.5620 | **0.0013** |
| *Rhoa*$^{βMHC-cre}$ vs *Rhoa*$^{βMHC-cre}$;*Rhoc*$^{-/-}$ | Tukey one-way ANOVA | -0.8910 | -1.508 to -0.2738 | **0.0074** |
| *Rhoa*$^{fl/fl}$;*Rhoc*$^{-/-}$ vs *Rhoa*$^{βMHC-cre}$;*Rhoc*$^{-/-}$ | Tukey one-way ANOVA | 0.2882 | -0.3289 to 0.9053 | 0.482 |

### Supplementary Table 3. Statistical comparisons for Figure 3.

| **Panel C — HW/BW (2-wk TAC)** | | | | |
|---|---|---|---|---|
| **Comparison** | **Test** | **Mean Diff.** | **95% CI** | **Adjusted P** |
| Omnibus one-way ANOVA | One-way ANOVA | F(9,79)=39.00 | — | **<0.0001** |
| $WT^{βMHC\text{-}cre}$ Sham vs $WT^{βMHC\text{-}cre}$ TAC | Tukey one-way ANOVA | -1.602 | -3.184 to -0.01895 | **0.045** |
| $WT^{βMHC\text{-}cre}$ Sham vs *Rhoa*$^{fl/fl}$ Sham | Tukey one-way ANOVA | 0.1686 | -1.702 to 2.040 | >0.9999 |
| $WT^{βMHC\text{-}cre}$ Sham vs *Rhoa*$^{fl/fl}$ TAC | Tukey one-way ANOVA | -2.178 | -3.893 to -0.4635 | **0.0033** |
| $WT^{βMHC\text{-}cre}$ Sham vs *Rhoa*$^{βMHC\text{-}cre}$ Sham | Tukey one-way ANOVA | 0.4449 | -1.426 to 2.316 | 0.9988 |
| $WT^{βMHC\text{-}cre}$ Sham vs *Rhoa*$^{βMHC\text{-}cre}$ TAC | Tukey one-way ANOVA | -4.05 | -5.806 to -2.293 | **<0.0001** |
| $WT^{βMHC\text{-}cre}$ Sham vs *Rhoa*$^{fl/fl}$;*Rhoc*$^{-/-}$ Sham | Tukey one-way ANOVA | 0.2932 | -1.514 to 2.101 | >0.9999 |
| $WT^{βMHC\text{-}cre}$ Sham vs *Rhoa*$^{fl/fl}$;*Rhoc*$^{-/-}$ TAC | Tukey one-way ANOVA | -4.283 | -5.998 to -2.568 | **<0.0001** |
| $WT^{βMHC\text{-}cre}$ Sham vs *Rhoa*$^{βMHC\text{-}cre}$;*Rhoc*$^{-/-}$ Sham | Tukey one-way ANOVA | 0.08121 | -1.790 to 1.952 | >0.9999 |
| $WT^{βMHC\text{-}cre}$ Sham vs *Rhoa*$^{βMHC\text{-}cre}$;*Rhoc*$^{-/-}$ TAC | Tukey one-way ANOVA | -6.496 | -8.304 to -4.689 | **<0.0001** |
| $WT^{βMHC\text{-}cre}$ TAC vs *Rhoa*$^{fl/fl}$ Sham | Tukey one-way ANOVA | 1.77 | 0.1155 to 3.425 | **0.0264** |
| $WT^{βMHC\text{-}cre}$ TAC vs *Rhoa*$^{fl/fl}$ TAC | Tukey one-way ANOVA | -0.5767 | -2.053 to 0.8992 | 0.9569 |
| $WT^{βMHC\text{-}cre}$ TAC vs *Rhoa*$^{βMHC\text{-}cre}$ Sham | Tukey one-way ANOVA | 2.047 | 0.3917 to 3.701 | **0.0048** |
| $WT^{βMHC\text{-}cre}$ TAC vs *Rhoa*$^{βMHC\text{-}cre}$ TAC | Tukey one-way ANOVA | -2.448 | -3.972 to -0.9238 | **<0.0001** |

| | | | | |
|---|---|---|---|---|
| WT$^{βMHC-cre}$ TAC vs *Rhoa*$^{fl/fl}$;*Rhoc*$^{-/-}$ Sham | Tukey one-way ANOVA | 1.895 | 0.3121 to 3.478 | **0.0073** |
| WT$^{βMHC-cre}$ TAC vs *Rhoa*$^{fl/fl}$;*Rhoc*$^{-/-}$ TAC | Tukey one-way ANOVA | -2.681 | -4.157 to -1.206 | **<0.0001** |
| WT$^{βMHC-cre}$ TAC vs *Rhoa*$^{βMHC-cre}$;*Rhoc*$^{-/-}$ Sham | Tukey one-way ANOVA | 1.683 | 0.02807 to 3.338 | **0.043** |
| WT$^{βMHC-cre}$ TAC vs *Rhoa*$^{βMHC-cre}$;*Rhoc*$^{-/-}$ TAC | Tukey one-way ANOVA | -4.895 | -6.477 to -3.312 | **<0.0001** |
| *Rhoa*$^{fl/fl}$ Sham vs *Rhoa*$^{fl/fl}$ TAC | Tukey one-way ANOVA | -2.347 | -4.128 to -0.5654 | **0.002** |
| *Rhoa*$^{fl/fl}$ Sham vs *Rhoa*$^{βMHC-cre}$ Sham | Tukey one-way ANOVA | 0.2762 | -1.656 to 2.209 | >0.9999 |
| *Rhoa*$^{fl/fl}$ Sham vs *Rhoa*$^{βMHC-cre}$ TAC | Tukey one-way ANOVA | -4.218 | -6.040 to -2.396 | **<0.0001** |
| *Rhoa*$^{fl/fl}$ Sham vs *Rhoa*$^{fl/fl}$;*Rhoc*$^{-/-}$ Sham | Tukey one-way ANOVA | 0.1246 | -1.746 to 1.996 | >0.9999 |
| *Rhoa*$^{fl/fl}$ Sham vs *Rhoa*$^{fl/fl}$;*Rhoc*$^{-/-}$ TAC | Tukey one-way ANOVA | -4.452 | -6.233 to -2.670 | **<0.0001** |
| *Rhoa*$^{fl/fl}$ Sham vs *Rhoa*$^{βMHC-cre}$;*Rhoc*$^{-/-}$ Sham | Tukey one-way ANOVA | -0.08742 | -2.020 to 1.845 | >0.9999 |
| *Rhoa*$^{fl/fl}$ Sham vs *Rhoa*$^{βMHC-cre}$;*Rhoc*$^{-/-}$ TAC | Tukey one-way ANOVA | -6.665 | -8.536 to -4.794 | **<0.0001** |
| *Rhoa*$^{fl/fl}$ TAC vs *Rhoa*$^{βMHC-cre}$ Sham | Tukey one-way ANOVA | 2.623 | 0.8416 to 4.405 | **0.0003** |
| *Rhoa*$^{fl/fl}$ TAC vs *Rhoa*$^{βMHC-cre}$ TAC | Tukey one-way ANOVA | -1.871 | -3.532 to -0.2104 | **0.0151** |
| *Rhoa*$^{fl/fl}$ TAC vs *Rhoa*$^{fl/fl}$;*Rhoc*$^{-/-}$ Sham | Tukey one-way ANOVA | 2.471 | 0.7567 to 4.186 | **0.0005** |
| *Rhoa*$^{fl/fl}$ TAC vs *Rhoa*$^{fl/fl}$;*Rhoc*$^{-/-}$ TAC | Tukey one-way ANOVA | -2.105 | -3.722 to -0.4881 | **0.0023** |

| | | | | |
|---|---|---|---|---|
| *Rhoa*$^{fl/fl}$ TAC vs *Rhoa*$^{βMHC-cre}$;*Rhoc*$^{-/-}$ Sham | Tukey one-way ANOVA | 2.26 | 0.4780 to 4.041 | **0.0034** |
| *Rhoa*$^{fl/fl}$ TAC vs *Rhoa*$^{βMHC-cre}$;*Rhoc*$^{-/-}$ TAC | Tukey one-way ANOVA | -4.318 | -6.033 to -2.603 | **<0.0001** |
| *Rhoa*$^{βMHC-cre}$ Sham vs *Rhoa*$^{βMHC-cre}$ TAC | Tukey one-way ANOVA | -4.495 | -6.316 to -2.673 | **<0.0001** |
| *Rhoa*$^{βMHC-cre}$ Sham vs *Rhoa*$^{fl/fl}$;*Rhoc*$^{-/-}$ Sham | Tukey one-way ANOVA | -0.1517 | -2.023 to 1.719 | >0.9999 |
| *Rhoa*$^{βMHC-cre}$ Sham vs *Rhoa*$^{fl/fl}$;*Rhoc*$^{-/-}$ TAC | Tukey one-way ANOVA | -4.728 | -6.510 to -2.946 | **<0.0001** |
| *Rhoa*$^{βMHC-cre}$ Sham vs *Rhoa*$^{βMHC-cre}$;*Rhoc*$^{-/-}$ Sham | Tukey one-way ANOVA | -0.3637 | -2.296 to 1.569 | 0.9998 |
| *Rhoa*$^{βMHC-cre}$ Sham vs *Rhoa*$^{βMHC-cre}$;*Rhoc*$^{-/-}$ TAC | Tukey one-way ANOVA | -6.941 | -8.812 to -5.070 | **<0.0001** |
| *Rhoa*$^{βMHC-cre}$ TAC vs *Rhoa*$^{fl/fl}$;*Rhoc*$^{-/-}$ Sham | Tukey one-way ANOVA | 4.343 | 2.586 to 6.100 | **<0.0001** |
| *Rhoa*$^{βMHC-cre}$ TAC vs *Rhoa*$^{fl/fl}$;*Rhoc*$^{-/-}$ TAC | Tukey one-way ANOVA | -0.2334 | -1.894 to 1.428 | >0.9999 |
| *Rhoa*$^{βMHC-cre}$ TAC vs *Rhoa*$^{βMHC-cre}$;*Rhoc*$^{-/-}$ Sham | Tukey one-way ANOVA | 4.131 | 2.309 to 5.953 | **<0.0001** |
| *Rhoa*$^{βMHC-cre}$ TAC vs *Rhoa*$^{βMHC-cre}$;*Rhoc*$^{-/-}$ TAC | Tukey one-way ANOVA | -2.447 | -4.203 to -0.6901 | **0.0008** |
| *Rhoa*$^{fl/fl}$;*Rhoc*$^{-/-}$ Sham vs *Rhoa*$^{fl/fl}$;*Rhoc*$^{-/-}$ TAC | Tukey one-way ANOVA | -4.576 | -6.291 to -2.861 | **<0.0001** |
| *Rhoa*$^{fl/fl}$;*Rhoc*$^{-/-}$ Sham vs *Rhoa*$^{βMHC-cre}$;*Rhoc*$^{-/-}$ Sham | Tukey one-way ANOVA | -0.212 | -2.083 to 1.659 | >0.9999 |
| *Rhoa*$^{fl/fl}$;*Rhoc*$^{-/-}$ Sham vs *Rhoa*$^{βMHC-cre}$;*Rhoc*$^{-/-}$ TAC | Tukey one-way ANOVA | -6.79 | -8.597 to -4.982 | **<0.0001** |
| *Rhoa*$^{fl/fl}$;*Rhoc*$^{-/-}$ TAC vs *Rhoa*$^{βMHC-cre}$;*Rhoc*$^{-/-}$ Sham | Tukey one-way ANOVA | 4.364 | 2.583 to 6.146 | **<0.0001** |

| | | | | |
|---|---|---|---|---|
| *Rhoa*$^{fl/fl}$;*Rhoc*$^{-/-}$ TAC vs *Rhoa*$^{βMHC\text{-}cre}$;*Rhoc*$^{-/-}$ TAC | Tukey one-way ANOVA | -2.213 | -3.928 to -0.4985 | **0.0026** |
| *Rhoa*$^{βMHC\text{-}cre}$;*Rhoc*$^{-/-}$ Sham vs *Rhoa*$^{βMHC\text{-}cre}$;*Rhoc*$^{-/-}$ TAC | Tukey one-way ANOVA | -6.578 | -8.449 to -4.707 | **<0.0001** |
| **Panel D — CSA µm² (2-wk TAC)** | | | | |
| **Comparison** | **Test** | **Mean Diff.** | **95% CI** | **Adjusted P** |
| Omnibus one-way ANOVA | One-way ANOVA | $F(9,59)=109.2$ | — | **<0.0001** |
| WT$^{βMHC\text{-}cre}$ Sham vs WT$^{βMHC\text{-}cre}$ TAC | Tukey one-way ANOVA | -97.57 | -130.7 to -64.44 | **<0.0001** |
| WT$^{βMHC\text{-}cre}$ Sham vs *Rhoa*$^{fl/fl}$ Sham | Tukey one-way ANOVA | 1.490 | -31.64 to 34.62 | >0.9999 |
| WT$^{βMHC\text{-}cre}$ Sham vs *Rhoa*$^{fl/fl}$ TAC | Tukey one-way ANOVA | -97.77 | -130.9 to -64.64 | **<0.0001** |
| WT$^{βMHC\text{-}cre}$ Sham vs *Rhoa*$^{βMHC\text{-}cre}$ Sham | Tukey one-way ANOVA | -1.704 | -34.84 to 31.43 | >0.9999 |
| WT$^{βMHC\text{-}cre}$ Sham vs *Rhoa*$^{βMHC\text{-}cre}$ TAC | Tukey one-way ANOVA | -188.2 | -220.2 to -156.1 | **<0.0001** |
| WT$^{βMHC\text{-}cre}$ Sham vs *Rhoa*$^{fl/fl}$;*Rhoc*$^{-/-}$ Sham | Tukey one-way ANOVA | 30.68 | -2.451 to 63.81 | 0.0920 |
| WT$^{βMHC\text{-}cre}$ Sham vs *Rhoa*$^{fl/fl}$;*Rhoc*$^{-/-}$ TAC | Tukey one-way ANOVA | -85.61 | -120.1 to -51.13 | **<0.0001** |
| WT$^{βMHC\text{-}cre}$ Sham vs *Rhoa*$^{βMHC\text{-}cre}$;*Rhoc*$^{-/-}$ Sham | Tukey one-way ANOVA | 0.7107 | -32.42 to 33.84 | >0.9999 |
| WT$^{βMHC\text{-}cre}$ Sham vs *Rhoa*$^{βMHC\text{-}cre}$;*Rhoc*$^{-/-}$ TAC | Tukey one-way ANOVA | -137.1 | -171.6 to -102.6 | **<0.0001** |
| WT$^{βMHC\text{-}cre}$ TAC vs *Rhoa*$^{fl/fl}$ Sham | Tukey one-way ANOVA | 99.06 | 65.93 to 132.2 | **<0.0001** |
| WT$^{βMHC\text{-}cre}$ TAC vs *Rhoa*$^{fl/fl}$ TAC | Tukey one-way ANOVA | -0.1997 | -33.33 to 32.93 | >0.9999 |
| WT$^{βMHC\text{-}cre}$ TAC vs *Rhoa*$^{βMHC\text{-}cre}$ Sham | Tukey one-way ANOVA | 95.87 | 62.74 to 129.0 | **<0.0001** |

| | | | | |
|---|---|---|---|---|
| WT$^{\beta MHC\text{-}cre}$ TAC vs *Rhoa*$^{\beta MHC\text{-}cre}$ TAC | Tukey one-way ANOVA | -90.59 | -122.7 to -58.51 | **<0.0001** |
| WT$^{\beta MHC\text{-}cre}$ TAC vs *Rhoa*$^{fl/fl}$;*Rhoc*$^{-/-}$ Sham | Tukey one-way ANOVA | 128.3 | 95.12 to 161.4 | **<0.0001** |
| WT$^{\beta MHC\text{-}cre}$ TAC vs *Rhoa*$^{fl/fl}$;*Rhoc*$^{-/-}$ TAC | Tukey one-way ANOVA | 11.96 | -22.52 to 46.45 | 0.9782 |
| WT$^{\beta MHC\text{-}cre}$ TAC vs *Rhoa*$^{\beta MHC\text{-}cre}$;*Rhoc*$^{-/-}$ Sham | Tukey one-way ANOVA | 98.29 | 65.15 to 131.4 | **<0.0001** |
| WT$^{\beta MHC\text{-}cre}$ TAC vs *Rhoa*$^{\beta MHC\text{-}cre}$;*Rhoc*$^{-/-}$ TAC | Tukey one-way ANOVA | -39.52 | -74.01 to -5.039 | **0.0130** |
| *Rhoa*$^{fl/fl}$ Sham vs *Rhoa*$^{fl/fl}$ TAC | Tukey one-way ANOVA | -99.26 | -132.4 to -66.13 | **<0.0001** |
| *Rhoa*$^{fl/fl}$ Sham vs *Rhoa*$^{\beta MHC\text{-}cre}$ Sham | Tukey one-way ANOVA | -3.194 | -36.33 to 29.94 | >0.9999 |
| *Rhoa*$^{fl/fl}$ Sham vs *Rhoa*$^{\beta MHC\text{-}cre}$ TAC | Tukey one-way ANOVA | -189.7 | -221.7 to -157.6 | **<0.0001** |
| *Rhoa*$^{fl/fl}$ Sham vs *Rhoa*$^{fl/fl}$;*Rhoc*$^{-/-}$ Sham | Tukey one-way ANOVA | 29.19 | -3.940 to 62.33 | 0.1297 |
| *Rhoa*$^{fl/fl}$ Sham vs *Rhoa*$^{fl/fl}$;*Rhoc*$^{-/-}$ TAC | Tukey one-way ANOVA | -87.10 | -121.6 to -52.62 | **<0.0001** |
| *Rhoa*$^{fl/fl}$ Sham vs *Rhoa*$^{\beta MHC\text{-}cre}$;*Rhoc*$^{-/-}$ Sham | Tukey one-way ANOVA | -0.7789 | -33.91 to 32.35 | >0.9999 |
| *Rhoa*$^{fl/fl}$ Sham vs *Rhoa*$^{\beta MHC\text{-}cre}$;*Rhoc*$^{-/-}$ TAC | Tukey one-way ANOVA | -138.6 | -173.1 to -104.1 | **<0.0001** |
| *Rhoa*$^{fl/fl}$ TAC vs *Rhoa*$^{\beta MHC\text{-}cre}$ Sham | Tukey one-way ANOVA | 96.07 | 62.94 to 129.2 | **<0.0001** |
| *Rhoa*$^{fl/fl}$ TAC vs *Rhoa*$^{\beta MHC\text{-}cre}$ TAC | Tukey one-way ANOVA | -90.39 | -122.5 to -58.31 | **<0.0001** |
| *Rhoa*$^{fl/fl}$ TAC vs *Rhoa*$^{fl/fl}$;*Rhoc*$^{-/-}$ Sham | Tukey one-way ANOVA | 128.5 | 95.32 to 161.6 | **<0.0001** |

| | | | | |
|---|---|---|---|---|
| *Rhoa*$^{fl/fl}$ TAC vs *Rhoa*$^{fl/fl}$;*Rhoc*$^{-/-}$ TAC | Tukey one-way ANOVA | 12.16 | -22.32 to 46.65 | 0.9756 |
| *Rhoa*$^{fl/fl}$ TAC vs *Rhoa*$^{βMHC-cre}$;*Rhoc*$^{-/-}$ Sham | Tukey one-way ANOVA | 98.49 | 65.35 to 131.6 | **<0.0001** |
| *Rhoa*$^{fl/fl}$ TAC vs *Rhoa*$^{βMHC-cre}$;*Rhoc*$^{-/-}$ TAC | Tukey one-way ANOVA | -39.32 | -73.81 to -4.839 | **0.0137** |
| *Rhoa*$^{βMHC-cre}$ Sham vs *Rhoa*$^{βMHC-cre}$ TAC | Tukey one-way ANOVA | -186.5 | -218.5 to -154.4 | **<0.0001** |
| *Rhoa*$^{βMHC-cre}$ Sham vs *Rhoa*$^{fl/fl}$;*Rhoc*$^{-/-}$ Sham | Tukey one-way ANOVA | 32.39 | -0.7462 to 65.52 | 0.0605 |
| *Rhoa*$^{βMHC-cre}$ Sham vs *Rhoa*$^{fl/fl}$;*Rhoc*$^{-/-}$ TAC | Tukey one-way ANOVA | -83.91 | -118.4 to -49.42 | **<0.0001** |
| *Rhoa*$^{βMHC-cre}$ Sham vs *Rhoa*$^{βMHC-cre}$;*Rhoc*$^{-/-}$ Sham | Tukey one-way ANOVA | 2.415 | -30.72 to 35.55 | >0.9999 |
| *Rhoa*$^{βMHC-cre}$ Sham vs *Rhoa*$^{βMHC-cre}$;*Rhoc*$^{-/-}$ TAC | Tukey one-way ANOVA | -135.4 | -169.9 to -100.9 | **<0.0001** |
| *Rhoa*$^{βMHC-cre}$ TAC vs *Rhoa*$^{fl/fl}$;*Rhoc*$^{-/-}$ Sham | Tukey one-way ANOVA | 218.8 | 186.8 to 250.9 | **<0.0001** |
| *Rhoa*$^{βMHC-cre}$ TAC vs *Rhoa*$^{fl/fl}$;*Rhoc*$^{-/-}$ TAC | Tukey one-way ANOVA | 102.5 | 69.07 to 136.0 | **<0.0001** |
| *Rhoa*$^{βMHC-cre}$ TAC vs *Rhoa*$^{βMHC-cre}$;*Rhoc*$^{-/-}$ Sham | Tukey one-way ANOVA | 188.9 | 156.8 to 221.0 | **<0.0001** |
| *Rhoa*$^{βMHC-cre}$ TAC vs *Rhoa*$^{βMHC-cre}$;*Rhoc*$^{-/-}$ TAC | Tukey one-way ANOVA | 51.06 | 17.59 to 84.54 | **0.0002** |
| *Rhoa*$^{fl/fl}$;*Rhoc*$^{-/-}$ Sham vs *Rhoa*$^{fl/fl}$;*Rhoc*$^{-/-}$ TAC | Tukey one-way ANOVA | -116.3 | -150.8 to -81.81 | **<0.0001** |
| *Rhoa*$^{fl/fl}$;*Rhoc*$^{-/-}$ Sham vs *Rhoa*$^{βMHC-cre}$;*Rhoc*$^{-/-}$ Sham | Tukey one-way ANOVA | -29.97 | -63.10 to 3.161 | 0.1087 |
| *Rhoa*$^{fl/fl}$;*Rhoc*$^{-/-}$ Sham vs *Rhoa*$^{βMHC-cre}$;*Rhoc*$^{-/-}$ TAC | Tukey one-way ANOVA | -167.8 | -202.3 to -133.3 | **<0.0001** |

| | | | | |
|---|---|---|---|---|
| *Rhoa*$^{fl/fl}$;*Rhoc*$^{-/-}$ TAC vs *Rhoa*$^{βMHC\text{-}cre}$;*Rhoc*$^{-/-}$ Sham | Tukey one-way ANOVA | 86.32 | 51.84 to 120.8 | **<0.0001** |
| *Rhoa*$^{fl/fl}$;*Rhoc*$^{-/-}$ TAC vs *Rhoa*$^{βMHC\text{-}cre}$;*Rhoc*$^{-/-}$ TAC | Tukey one-way ANOVA | -51.49 | -87.27 to -15.70 | **0.0006** |
| *Rhoa*$^{βMHC\text{-}cre}$;*Rhoc*$^{-/-}$ Sham vs *Rhoa*$^{βMHC\text{-}cre}$;*Rhoc*$^{-/-}$ TAC | Tukey one-way ANOVA | -137.8 | -172.3 to -103.3 | **<0.0001** |
| **Panel E — FS % (2-wk TAC)** | | | | |
| **Comparison** | **Test** | **Mean Diff.** | **95% CI** | **Adjusted P** |
| Omnibus one-way ANOVA | One-way ANOVA | F(9,73)=10.42 | — | **<0.0001** |
| WT$^{βMHC\text{-}cre}$ Sham vs WT$^{βMHC\text{-}cre}$ TAC | Tukey one-way ANOVA | 1.541 | -4.371 to 7.453 | 0.9974 |
| WT$^{βMHC\text{-}cre}$ Sham vs *Rhoa*$^{fl/fl}$ Sham | Tukey one-way ANOVA | 0.6929 | -5.892 to 7.278 | >0.9999 |
| WT$^{βMHC\text{-}cre}$ Sham vs *Rhoa*$^{fl/fl}$ TAC | Tukey one-way ANOVA | 0.9875 | -5.374 to 7.349 | >0.9999 |
| WT$^{βMHC\text{-}cre}$ Sham vs *Rhoa*$^{βMHC\text{-}cre}$ Sham | Tukey one-way ANOVA | -0.9071 | -7.492 to 5.678 | >0.9999 |
| WT$^{βMHC\text{-}cre}$ Sham vs *Rhoa*$^{βMHC\text{-}cre}$ TAC | Tukey one-way ANOVA | 1.125 | -5.237 to 7.487 | 0.9999 |
| WT$^{βMHC\text{-}cre}$ Sham vs *Rhoa*$^{fl/fl}$;*Rhoc*$^{-/-}$ Sham | Tukey one-way ANOVA | -2.413 | -8.774 to 3.949 | 0.9636 |
| WT$^{βMHC\text{-}cre}$ Sham vs *Rhoa*$^{fl/fl}$;*Rhoc*$^{-/-}$ TAC | Tukey one-way ANOVA | 5.732 | -0.1805 to 11.64 | 0.065 |
| WT$^{βMHC\text{-}cre}$ Sham vs *Rhoa*$^{βMHC\text{-}cre}$;*Rhoc*$^{-/-}$ Sham | Tukey one-way ANOVA | -3.593 | -10.18 to 2.992 | 0.7449 |
| WT$^{βMHC\text{-}cre}$ Sham vs *Rhoa*$^{βMHC\text{-}cre}$;*Rhoc*$^{-/-}$ TAC | Tukey one-way ANOVA | 11.89 | 5.526 to 18.25 | **<0.0001** |
| WT$^{βMHC\text{-}cre}$ TAC vs *Rhoa*$^{fl/fl}$ Sham | Tukey one-way ANOVA | -0.8481 | -7.000 to 5.304 | >0.9999 |
| WT$^{βMHC\text{-}cre}$ TAC vs *Rhoa*$^{fl/fl}$ TAC | Tukey one-way ANOVA | -0.5534 | -6.466 to 5.359 | >0.9999 |

| | | | | |
|---|---|---|---|---|
| WT$^{βMHC-cre}$ TAC vs *Rhoa*$^{βMHC-cre}$ Sham | Tukey one-way ANOVA | -2.448 | -8.600 to 3.704 | 0.9509 |
| WT$^{βMHC-cre}$ TAC vs *Rhoa*$^{βMHC-cre}$ TAC | Tukey one-way ANOVA | -0.4159 | -6.328 to 5.496 | >0.9999 |
| WT$^{βMHC-cre}$ TAC vs *Rhoa*$^{fl/fl}$;*Rhoc*$^{-/-}$ Sham | Tukey one-way ANOVA | -3.953 | -9.866 to 1.959 | 0.478 |
| WT$^{βMHC-cre}$ TAC vs *Rhoa*$^{fl/fl}$;*Rhoc*$^{-/-}$ TAC | Tukey one-way ANOVA | 4.191 | -1.235 to 9.616 | 0.2751 |
| WT$^{βMHC-cre}$ TAC vs *Rhoa*$^{βMHC-cre}$;*Rhoc*$^{-/-}$ Sham | Tukey one-way ANOVA | -5.134 | -11.29 to 1.018 | 0.184 |
| WT$^{βMHC-cre}$ TAC vs *Rhoa*$^{βMHC-cre}$;*Rhoc*$^{-/-}$ TAC | Tukey one-way ANOVA | 10.35 | 4.434 to 16.26 | **<0.0001** |
| *Rhoa*$^{fl/fl}$ Sham vs *Rhoa*$^{fl/fl}$ TAC | Tukey one-way ANOVA | 0.2946 | -6.291 to 6.880 | >0.9999 |
| *Rhoa*$^{fl/fl}$ Sham vs *Rhoa*$^{βMHC-cre}$ Sham | Tukey one-way ANOVA | -1.6 | -8.401 to 5.201 | 0.9988 |
| *Rhoa*$^{fl/fl}$ Sham vs *Rhoa*$^{βMHC-cre}$ TAC | Tukey one-way ANOVA | 0.4321 | -6.153 to 7.017 | >0.9999 |
| *Rhoa*$^{fl/fl}$ Sham vs *Rhoa*$^{fl/fl}$;*Rhoc*$^{-/-}$ Sham | Tukey one-way ANOVA | -3.105 | -9.691 to 3.480 | 0.8717 |
| *Rhoa*$^{fl/fl}$ Sham vs *Rhoa*$^{fl/fl}$;*Rhoc*$^{-/-}$ TAC | Tukey one-way ANOVA | 5.039 | -1.113 to 11.19 | 0.2043 |
| *Rhoa*$^{fl/fl}$ Sham vs *Rhoa*$^{βMHC-cre}$;*Rhoc*$^{-/-}$ Sham | Tukey one-way ANOVA | -4.286 | -11.09 to 2.516 | 0.5633 |
| *Rhoa*$^{fl/fl}$ Sham vs *Rhoa*$^{βMHC-cre}$;*Rhoc*$^{-/-}$ TAC | Tukey one-way ANOVA | 11.19 | 4.609 to 17.78 | **<0.0001** |
| *Rhoa*$^{fl/fl}$ TAC vs *Rhoa*$^{βMHC-cre}$ Sham | Tukey one-way ANOVA | -1.895 | -8.480 to 4.691 | 0.9946 |
| *Rhoa*$^{fl/fl}$ TAC vs *Rhoa*$^{βMHC-cre}$ TAC | Tukey one-way ANOVA | 0.1375 | -6.224 to 6.499 | >0.9999 |

| | | | | |
|---|---|---|---|---|
| *Rhoa*$^{fl/fl}$ TAC vs *Rhoa*$^{fl/fl}$;*Rhoc*$^{-/-}$ Sham | Tukey one-way ANOVA | -3.4 | -9.762 to 2.962 | 0.7667 |
| *Rhoa*$^{fl/fl}$ TAC vs *Rhoa*$^{fl/fl}$;*Rhoc*$^{-/-}$ TAC | Tukey one-way ANOVA | 4.744 | -1.168 to 10.66 | 0.2279 |
| *Rhoa*$^{fl/fl}$ TAC vs *Rhoa*$^{\beta MHC-cre}$;*Rhoc*$^{-/-}$ Sham | Tukey one-way ANOVA | -4.58 | -11.17 to 2.005 | 0.4207 |
| *Rhoa*$^{fl/fl}$ TAC vs *Rhoa*$^{\beta MHC-cre}$;*Rhoc*$^{-/-}$ TAC | Tukey one-way ANOVA | 10.9 | 4.538 to 17.26 | **<0.0001** |
| *Rhoa*$^{\beta MHC-cre}$ Sham vs *Rhoa*$^{\beta MHC-cre}$ TAC | Tukey one-way ANOVA | 2.032 | -4.553 to 8.617 | 0.9909 |
| *Rhoa*$^{\beta MHC-cre}$ Sham vs *Rhoa*$^{fl/fl}$;*Rhoc*$^{-/-}$ Sham | Tukey one-way ANOVA | -1.505 | -8.091 to 5.080 | 0.9991 |
| *Rhoa*$^{\beta MHC-cre}$ Sham vs *Rhoa*$^{fl/fl}$;*Rhoc*$^{-/-}$ TAC | Tukey one-way ANOVA | 6.639 | 0.4870 to 12.79 | **0.0242** |
| *Rhoa*$^{\beta MHC-cre}$ Sham vs *Rhoa*$^{\beta MHC-cre}$;*Rhoc*$^{-/-}$ Sham | Tukey one-way ANOVA | -2.686 | -9.487 to 4.116 | 0.9531 |
| *Rhoa*$^{\beta MHC-cre}$ Sham vs *Rhoa*$^{\beta MHC-cre}$;*Rhoc*$^{-/-}$ TAC | Tukey one-way ANOVA | 12.79 | 6.209 to 19.38 | **<0.0001** |
| *Rhoa*$^{\beta MHC-cre}$ TAC vs *Rhoa*$^{fl/fl}$;*Rhoc*$^{-/-}$ Sham | Tukey one-way ANOVA | -3.538 | -9.899 to 2.824 | 0.7239 |
| *Rhoa*$^{\beta MHC-cre}$ TAC vs *Rhoa*$^{fl/fl}$;*Rhoc*$^{-/-}$ TAC | Tukey one-way ANOVA | 4.607 | -1.306 to 10.52 | 0.264 |
| *Rhoa*$^{\beta MHC-cre}$ TAC vs *Rhoa*$^{\beta MHC-cre}$;*Rhoc*$^{-/-}$ Sham | Tukey one-way ANOVA | -4.718 | -11.30 to 1.867 | 0.3781 |
| *Rhoa*$^{\beta MHC-cre}$ TAC vs *Rhoa*$^{\beta MHC-cre}$;*Rhoc*$^{-/-}$ TAC | Tukey one-way ANOVA | 10.76 | 4.401 to 17.12 | **<0.0001** |
| *Rhoa*$^{fl/fl}$;*Rhoc*$^{-/-}$ Sham vs *Rhoa*$^{fl/fl}$;*Rhoc*$^{-/-}$ TAC | Tukey one-way ANOVA | 8.144 | 2.232 to 14.06 | **0.001** |
| *Rhoa*$^{fl/fl}$;*Rhoc*$^{-/-}$ Sham vs *Rhoa*$^{\beta MHC-cre}$;*Rhoc*$^{-/-}$ Sham | Tukey one-way ANOVA | -1.18 | -7.766 to 5.405 | 0.9999 |

| | | | | |
|---|---|---|---|---|
| *Rhoa*$^{fl/fl}$;*Rhoc*$^{-/-}$ Sham vs *Rhoa*$^{βMHC-cre}$;*Rhoc*$^{-/-}$ TAC | Tukey one-way ANOVA | 14.3 | 7.938 to 20.66 | **<0.0001** |
| *Rhoa*$^{fl/fl}$;*Rhoc*$^{-/-}$ TAC vs *Rhoa*$^{βMHC-cre}$;*Rhoc*$^{-/-}$ Sham | Tukey one-way ANOVA | -9.325 | -15.48 to -3.173 | **0.0002** |
| *Rhoa*$^{fl/fl}$;*Rhoc*$^{-/-}$ TAC vs *Rhoa*$^{βMHC-cre}$;*Rhoc*$^{-/-}$ TAC | Tukey one-way ANOVA | 6.156 | 0.2433 to 12.07 | **0.0346** |
| *Rhoa*$^{βMHC-cre}$;*Rhoc*$^{-/-}$ Sham vs *Rhoa*$^{βMHC-cre}$;*Rhoc*$^{-/-}$ TAC | Tukey one-way ANOVA | 15.48 | 8.895 to 22.07 | **<0.0001** |
| **Panel F — LW/BW (2-wk TAC)** | | | | |
| **Comparison** | **Test** | **Mean Diff.** | **95% CI** | **Adjusted P** |
| Omnibus one-way ANOVA | One-way ANOVA | F(9,79)=11.06 | — | **<0.0001** |
| WT$^{βMHC-cre}$ Sham vs WT$^{βMHC-cre}$ TAC | Tukey one-way ANOVA | -0.8417 | -5.453 to 3.770 | 0.9999 |
| WT$^{βMHC-cre}$ Sham vs *Rhoa*$^{fl/fl}$ Sham | Tukey one-way ANOVA | 0.04719 | -5.464 to 5.559 | >0.9999 |
| WT$^{βMHC-cre}$ Sham vs *Rhoa*$^{fl/fl}$ TAC | Tukey one-way ANOVA | -0.4851 | -5.810 to 4.840 | >0.9999 |
| WT$^{βMHC-cre}$ Sham vs *Rhoa*$^{βMHC-cre}$ Sham | Tukey one-way ANOVA | -0.01461 | -5.526 to 5.497 | >0.9999 |
| WT$^{βMHC-cre}$ Sham vs *Rhoa*$^{βMHC-cre}$ TAC | Tukey one-way ANOVA | -4.093 | -9.418 to 1.231 | 0.2831 |
| WT$^{βMHC-cre}$ Sham vs *Rhoa*$^{fl/fl}$;*Rhoc*$^{-/-}$ Sham | Tukey one-way ANOVA | 0.04309 | -5.282 to 5.368 | >0.9999 |
| WT$^{βMHC-cre}$ Sham vs *Rhoa*$^{fl/fl}$;*Rhoc*$^{-/-}$ TAC | Tukey one-way ANOVA | -4.89 | -9.751 to -0.02961 | **0.0474** |
| WT$^{βMHC-cre}$ Sham vs *Rhoa*$^{βMHC-cre}$;*Rhoc*$^{-/-}$ Sham | Tukey one-way ANOVA | 0.4068 | -5.105 to 5.918 | >0.9999 |
| WT$^{βMHC-cre}$ Sham vs *Rhoa*$^{βMHC-cre}$;*Rhoc*$^{-/-}$ TAC | Tukey one-way ANOVA | -11.56 | -16.89 to -6.239 | **<0.0001** |
| WT$^{βMHC-cre}$ TAC vs *Rhoa*$^{fl/fl}$ Sham | Tukey one-way ANOVA | 0.8889 | -3.937 to 5.715 | 0.9998 |

| | | | | |
|---|---|---|---|---|
| $WT^{\beta MHC\text{-}cre}$ TAC vs $Rhoa^{fl/fl}$ TAC | Tukey one-way ANOVA | 0.3566 | -4.255 to 4.968 | >0.9999 |
| $WT^{\beta MHC\text{-}cre}$ TAC vs $Rhoa^{\beta MHC\text{-}cre}$ Sham | Tukey one-way ANOVA | 0.8271 | -3.999 to 5.653 | >0.9999 |
| $WT^{\beta MHC\text{-}cre}$ TAC vs $Rhoa^{\beta MHC\text{-}cre}$ TAC | Tukey one-way ANOVA | -3.252 | -7.863 to 1.360 | 0.4033 |
| $WT^{\beta MHC\text{-}cre}$ TAC vs $Rhoa^{fl/fl}$;$Rhoc^{-/-}$ Sham | Tukey one-way ANOVA | 0.8848 | -3.727 to 5.496 | 0.9998 |
| $WT^{\beta MHC\text{-}cre}$ TAC vs $Rhoa^{fl/fl}$;$Rhoc^{-/-}$ TAC | Tukey one-way ANOVA | -4.049 | -8.116 to 0.01815 | 0.052 |
| $WT^{\beta MHC\text{-}cre}$ TAC vs $Rhoa^{\beta MHC\text{-}cre}$;$Rhoc^{-/-}$ Sham | Tukey one-way ANOVA | 1.249 | -3.577 to 6.075 | 0.9976 |
| $WT^{\beta MHC\text{-}cre}$ TAC vs $Rhoa^{\beta MHC\text{-}cre}$;$Rhoc^{-/-}$ TAC | Tukey one-way ANOVA | -10.72 | -15.33 to -6.111 | **<0.0001** |
| $Rhoa^{fl/fl}$ Sham vs $Rhoa^{fl/fl}$ TAC | Tukey one-way ANOVA | -0.5323 | -6.044 to 4.979 | >0.9999 |
| $Rhoa^{fl/fl}$ Sham vs $Rhoa^{\beta MHC\text{-}cre}$ Sham | Tukey one-way ANOVA | -0.0618 | -5.754 to 5.631 | >0.9999 |
| $Rhoa^{fl/fl}$ Sham vs $Rhoa^{\beta MHC\text{-}cre}$ TAC | Tukey one-way ANOVA | -4.141 | -9.652 to 1.371 | 0.3139 |
| $Rhoa^{fl/fl}$ Sham vs $Rhoa^{fl/fl}$;$Rhoc^{-/-}$ Sham | Tukey one-way ANOVA | -0.0041 | -5.516 to 5.508 | >0.9999 |
| $Rhoa^{fl/fl}$ Sham vs $Rhoa^{fl/fl}$;$Rhoc^{-/-}$ TAC | Tukey one-way ANOVA | -4.938 | -10.00 to 0.1272 | 0.0622 |
| $Rhoa^{fl/fl}$ Sham vs $Rhoa^{\beta MHC\text{-}cre}$;$Rhoc^{-/-}$ Sham | Tukey one-way ANOVA | 0.3596 | -5.333 to 6.052 | >0.9999 |
| $Rhoa^{fl/fl}$ Sham vs $Rhoa^{\beta MHC\text{-}cre}$;$Rhoc^{-/-}$ TAC | Tukey one-way ANOVA | -11.61 | -17.12 to -6.100 | **<0.0001** |
| $Rhoa^{fl/fl}$ TAC vs $Rhoa^{\beta MHC\text{-}cre}$ Sham | Tukey one-way ANOVA | 0.4705 | -5.041 to 5.982 | >0.9999 |
| $Rhoa^{fl/fl}$ TAC vs $Rhoa^{\beta MHC\text{-}cre}$ TAC | Tukey one-way ANOVA | -3.608 | -8.933 to 1.716 | 0.4614 |

| | | | | |
|---|---|---|---|---|
| *Rhoa*$^{fl/fl}$ TAC vs *Rhoa*$^{fl/fl}$;*Rhoc*$^{-/-}$ Sham | Tukey one-way ANOVA | 0.5282 | -4.797 to 5.853 | >0.9999 |
| *Rhoa*$^{fl/fl}$ TAC vs *Rhoa*$^{fl/fl}$;*Rhoc*$^{-/-}$ TAC | Tukey one-way ANOVA | -4.405 | -9.266 to 0.4555 | 0.1091 |
| *Rhoa*$^{fl/fl}$ TAC vs *Rhoa*$^{βMHC-cre}$;*Rhoc*$^{-/-}$ Sham | Tukey one-way ANOVA | 0.8919 | -4.620 to 6.404 | >0.9999 |
| *Rhoa*$^{fl/fl}$ TAC vs *Rhoa*$^{βMHC-cre}$;*Rhoc*$^{-/-}$ TAC | Tukey one-way ANOVA | -11.08 | -16.40 to -5.754 | **<0.0001** |
| *Rhoa*$^{βMHC-cre}$ Sham vs *Rhoa*$^{βMHC-cre}$ TAC | Tukey one-way ANOVA | -4.079 | -9.590 to 1.433 | 0.3346 |
| *Rhoa*$^{βMHC-cre}$ Sham vs *Rhoa*$^{fl/fl}$;*Rhoc*$^{-/-}$ Sham | Tukey one-way ANOVA | 0.0577 | -5.454 to 5.569 | >0.9999 |
| *Rhoa*$^{βMHC-cre}$ Sham vs *Rhoa*$^{fl/fl}$;*Rhoc*$^{-/-}$ TAC | Tukey one-way ANOVA | -4.876 | -9.941 to 0.1890 | 0.069 |
| *Rhoa*$^{βMHC-cre}$ Sham vs *Rhoa*$^{βMHC-cre}$;*Rhoc*$^{-/-}$ Sham | Tukey one-way ANOVA | 0.4214 | -5.271 to 6.114 | >0.9999 |
| *Rhoa*$^{βMHC-cre}$ Sham vs *Rhoa*$^{βMHC-cre}$;*Rhoc*$^{-/-}$ TAC | Tukey one-way ANOVA | -11.55 | -17.06 to -6.038 | **<0.0001** |
| *Rhoa*$^{βMHC-cre}$ TAC vs *Rhoa*$^{fl/fl}$;*Rhoc*$^{-/-}$ Sham | Tukey one-way ANOVA | 4.137 | -1.188 to 9.461 | 0.2696 |
| *Rhoa*$^{βMHC-cre}$ TAC vs *Rhoa*$^{fl/fl}$;*Rhoc*$^{-/-}$ TAC | Tukey one-way ANOVA | -0.797 | -5.658 to 4.064 | >0.9999 |
| *Rhoa*$^{βMHC-cre}$ TAC vs *Rhoa*$^{βMHC-cre}$;*Rhoc*$^{-/-}$ Sham | Tukey one-way ANOVA | 4.5 | -1.011 to 10.01 | 0.2091 |
| *Rhoa*$^{βMHC-cre}$ TAC vs *Rhoa*$^{βMHC-cre}$;*Rhoc*$^{-/-}$ TAC | Tukey one-way ANOVA | -7.471 | -12.80 to -2.146 | **0.0007** |
| *Rhoa*$^{fl/fl}$;*Rhoc*$^{-/-}$ Sham vs *Rhoa*$^{fl/fl}$;*Rhoc*$^{-/-}$ TAC | Tukey one-way ANOVA | -4.934 | -9.794 to -0.07270 | **0.0438** |
| *Rhoa*$^{fl/fl}$;*Rhoc*$^{-/-}$ Sham vs *Rhoa*$^{βMHC-cre}$;*Rhoc*$^{-/-}$ Sham | Tukey one-way ANOVA | 0.3637 | -5.148 to 5.875 | >0.9999 |

| | | | | |
|---|---|---|---|---|
| $Rhoa^{fl/fl};Rhoc^{-/-}$ Sham vs $Rhoa^{\beta MHC\text{-}cre};Rhoc^{-/-}$ TAC | Tukey one-way ANOVA | -11.61 | -16.93 to -6.283 | **<0.0001** |
| $Rhoa^{fl/fl};Rhoc^{-/-}$ TAC vs $Rhoa^{\beta MHC\text{-}cre};Rhoc^{-/-}$ Sham | Tukey one-way ANOVA | 5.297 | 0.2323 to 10.36 | **0.033** |
| $Rhoa^{fl/fl};Rhoc^{-/-}$ TAC vs $Rhoa^{\beta MHC\text{-}cre};Rhoc^{-/-}$ TAC | Tukey one-way ANOVA | -6.674 | -11.53 to -1.813 | **0.001** |
| $Rhoa^{\beta MHC\text{-}cre};Rhoc^{-/-}$ Sham vs $Rhoa^{\beta MHC\text{-}cre};Rhoc^{-/-}$ TAC | Tukey one-way ANOVA | -11.97 | -17.48 to -6.459 | **<0.0001** |
| **Panel H — Fibrosis % (2-wk TAC)** | | | | |
| **Comparison** | **Test** | **Mean Diff.** | **95% CI** | **Adjusted P** |
| Omnibus one-way ANOVA | One-way ANOVA | $F(9,60)=50.52$ | — | **<0.0001** |
| $WT^{\beta MHC\text{-}cre}$ Sham vs $WT^{\beta MHC\text{-}cre}$ TAC | Tukey one-way ANOVA | -5.618 | -11.09 to -0.1505 | **0.0393** |
| $WT^{\beta MHC\text{-}cre}$ Sham vs $Rhoa^{fl/fl}$ Sham | Tukey one-way ANOVA | -0.3024 | -6.074 to 5.469 | >0.9999 |
| $WT^{\beta MHC\text{-}cre}$ Sham vs $Rhoa^{fl/fl}$ TAC | Tukey one-way ANOVA | -6.611 | -12.60 to -0.6214 | **0.0195** |
| $WT^{\beta MHC\text{-}cre}$ Sham vs $Rhoa^{\beta MHC\text{-}cre}$ Sham | Tukey one-way ANOVA | -0.8785 | -6.481 to 4.724 | >0.9999 |
| $WT^{\beta MHC\text{-}cre}$ Sham vs $Rhoa^{\beta MHC\text{-}cre}$ TAC | Tukey one-way ANOVA | -15.25 | -20.72 to -9.780 | **<0.0001** |
| $WT^{\beta MHC\text{-}cre}$ Sham vs $Rhoa^{fl/fl};Rhoc^{-/-}$ Sham | Tukey one-way ANOVA | -1.13 | -7.412 to 5.152 | 0.9999 |
| $WT^{\beta MHC\text{-}cre}$ Sham vs $Rhoa^{fl/fl};Rhoc^{-/-}$ TAC | Tukey one-way ANOVA | -18.25 | -23.72 to -12.79 | **<0.0001** |
| $WT^{\beta MHC\text{-}cre}$ Sham vs $Rhoa^{\beta MHC\text{-}cre};Rhoc^{-/-}$ Sham | Tukey one-way ANOVA | -0.8516 | -6.841 to 5.138 | >0.9999 |
| $WT^{\beta MHC\text{-}cre}$ Sham vs $Rhoa^{\beta MHC\text{-}cre};Rhoc^{-/-}$ TAC | Tukey one-way ANOVA | -24.21 | -30.50 to -17.93 | **<0.0001** |
| $WT^{\beta MHC\text{-}cre}$ TAC vs $Rhoa^{fl/fl}$ Sham | Tukey one-way ANOVA | 5.316 | 0.08767 to 10.54 | **0.0432** |

| | | | | |
|---|---|---|---|---|
| WT$^{βMHC-cre}$ TAC vs *Rhoa*$^{fl/fl}$ TAC | Tukey one-way ANOVA | -0.9928 | -6.460 to 4.475 | 0.9998 |
| WT$^{βMHC-cre}$ TAC vs *Rhoa*$^{βMHC-cre}$ Sham | Tukey one-way ANOVA | 4.739 | -0.3013 to 9.780 | 0.0823 |
| WT$^{βMHC-cre}$ TAC vs *Rhoa*$^{βMHC-cre}$ TAC | Tukey one-way ANOVA | -9.63 | -14.52 to -4.740 | **<0.0001** |
| WT$^{βMHC-cre}$ TAC vs *Rhoa*$^{fl/fl}$;*Rhoc*$^{-/-}$ Sham | Tukey one-way ANOVA | 4.488 | -1.298 to 10.27 | 0.2651 |
| WT$^{βMHC-cre}$ TAC vs *Rhoa*$^{fl/fl}$;*Rhoc*$^{-/-}$ TAC | Tukey one-way ANOVA | -12.64 | -17.53 to -7.746 | **<0.0001** |
| WT$^{βMHC-cre}$ TAC vs *Rhoa*$^{βMHC-cre}$;*Rhoc*$^{-/-}$ Sham | Tukey one-way ANOVA | 4.766 | -0.7011 to 10.23 | 0.1391 |
| WT$^{βMHC-cre}$ TAC vs *Rhoa*$^{βMHC-cre}$;*Rhoc*$^{-/-}$ TAC | Tukey one-way ANOVA | -18.6 | -24.38 to -12.81 | **<0.0001** |
| *Rhoa*$^{fl/fl}$ Sham vs *Rhoa*$^{fl/fl}$ TAC | Tukey one-way ANOVA | -6.308 | -12.08 to -0.5369 | **0.0216** |
| *Rhoa*$^{fl/fl}$ Sham vs *Rhoa*$^{βMHC-cre}$ Sham | Tukey one-way ANOVA | -0.5761 | -5.945 to 4.793 | >0.9999 |
| *Rhoa*$^{fl/fl}$ Sham vs *Rhoa*$^{βMHC-cre}$ TAC | Tukey one-way ANOVA | -14.95 | -20.17 to -9.718 | **<0.0001** |
| *Rhoa*$^{fl/fl}$ Sham vs *Rhoa*$^{fl/fl}$;*Rhoc*$^{-/-}$ Sham | Tukey one-way ANOVA | -0.8278 | -6.902 to 5.247 | >0.9999 |
| *Rhoa*$^{fl/fl}$ Sham vs *Rhoa*$^{fl/fl}$;*Rhoc*$^{-/-}$ TAC | Tukey one-way ANOVA | -17.95 | -23.18 to -12.72 | **<0.0001** |
| *Rhoa*$^{fl/fl}$ Sham vs *Rhoa*$^{βMHC-cre}$;*Rhoc*$^{-/-}$ Sham | Tukey one-way ANOVA | -0.5492 | -6.321 to 5.222 | >0.9999 |
| *Rhoa*$^{fl/fl}$ Sham vs *Rhoa*$^{βMHC-cre}$;*Rhoc*$^{-/-}$ TAC | Tukey one-way ANOVA | -23.91 | -29.99 to -17.84 | **<0.0001** |
| *Rhoa*$^{fl/fl}$ TAC vs *Rhoa*$^{βMHC-cre}$ Sham | Tukey one-way ANOVA | 5.732 | 0.1297 to 11.33 | **0.0409** |
| *Rhoa*$^{fl/fl}$ TAC vs *Rhoa*$^{βMHC-cre}$ TAC | Tukey one-way ANOVA | -8.637 | -14.10 to -3.170 | **0.0001** |

| | | | | |
|---|---|---|---|---|
| *Rhoa*$^{fl/fl}$ TAC vs *Rhoa*$^{fl/fl}$;*Rhoc*$^{-/-}$ Sham | Tukey one-way ANOVA | 5.481 | -0.8011 to 11.76 | 0.1384 |
| *Rhoa*$^{fl/fl}$ TAC vs *Rhoa*$^{fl/fl}$;*Rhoc*$^{-/-}$ TAC | Tukey one-way ANOVA | -11.64 | -17.11 to -6.176 | **<0.0001** |
| *Rhoa*$^{fl/fl}$ TAC vs *Rhoa*$^{βMHC-cre}$;*Rhoc*$^{-/-}$ Sham | Tukey one-way ANOVA | 5.759 | -0.2302 to 11.75 | 0.0691 |
| *Rhoa*$^{fl/fl}$ TAC vs *Rhoa*$^{βMHC-cre}$;*Rhoc*$^{-/-}$ TAC | Tukey one-way ANOVA | -17.6 | -23.89 to -11.32 | **<0.0001** |
| *Rhoa*$^{βMHC-cre}$ Sham vs *Rhoa*$^{βMHC-cre}$ TAC | Tukey one-way ANOVA | -14.37 | -19.41 to -9.329 | **<0.0001** |
| *Rhoa*$^{βMHC-cre}$ Sham vs *Rhoa*$^{fl/fl}$;*Rhoc*$^{-/-}$ Sham | Tukey one-way ANOVA | -0.2516 | -6.166 to 5.662 | >0.9999 |
| *Rhoa*$^{βMHC-cre}$ Sham vs *Rhoa*$^{fl/fl}$;*Rhoc*$^{-/-}$ TAC | Tukey one-way ANOVA | -17.38 | -22.42 to -12.34 | **<0.0001** |
| *Rhoa*$^{βMHC-cre}$ Sham vs *Rhoa*$^{βMHC-cre}$;*Rhoc*$^{-/-}$ Sham | Tukey one-way ANOVA | 0.02693 | -5.576 to 5.629 | >0.9999 |
| *Rhoa*$^{βMHC-cre}$ Sham vs *Rhoa*$^{βMHC-cre}$;*Rhoc*$^{-/-}$ TAC | Tukey one-way ANOVA | -23.34 | -29.25 to -17.42 | **<0.0001** |
| *Rhoa*$^{βMHC-cre}$ TAC vs *Rhoa*$^{fl/fl}$;*Rhoc*$^{-/-}$ Sham | Tukey one-way ANOVA | 14.12 | 8.332 to 19.90 | **<0.0001** |
| *Rhoa*$^{βMHC-cre}$ TAC vs *Rhoa*$^{fl/fl}$;*Rhoc*$^{-/-}$ TAC | Tukey one-way ANOVA | -3.007 | -7.897 to 1.884 | 0.5889 |
| *Rhoa*$^{βMHC-cre}$ TAC vs *Rhoa*$^{βMHC-cre}$;*Rhoc*$^{-/-}$ Sham | Tukey one-way ANOVA | 14.4 | 8.929 to 19.86 | **<0.0001** |
| *Rhoa*$^{βMHC-cre}$ TAC vs *Rhoa*$^{βMHC-cre}$;*Rhoc*$^{-/-}$ TAC | Tukey one-way ANOVA | -8.967 | -14.75 to -3.181 | **0.0002** |
| *Rhoa*$^{fl/fl}$;*Rhoc*$^{-/-}$ Sham vs *Rhoa*$^{fl/fl}$;*Rhoc*$^{-/-}$ TAC | Tukey one-way ANOVA | -17.12 | -22.91 to -11.34 | **<0.0001** |
| *Rhoa*$^{fl/fl}$;*Rhoc*$^{-/-}$ Sham vs *Rhoa*$^{βMHC-cre}$;*Rhoc*$^{-/-}$ Sham | Tukey one-way ANOVA | 0.2786 | -6.003 to 6.560 | >0.9999 |

| | | | | |
|---|---|---|---|---|
| *Rhoa*$^{fl/fl}$;*Rhoc*$^{-/-}$ Sham vs *Rhoa*$^{\beta MHC\text{-}cre}$;*Rhoc*$^{-/-}$ TAC | Tukey one-way ANOVA | -23.08 | -29.65 to -16.52 | **<0.0001** |
| *Rhoa*$^{fl/fl}$;*Rhoc*$^{-/-}$ TAC vs *Rhoa*$^{\beta MHC\text{-}cre}$;*Rhoc*$^{-/-}$ Sham | Tukey one-way ANOVA | 17.4 | 11.94 to 22.87 | **<0.0001** |
| *Rhoa*$^{fl/fl}$;*Rhoc*$^{-/-}$ TAC vs *Rhoa*$^{\beta MHC\text{-}cre}$;*Rhoc*$^{-/-}$ TAC | Tukey one-way ANOVA | -5.96 | -11.75 to -0.1740 | **0.0384** |
| *Rhoa*$^{\beta MHC\text{-}cre}$;*Rhoc*$^{-/-}$ Sham vs *Rhoa*$^{\beta MHC\text{-}cre}$;*Rhoc*$^{-/-}$ TAC | Tukey one-way ANOVA | -23.36 | -29.65 to -17.08 | **<0.0001** |

**Supplementary Table 4. Statistical comparisons for Figure 4.**

| **Panel B — HW/BW (12-wk TAC)** | | | | |
|---|---|---|---|---|
| **Comparison** | **Test** | **Mean Diff.** | **95% CI** | **Adjusted P** |
| Omnibus one-way ANOVA | One-way ANOVA | F(5,32)=19.95 | — | **<0.0001** |
| $WT^{\beta MHC\text{-}cre}$ Sham vs $WT^{\beta MHC\text{-}cre}$ TAC | Tukey one-way ANOVA | -3.029 | -5.664 to -0.3940 | **0.0167** |
| $WT^{\beta MHC\text{-}cre}$ Sham vs $Rhoa^{fl/fl}$ Sham | Tukey one-way ANOVA | -0.1136 | -2.877 to 2.650 | >0.9999 |
| $WT^{\beta MHC\text{-}cre}$ Sham vs $Rhoa^{fl/fl}$ TAC | Tukey one-way ANOVA | -3.601 | -6.140 to -1.062 | **0.0019** |
| $WT^{\beta MHC\text{-}cre}$ Sham vs $Rhoa^{\beta MHC\text{-}cre}$ Sham | Tukey one-way ANOVA | -0.2170 | -2.852 to 2.418 | 0.9999 |
| $WT^{\beta MHC\text{-}cre}$ Sham vs $Rhoa^{\beta MHC\text{-}cre}$ TAC | Tukey one-way ANOVA | -6.358 | -8.822 to -3.893 | **<0.0001** |
| $WT^{\beta MHC\text{-}cre}$ TAC vs $Rhoa^{fl/fl}$ Sham | Tukey one-way ANOVA | 2.915 | 0.1518 to 5.679 | **0.0338** |
| $WT^{\beta MHC\text{-}cre}$ TAC vs $Rhoa^{fl/fl}$ TAC | Tukey one-way ANOVA | -0.5720 | -3.111 to 1.967 | 0.9827 |
| $WT^{\beta MHC\text{-}cre}$ TAC vs $Rhoa^{\beta MHC\text{-}cre}$ Sham | Tukey one-way ANOVA | 2.812 | 0.1770 to 5.447 | **0.0310** |
| $WT^{\beta MHC\text{-}cre}$ TAC vs $Rhoa^{\beta MHC\text{-}cre}$ TAC | Tukey one-way ANOVA | -3.328 | -5.793 to -0.8636 | **0.0034** |
| $Rhoa^{fl/fl}$ Sham vs $Rhoa^{fl/fl}$ TAC | Tukey one-way ANOVA | -3.488 | -6.160 to -0.8151 | **0.0049** |
| $Rhoa^{fl/fl}$ Sham vs $Rhoa^{\beta MHC\text{-}cre}$ Sham | Tukey one-way ANOVA | -0.1035 | -2.867 to 2.660 | >0.9999 |
| $Rhoa^{fl/fl}$ Sham vs $Rhoa^{\beta MHC\text{-}cre}$ TAC | Tukey one-way ANOVA | -6.244 | -8.846 to -3.642 | **<0.0001** |
| $Rhoa^{fl/fl}$ TAC vs $Rhoa^{\beta MHC\text{-}cre}$ Sham | Tukey one-way ANOVA | 3.384 | 0.8449 to 5.923 | **0.0039** |

| | | | | |
|---|---|---|---|---|
| *Rhoa*$^{fl/fl}$ TAC vs *Rhoa*$^{βMHC-cre}$ TAC | Tukey one-way ANOVA | -2.756 | -5.119 to -0.3943 | **0.0147** |
| *Rhoa*$^{βMHC-cre}$ Sham vs *Rhoa*$^{βMHC-cre}$ TAC | Tukey one-way ANOVA | -6.140 | -8.605 to -3.676 | **<0.0001** |
| **Panel C — CSA µm² (12-wk TAC)** | | | | |
| **Comparison** | **Test** | **Mean Diff.** | **95% CI** | **Adjusted P** |
| Omnibus one-way ANOVA | One-way ANOVA | F(5,31)=50.00 | — | **<0.0001** |
| WT$^{βMHC-cre}$ Sham vs WT$^{βMHC-cre}$ TAC | Tukey one-way ANOVA | -111.6 | -154.1 to -69.00 | **<0.0001** |
| WT$^{βMHC-cre}$ Sham vs *Rhoa*$^{fl/fl}$ Sham | Tukey one-way ANOVA | -12.44 | -57.08 to 32.19 | 0.9562 |
| WT$^{βMHC-cre}$ Sham vs *Rhoa*$^{fl/fl}$ TAC | Tukey one-way ANOVA | -104.9 | -145.9 to -63.90 | **<0.0001** |
| WT$^{βMHC-cre}$ Sham vs *Rhoa*$^{βMHC-cre}$ Sham | Tukey one-way ANOVA | -5.668 | -50.30 to 38.97 | 0.9988 |
| WT$^{βMHC-cre}$ Sham vs *Rhoa*$^{βMHC-cre}$ TAC | Tukey one-way ANOVA | -158.5 | -198.3 to -118.7 | **<0.0001** |
| WT$^{βMHC-cre}$ TAC vs *Rhoa*$^{fl/fl}$ Sham | Tukey one-way ANOVA | 99.12 | 54.48 to 143.8 | **<0.0001** |
| WT$^{βMHC-cre}$ TAC vs *Rhoa*$^{fl/fl}$ TAC | Tukey one-way ANOVA | 6.652 | -34.36 to 47.66 | 0.9961 |
| WT$^{βMHC-cre}$ TAC vs *Rhoa*$^{βMHC-cre}$ Sham | Tukey one-way ANOVA | 105.9 | 61.26 to 150.5 | **<0.0001** |
| WT$^{βMHC-cre}$ TAC vs *Rhoa*$^{βMHC-cre}$ TAC | Tukey one-way ANOVA | -46.97 | -86.79 to -7.164 | **0.0134** |
| *Rhoa*$^{fl/fl}$ Sham vs *Rhoa*$^{fl/fl}$ TAC | Tukey one-way ANOVA | -92.46 | -135.6 to -49.30 | **<0.0001** |
| *Rhoa*$^{fl/fl}$ Sham vs *Rhoa*$^{βMHC-cre}$ Sham | Tukey one-way ANOVA | 6.775 | -39.85 to 53.40 | 0.9977 |

| | | | | |
|---|---|---|---|---|
| *Rhoa*$^{fl/fl}$ Sham vs *Rhoa*$^{\beta MHC\text{-}cre}$ TAC | Tukey one-way ANOVA | -146.1 | -188.1 to -104.1 | **<0.0001** |
| *Rhoa*$^{fl/fl}$ TAC vs *Rhoa*$^{\beta MHC\text{-}cre}$ Sham | Tukey one-way ANOVA | 99.24 | 56.08 to 142.4 | **<0.0001** |
| *Rhoa*$^{fl/fl}$ TAC vs *Rhoa*$^{\beta MHC\text{-}cre}$ TAC | Tukey one-way ANOVA | -53.63 | -91.78 to -15.48 | **0.0022** |
| *Rhoa*$^{\beta MHC\text{-}cre}$ Sham vs *Rhoa*$^{\beta MHC\text{-}cre}$ TAC | Tukey one-way ANOVA | -152.9 | -194.9 to -110.8 | **<0.0001** |
| **Panel D — FS % (12-wk TAC)** | | | | |
| **Comparison** | **Test** | **Mean Diff.** | **95% CI** | **Adjusted P** |
| Omnibus one-way ANOVA | One-way ANOVA | F(5,30)=25.59 | — | **<0.0001** |
| WT$^{\beta MHC\text{-}cre}$ Sham vs WT$^{\beta MHC\text{-}cre}$ TAC | Tukey one-way ANOVA | 5.817 | 1.105 to 10.53 | **0.0088** |
| WT$^{\beta MHC\text{-}cre}$ Sham vs *Rhoa*$^{fl/fl}$ Sham | Tukey one-way ANOVA | 2.100 | -2.821 to 7.021 | 0.7838 |
| WT$^{\beta MHC\text{-}cre}$ Sham vs *Rhoa*$^{fl/fl}$ TAC | Tukey one-way ANOVA | 6.833 | 2.122 to 11.54 | **0.0016** |
| WT$^{\beta MHC\text{-}cre}$ Sham vs *Rhoa*$^{\beta MHC\text{-}cre}$ Sham | Tukey one-way ANOVA | -1.890 | -6.602 to 2.822 | 0.8237 |
| WT$^{\beta MHC\text{-}cre}$ Sham vs *Rhoa*$^{\beta MHC\text{-}cre}$ TAC | Tukey one-way ANOVA | 11.75 | 7.314 to 16.19 | **<0.0001** |
| WT$^{\beta MHC\text{-}cre}$ TAC vs *Rhoa*$^{fl/fl}$ Sham | Tukey one-way ANOVA | -3.717 | -8.428 to 0.9950 | 0.1886 |
| WT$^{\beta MHC\text{-}cre}$ TAC vs *Rhoa*$^{fl/fl}$ TAC | Tukey one-way ANOVA | 1.017 | -3.476 to 5.509 | 0.9819 |
| WT$^{\beta MHC\text{-}cre}$ TAC vs *Rhoa*$^{\beta MHC\text{-}cre}$ Sham | Tukey one-way ANOVA | -7.707 | -12.20 to -3.214 | **0.0002** |
| WT$^{\beta MHC\text{-}cre}$ TAC vs *Rhoa*$^{\beta MHC\text{-}cre}$ TAC | Tukey one-way ANOVA | 5.933 | 1.731 to 10.14 | **0.0021** |

| | | | | |
|---|---|---|---|---|
| *Rhoa*$^{fl/fl}$ Sham vs *Rhoa*$^{fl/fl}$ TAC | Tukey one-way ANOVA | 4.733 | 0.02168 to 9.445 | **0.0484** |
| *Rhoa*$^{fl/fl}$ Sham vs *Rhoa*$^{\beta MHC\text{-}cre}$ Sham | Tukey one-way ANOVA | -3.990 | -8.702 to 0.7217 | 0.1348 |
| *Rhoa*$^{fl/fl}$ Sham vs *Rhoa*$^{\beta MHC\text{-}cre}$ TAC | Tukey one-way ANOVA | 9.650 | 5.214 to 14.09 | **<0.0001** |
| *Rhoa*$^{fl/fl}$ TAC vs *Rhoa*$^{\beta MHC\text{-}cre}$ Sham | Tukey one-way ANOVA | -8.723 | -13.22 to -4.231 | **<0.0001** |
| *Rhoa*$^{fl/fl}$ TAC vs *Rhoa*$^{\beta MHC\text{-}cre}$ TAC | Tukey one-way ANOVA | 4.917 | 0.7144 to 9.119 | **0.0145** |
| *Rhoa*$^{\beta MHC\text{-}cre}$ Sham vs *Rhoa*$^{\beta MHC\text{-}cre}$ TAC | Tukey one-way ANOVA | 13.64 | 9.438 to 17.84 | **<0.0001** |
| **Panel E — LW/BW (12-wk TAC)** | | | | |
| **Comparison** | **Test** | **Mean Diff.** | **95% CI** | **Adjusted P** |
| Omnibus one-way ANOVA | One-way ANOVA | F(5,32)=8.981 | — | **<0.0001** |
| WT$^{\beta MHC\text{-}cre}$ Sham vs WT$^{\beta MHC\text{-}cre}$ TAC | Tukey one-way ANOVA | -3.334 | -10.40 to 3.730 | 0.7092 |
| WT$^{\beta MHC\text{-}cre}$ Sham vs *Rhoa*$^{fl/fl}$ Sham | Tukey one-way ANOVA | 0.4336 | -6.975 to 7.842 | >0.9999 |
| WT$^{\beta MHC\text{-}cre}$ Sham vs *Rhoa*$^{fl/fl}$ TAC | Tukey one-way ANOVA | -3.354 | -10.16 to 3.453 | 0.6712 |
| WT$^{\beta MHC\text{-}cre}$ Sham vs *Rhoa*$^{\beta MHC\text{-}cre}$ Sham | Tukey one-way ANOVA | -0.2253 | -7.289 to 6.838 | >0.9999 |
| WT$^{\beta MHC\text{-}cre}$ Sham vs *Rhoa*$^{\beta MHC\text{-}cre}$ TAC | Tukey one-way ANOVA | -11.54 | -18.15 to -4.932 | **0.0001** |
| WT$^{\beta MHC\text{-}cre}$ TAC vs *Rhoa*$^{fl/fl}$ Sham | Tukey one-way ANOVA | 3.768 | -3.641 to 11.18 | 0.6417 |
| WT$^{\beta MHC\text{-}cre}$ TAC vs *Rhoa*$^{fl/fl}$ TAC | Tukey one-way ANOVA | -0.02010 | -6.827 to 6.786 | >0.9999 |

| | | | | |
|---|---|---|---|---|
| WT$^{βMHC-cre}$ TAC vs *Rhoa*$^{βMHC-cre}$ Sham | Tukey one-way ANOVA | 3.109 | -3.955 to 10.17 | 0.7648 |
| WT$^{βMHC-cre}$ TAC vs *Rhoa*$^{βMHC-cre}$ TAC | Tukey one-way ANOVA | -8.206 | -14.81 to -1.598 | **0.0082** |
| *Rhoa*$^{fl/fl}$ Sham vs *Rhoa*$^{fl/fl}$ TAC | Tukey one-way ANOVA | -3.788 | -10.95 to 3.376 | 0.6035 |
| *Rhoa*$^{fl/fl}$ Sham vs *Rhoa*$^{βMHC-cre}$ Sham | Tukey one-way ANOVA | -0.6590 | -8.067 to 6.749 | 0.9998 |
| *Rhoa*$^{fl/fl}$ Sham vs *Rhoa*$^{βMHC-cre}$ TAC | Tukey one-way ANOVA | -11.97 | -18.95 to -4.999 | **0.0002** |
| *Rhoa*$^{fl/fl}$ TAC vs *Rhoa*$^{βMHC-cre}$ Sham | Tukey one-way ANOVA | 3.129 | -3.678 to 9.935 | 0.7311 |
| *Rhoa*$^{fl/fl}$ TAC vs *Rhoa*$^{βMHC-cre}$ TAC | Tukey one-way ANOVA | -8.186 | -14.52 to -1.854 | **0.0054** |
| *Rhoa*$^{βMHC-cre}$ Sham vs *Rhoa*$^{βMHC-cre}$ TAC | Tukey one-way ANOVA | -11.31 | -17.92 to -4.707 | **0.0002** |
| **Panel F — Fibrosis % (12-wk TAC)** | | | | |
| **Comparison** | **Test** | **Mean Diff.** | **95% CI** | **Adjusted P** |
| Omnibus one-way ANOVA | One-way ANOVA | F(5,22)=15.31 | — | **<0.0001** |
| WT$^{βMHC-cre}$ Sham vs WT$^{βMHC-cre}$ TAC | Tukey one-way ANOVA | -5.943 | -9.647 to -2.239 | **0.0007** |
| WT$^{βMHC-cre}$ Sham vs *Rhoa*$^{fl/fl}$ Sham | Tukey one-way ANOVA | -0.5260 | -4.018 to 2.966 | 0.9968 |
| WT$^{βMHC-cre}$ Sham vs *Rhoa*$^{fl/fl}$ TAC | Tukey one-way ANOVA | -6.583 | -10.29 to -2.879 | **0.0002** |
| WT$^{βMHC-cre}$ Sham vs *Rhoa*$^{βMHC-cre}$ Sham | Tukey one-way ANOVA | -0.4393 | -3.931 to 3.053 | 0.9986 |
| WT$^{βMHC-cre}$ Sham vs *Rhoa*$^{βMHC-cre}$ TAC | Tukey one-way ANOVA | -6.041 | -9.533 to -2.549 | **0.0003** |

| | | | | |
|---|---|---|---|---|
| WT$^{βMHC-cre}$ TAC vs *Rhoa*$^{fl/fl}$ Sham | Tukey one-way ANOVA | 5.417 | 1.713 to 9.121 | **0.0019** |
| WT$^{βMHC-cre}$ TAC vs *Rhoa*$^{fl/fl}$ TAC | Tukey one-way ANOVA | -0.6398 | -4.544 to 3.265 | 0.9952 |
| WT$^{βMHC-cre}$ TAC vs *Rhoa*$^{βMHC-cre}$ Sham | Tukey one-way ANOVA | 5.503 | 1.800 to 9.207 | **0.0016** |
| WT$^{βMHC-cre}$ TAC vs *Rhoa*$^{βMHC-cre}$ TAC | Tukey one-way ANOVA | -0.09821 | -3.802 to 3.606 | >0.9999 |
| *Rhoa*$^{fl/fl}$ Sham vs *Rhoa*$^{fl/fl}$ TAC | Tukey one-way ANOVA | -6.057 | -9.760 to -2.353 | **0.0005** |
| *Rhoa*$^{fl/fl}$ Sham vs *Rhoa*$^{βMHC-cre}$ Sham | Tukey one-way ANOVA | 0.08673 | -3.405 to 3.579 | >0.9999 |
| *Rhoa*$^{fl/fl}$ Sham vs *Rhoa*$^{βMHC-cre}$ TAC | Tukey one-way ANOVA | -5.515 | -9.007 to -2.023 | **0.0008** |
| *Rhoa*$^{fl/fl}$ TAC vs *Rhoa*$^{βMHC-cre}$ Sham | Tukey one-way ANOVA | 6.143 | 2.439 to 9.847 | **0.0004** |
| *Rhoa*$^{fl/fl}$ TAC vs *Rhoa*$^{βMHC-cre}$ TAC | Tukey one-way ANOVA | 0.5416 | -3.162 to 4.246 | 0.9972 |
| *Rhoa*$^{βMHC-cre}$ Sham vs *Rhoa*$^{βMHC-cre}$ TAC | Tukey one-way ANOVA | -5.602 | -9.094 to -2.110 | **0.0007** |

**Supplementary Table 5. Statistical comparisons for Figure 5.**

| **Panel B — HW/BW (Ang/PE)** | | | | |
|---|---|---|---|---|
| **Comparison** | **Test** | **Mean Diff.** | **95% CI** | **Adjusted P** |
| Omnibus one-way ANOVA | One-way ANOVA | F(9,72)=22.41 | — | **<0.0001** |
| WT$^{\beta MHC\text{-}cre}$ Veh vs WT$^{\beta MHC\text{-}cre}$ Ang/PE | Tukey one-way ANOVA | -1.095 | -2.412 to 0.2231 | 0.1887 |
| WT$^{\beta MHC\text{-}cre}$ Veh vs *Rhoa*$^{fl/fl}$ Veh | Tukey one-way ANOVA | 0.02341 | -1.332 to 1.379 | >0.9999 |
| WT$^{\beta MHC\text{-}cre}$ Veh vs *Rhoa*$^{fl/fl}$ Ang/PE | Tukey one-way ANOVA | -0.929 | -2.189 to 0.3310 | 0.3372 |
| WT$^{\beta MHC\text{-}cre}$ Veh vs *Rhoa*$^{\beta MHC\text{-}cre}$ Veh | Tukey one-way ANOVA | -0.05819 | -1.462 to 1.345 | >0.9999 |
| WT$^{\beta MHC\text{-}cre}$ Veh vs *Rhoa*$^{\beta MHC\text{-}cre}$ Ang/PE | Tukey one-way ANOVA | -1.668 | -2.886 to -0.4492 | **0.0011** |
| WT$^{\beta MHC\text{-}cre}$ Veh vs *Rhoa*$^{fl/fl}$;*Rhoc*$^{-/-}$ Veh | Tukey one-way ANOVA | 0.1156 | -1.349 to 1.580 | >0.9999 |
| WT$^{\beta MHC\text{-}cre}$ Veh vs *Rhoa*$^{fl/fl}$;*Rhoc*$^{-/-}$ Ang/PE | Tukey one-way ANOVA | -2.206 | -3.610 to -0.8029 | **<0.0001** |
| WT$^{\beta MHC\text{-}cre}$ Veh vs *Rhoa*$^{\beta MHC\text{-}cre}$;*Rhoc*$^{-/-}$ Veh | Tukey one-way ANOVA | -0.04684 | -1.511 to 1.418 | >0.9999 |
| WT$^{\beta MHC\text{-}cre}$ Veh vs *Rhoa*$^{\beta MHC\text{-}cre}$;*Rhoc*$^{-/-}$ Ang/PE | Tukey one-way ANOVA | -4.547 | -5.951 to -3.144 | **<0.0001** |
| WT$^{\beta MHC\text{-}cre}$ Ang/PE vs *Rhoa*$^{fl/fl}$ Veh | Tukey one-way ANOVA | 1.118 | -0.1997 to 2.436 | 0.1667 |
| WT$^{\beta MHC\text{-}cre}$ Ang/PE vs *Rhoa*$^{fl/fl}$ Ang/PE | Tukey one-way ANOVA | 0.1655 | -1.053 to 1.384 | >0.9999 |
| WT$^{\beta MHC\text{-}cre}$ Ang/PE vs *Rhoa*$^{\beta MHC\text{-}cre}$ Veh | Tukey one-way ANOVA | 1.036 | -0.3302 to 2.403 | 0.2989 |
| WT$^{\beta MHC\text{-}cre}$ Ang/PE vs *Rhoa*$^{\beta MHC\text{-}cre}$ Ang/PE | Tukey one-way ANOVA | -0.5732 | -1.749 to 0.6027 | 0.8479 |

| | | | | |
|---|---|---|---|---|
| WT$^{βMHC-cre}$ Ang/PE vs *Rhoa*$^{fl/fl}$;*Rhoc*$^{-/-}$ Veh | Tukey one-way ANOVA | 1.21 | -0.2191 to 2.639 | 0.1688 |
| WT$^{βMHC-cre}$ Ang/PE vs *Rhoa*$^{fl/fl}$;*Rhoc*$^{-/-}$ Ang/PE | Tukey one-way ANOVA | -1.112 | -2.478 to 0.2548 | 0.2117 |
| WT$^{βMHC-cre}$ Ang/PE vs *Rhoa*$^{βMHC-cre}$;*Rhoc*$^{-/-}$ Veh | Tukey one-way ANOVA | 1.048 | -0.3815 to 2.477 | 0.3452 |
| WT$^{βMHC-cre}$ Ang/PE vs *Rhoa*$^{βMHC-cre}$;*Rhoc*$^{-/-}$ Ang/PE | Tukey one-way ANOVA | -3.453 | -4.819 to - 2.086 | **<0.0001** |
| *Rhoa*$^{fl/fl}$ Veh vs *Rhoa*$^{fl/fl}$ Ang/PE | Tukey one-way ANOVA | -0.9524 | -2.212 to 0.3076 | 0.3033 |
| *Rhoa*$^{fl/fl}$ Veh vs *Rhoa*$^{βMHC-cre}$ Veh | Tukey one-way ANOVA | -0.0816 | -1.485 to 1.322 | >0.9999 |
| *Rhoa*$^{fl/fl}$ Veh vs *Rhoa*$^{βMHC-cre}$ Ang/PE | Tukey one-way ANOVA | -1.691 | -2.910 to - 0.4726 | **0.0009** |
| *Rhoa*$^{fl/fl}$ Veh vs *Rhoa*$^{fl/fl}$;*Rhoc*$^{-/-}$ Veh | Tukey one-way ANOVA | 0.09215 | -1.372 to 1.557 | >0.9999 |
| *Rhoa*$^{fl/fl}$ Veh vs *Rhoa*$^{fl/fl}$;*Rhoc*$^{-/-}$ Ang/PE | Tukey one-way ANOVA | -2.23 | -3.633 to - 0.8263 | **<0.0001** |
| *Rhoa*$^{fl/fl}$ Veh vs *Rhoa*$^{βMHC-cre}$;*Rhoc*$^{-/-}$ Veh | Tukey one-way ANOVA | -0.07025 | -1.535 to 1.394 | >0.9999 |
| *Rhoa*$^{fl/fl}$ Veh vs *Rhoa*$^{βMHC-cre}$;*Rhoc*$^{-/-}$ Ang/PE | Tukey one-way ANOVA | -4.571 | -5.974 to - 3.167 | **<0.0001** |
| *Rhoa*$^{fl/fl}$ Ang/PE vs *Rhoa*$^{βMHC-cre}$ Veh | Tukey one-way ANOVA | 0.8708 | -0.4402 to 2.182 | 0.4872 |
| *Rhoa*$^{fl/fl}$ Ang/PE vs *Rhoa*$^{βMHC-cre}$ Ang/PE | Tukey one-way ANOVA | -0.7387 | -1.850 to 0.3722 | 0.4857 |
| *Rhoa*$^{fl/fl}$ Ang/PE vs *Rhoa*$^{fl/fl}$;*Rhoc*$^{-/-}$ Veh | Tukey one-way ANOVA | 1.045 | -0.3316 to 2.421 | 0.2978 |
| *Rhoa*$^{fl/fl}$ Ang/PE vs *Rhoa*$^{fl/fl}$;*Rhoc*$^{-/-}$ Ang/PE | Tukey one-way ANOVA | -1.277 | -2.588 to 0.03379 | 0.0625 |

| | | | | |
|---|---|---|---|---|
| *Rhoa*$^{fl/fl}$ Ang/PE vs *Rhoa*$^{βMHC-cre}$;*Rhoc*$^{-/-}$ Veh | Tukey one-way ANOVA | 0.8822 | -0.4940 to 2.258 | 0.5385 |
| *Rhoa*$^{fl/fl}$ Ang/PE vs *Rhoa*$^{βMHC-cre}$;*Rhoc*$^{-/-}$ Ang/PE | Tukey one-way ANOVA | -3.618 | -4.929 to - 2.307 | **<0.0001** |
| *Rhoa*$^{βMHC-cre}$ Veh vs *Rhoa*$^{βMHC-cre}$ Ang/PE | Tukey one-way ANOVA | -1.609 | -2.881 to - 0.3383 | **0.0036** |
| *Rhoa*$^{βMHC-cre}$ Veh vs *Rhoa*$^{fl/fl}$;*Rhoc*$^{-/-}$ Veh | Tukey one-way ANOVA | 0.1737 | -1.335 to 1.682 | >0.9999 |
| *Rhoa*$^{βMHC-cre}$ Veh vs *Rhoa*$^{fl/fl}$;*Rhoc*$^{-/-}$ Ang/PE | Tukey one-way ANOVA | -2.148 | -3.598 to - 0.6987 | **0.0003** |
| *Rhoa*$^{βMHC-cre}$ Veh vs *Rhoa*$^{βMHC-cre}$;*Rhoc*$^{-/-}$ Veh | Tukey one-way ANOVA | 0.01135 | -1.497 to 1.520 | >0.9999 |
| *Rhoa*$^{βMHC-cre}$ Veh vs *Rhoa*$^{βMHC-cre}$;*Rhoc*$^{-/-}$ Ang/PE | Tukey one-way ANOVA | -4.489 | -5.939 to - 3.040 | **<0.0001** |
| *Rhoa*$^{βMHC-cre}$ Ang/PE vs *Rhoa*$^{fl/fl}$;*Rhoc*$^{-/-}$ Veh | Tukey one-way ANOVA | 1.783 | 0.4449 to 3.122 | **0.0017** |
| *Rhoa*$^{βMHC-cre}$ Ang/PE vs *Rhoa*$^{fl/fl}$;*Rhoc*$^{-/-}$ Ang/PE | Tukey one-way ANOVA | -0.5386 | -1.810 to 0.7326 | 0.9284 |
| *Rhoa*$^{βMHC-cre}$ Ang/PE vs *Rhoa*$^{βMHC-cre}$;*Rhoc*$^{-/-}$ Veh | Tukey one-way ANOVA | 1.621 | 0.2825 to 2.959 | **0.0064** |
| *Rhoa*$^{βMHC-cre}$ Ang/PE vs *Rhoa*$^{βMHC-cre}$;*Rhoc*$^{-/-}$ Ang/PE | Tukey one-way ANOVA | -2.88 | -4.151 to - 1.608 | **<0.0001** |
| *Rhoa*$^{fl/fl}$;*Rhoc*$^{-/-}$ Veh vs *Rhoa*$^{fl/fl}$;*Rhoc*$^{-/-}$ Ang/PE | Tukey one-way ANOVA | -2.322 | -3.830 to - 0.8132 | **0.0001** |
| *Rhoa*$^{fl/fl}$;*Rhoc*$^{-/-}$ Veh vs *Rhoa*$^{βMHC-cre}$;*Rhoc*$^{-/-}$ Veh | Tukey one-way ANOVA | -0.1624 | -1.728 to 1.403 | >0.9999 |
| *Rhoa*$^{fl/fl}$;*Rhoc*$^{-/-}$ Veh vs *Rhoa*$^{βMHC-cre}$;*Rhoc*$^{-/-}$ Ang/PE | Tukey one-way ANOVA | -4.663 | -6.171 to - 3.154 | **<0.0001** |
| *Rhoa*$^{fl/fl}$;*Rhoc*$^{-/-}$ Ang/PE vs *Rhoa*$^{βMHC-cre}$;*Rhoc*$^{-/-}$ Veh | Tukey one-way ANOVA | 2.159 | 0.6508 to 3.668 | **0.0005** |

| | | | | |
|---|---|---|---|---|
| *Rhoa*$^{fl/fl}$;*Rhoc*$^{-/-}$ Ang/PE vs *Rhoa*$^{βMHC-cre}$;*Rhoc*$^{-/-}$ Ang/PE | Tukey one-way ANOVA | -2.341 | -3.790 to -0.8916 | **<0.0001** |
| *Rhoa*$^{βMHC-cre}$;*Rhoc*$^{-/-}$ Veh vs *Rhoa*$^{βMHC-cre}$;*Rhoc*$^{-/-}$ Ang/PE | Tukey one-way ANOVA | -4.5 | -6.009 to -2.992 | **<0.0001** |
| **Panel C — CSA µm² (Ang/PE)** | | | | |
| **Comparison** | **Test** | **Mean Diff.** | **95% CI** | **Adjusted P** |
| Omnibus one-way ANOVA | One-way ANOVA | $F(9,66)=19.17$ | — | **<0.0001** |
| WT$^{βMHC-cre}$ Veh vs WT$^{βMHC-cre}$ Ang/PE | Tukey one-way ANOVA | -21.17 | -40.25 to -2.086 | **0.0183** |
| WT$^{βMHC-cre}$ Veh vs *Rhoa*$^{fl/fl}$ Veh | Tukey one-way ANOVA | -0.8652 | -21.93 to 20.20 | >0.9999 |
| WT$^{βMHC-cre}$ Veh vs *Rhoa*$^{fl/fl}$ Ang/PE | Tukey one-way ANOVA | -21.87 | -40.96 to -2.793 | **0.0127** |
| WT$^{βMHC-cre}$ Veh vs *Rhoa*$^{βMHC-cre}$ Veh | Tukey one-way ANOVA | -1.221 | -22.29 to 19.84 | >0.9999 |
| WT$^{βMHC-cre}$ Veh vs *Rhoa*$^{βMHC-cre}$ Ang/PE | Tukey one-way ANOVA | -23.99 | -42.65 to -5.330 | **0.003** |
| WT$^{βMHC-cre}$ Veh vs *Rhoa*$^{fl/fl}$;*Rhoc*$^{-/-}$ Veh | Tukey one-way ANOVA | 9.657 | -10.58 to 29.90 | 0.8609 |
| WT$^{βMHC-cre}$ Veh vs *Rhoa*$^{fl/fl}$;*Rhoc*$^{-/-}$ Ang/PE | Tukey one-way ANOVA | -23.53 | -43.76 to -3.286 | **0.0108** |
| WT$^{βMHC-cre}$ Veh vs *Rhoa*$^{βMHC-cre}$;*Rhoc*$^{-/-}$ Veh | Tukey one-way ANOVA | -1.332 | -20.93 to 18.26 | >0.9999 |
| WT$^{βMHC-cre}$ Veh vs *Rhoa*$^{βMHC-cre}$;*Rhoc*$^{-/-}$ Ang/PE | Tukey one-way ANOVA | -55.21 | -75.45 to -34.97 | **<0.0001** |
| WT$^{βMHC-cre}$ Ang/PE vs *Rhoa*$^{fl/fl}$ Veh | Tukey one-way ANOVA | 20.3 | 0.3460 to 40.26 | **0.0429** |
| WT$^{βMHC-cre}$ Ang/PE vs *Rhoa*$^{fl/fl}$ Ang/PE | Tukey one-way ANOVA | -0.7074 | -18.56 to 17.14 | >0.9999 |
| WT$^{βMHC-cre}$ Ang/PE vs *Rhoa*$^{βMHC-cre}$ Veh | Tukey one-way ANOVA | 19.95 | -0.009580 to 39.90 | 0.0502 |

| | | | | |
|---|---|---|---|---|
| WT$^{βMHC-cre}$ Ang/PE vs *Rhoa*$^{βMHC-cre}$ Ang/PE | Tukey one-way ANOVA | -2.822 | -20.22 to 14.57 | >0.9999 |
| WT$^{βMHC-cre}$ Ang/PE vs *Rhoa*$^{fl/fl}$;*Rhoc*$^{-/-}$ Veh | Tukey one-way ANOVA | 30.82 | 11.74 to 49.91 | **<0.0001** |
| WT$^{βMHC-cre}$ Ang/PE vs *Rhoa*$^{fl/fl}$;*Rhoc*$^{-/-}$ Ang/PE | Tukey one-way ANOVA | -2.358 | -21.44 to 16.72 | >0.9999 |
| WT$^{βMHC-cre}$ Ang/PE vs *Rhoa*$^{βMHC-cre}$;*Rhoc*$^{-/-}$ Veh | Tukey one-way ANOVA | 19.83 | 1.437 to 38.23 | **0.0247** |
| WT$^{βMHC-cre}$ Ang/PE vs *Rhoa*$^{βMHC-cre}$;*Rhoc*$^{-/-}$ Ang/PE | Tukey one-way ANOVA | -34.05 | -53.13 to -14.97 | **<0.0001** |
| *Rhoa*$^{fl/fl}$ Veh vs *Rhoa*$^{fl/fl}$ Ang/PE | Tukey one-way ANOVA | -21.01 | -40.96 to -1.053 | **0.0312** |
| *Rhoa*$^{fl/fl}$ Veh vs *Rhoa*$^{βMHC-cre}$ Veh | Tukey one-way ANOVA | -0.3556 | -22.22 to 21.50 | >0.9999 |
| *Rhoa*$^{fl/fl}$ Veh vs *Rhoa*$^{βMHC-cre}$ Ang/PE | Tukey one-way ANOVA | -23.12 | -42.68 to -3.572 | **0.0088** |
| *Rhoa*$^{fl/fl}$ Veh vs *Rhoa*$^{fl/fl}$;*Rhoc*$^{-/-}$ Veh | Tukey one-way ANOVA | 10.52 | -10.54 to 31.59 | 0.8258 |
| *Rhoa*$^{fl/fl}$ Veh vs *Rhoa*$^{fl/fl}$;*Rhoc*$^{-/-}$ Ang/PE | Tukey one-way ANOVA | -22.66 | -43.72 to -1.595 | **0.0252** |
| *Rhoa*$^{fl/fl}$ Veh vs *Rhoa*$^{βMHC-cre}$;*Rhoc*$^{-/-}$ Veh | Tukey one-way ANOVA | -0.4669 | -20.92 to 19.98 | >0.9999 |
| *Rhoa*$^{fl/fl}$ Veh vs *Rhoa*$^{βMHC-cre}$;*Rhoc*$^{-/-}$ Ang/PE | Tukey one-way ANOVA | -54.35 | -75.41 to -33.28 | **<0.0001** |
| *Rhoa*$^{fl/fl}$ Ang/PE vs *Rhoa*$^{βMHC-cre}$ Veh | Tukey one-way ANOVA | 20.65 | 0.6978 to 40.61 | **0.0367** |
| *Rhoa*$^{fl/fl}$ Ang/PE vs *Rhoa*$^{βMHC-cre}$ Ang/PE | Tukey one-way ANOVA | -2.115 | -19.51 to 15.28 | >0.9999 |
| *Rhoa*$^{fl/fl}$ Ang/PE vs *Rhoa*$^{fl/fl}$;*Rhoc*$^{-/-}$ Veh | Tukey one-way ANOVA | 31.53 | 12.45 to 50.61 | **<0.0001** |

| | | | | |
|---|---|---|---|---|
| *Rhoa*$^{fl/fl}$ Ang/PE vs *Rhoa*$^{fl/fl}$;*Rhoc*$^{-/-}$ Ang/PE | Tukey one-way ANOVA | -1.651 | -20.73 to 17.43 | >0.9999 |
| *Rhoa*$^{fl/fl}$ Ang/PE vs *Rhoa*$^{βMHC-cre}$;*Rhoc*$^{-/-}$ Veh | Tukey one-way ANOVA | 20.54 | 2.144 to 38.94 | **0.0171** |
| *Rhoa*$^{fl/fl}$ Ang/PE vs *Rhoa*$^{βMHC-cre}$;*Rhoc*$^{-/-}$ Ang/PE | Tukey one-way ANOVA | -33.34 | -52.42 to -14.26 | **<0.0001** |
| *Rhoa*$^{βMHC-cre}$ Veh vs *Rhoa*$^{βMHC-cre}$ Ang/PE | Tukey one-way ANOVA | -22.77 | -42.32 to -3.216 | **0.0106** |
| *Rhoa*$^{βMHC-cre}$ Veh vs *Rhoa*$^{fl/fl}$;*Rhoc*$^{-/-}$ Veh | Tukey one-way ANOVA | 10.88 | -10.19 to 31.94 | 0.7965 |
| *Rhoa*$^{βMHC-cre}$ Veh vs *Rhoa*$^{fl/fl}$;*Rhoc*$^{-/-}$ Ang/PE | Tukey one-way ANOVA | -22.3 | -43.37 to -1.239 | **0.0295** |
| *Rhoa*$^{βMHC-cre}$ Veh vs *Rhoa*$^{βMHC-cre}$;*Rhoc*$^{-/-}$ Veh | Tukey one-way ANOVA | -0.1113 | -20.56 to 20.34 | >0.9999 |
| *Rhoa*$^{βMHC-cre}$ Veh vs *Rhoa*$^{βMHC-cre}$;*Rhoc*$^{-/-}$ Ang/PE | Tukey one-way ANOVA | -53.99 | -75.06 to -32.93 | **<0.0001** |
| *Rhoa*$^{βMHC-cre}$ Ang/PE vs *Rhoa*$^{fl/fl}$;*Rhoc*$^{-/-}$ Veh | Tukey one-way ANOVA | 33.65 | 14.99 to 52.31 | **<0.0001** |
| *Rhoa*$^{βMHC-cre}$ Ang/PE vs *Rhoa*$^{fl/fl}$;*Rhoc*$^{-/-}$ Ang/PE | Tukey one-way ANOVA | 0.4641 | -18.19 to 19.12 | >0.9999 |
| *Rhoa*$^{βMHC-cre}$ Ang/PE vs *Rhoa*$^{βMHC-cre}$;*Rhoc*$^{-/-}$ Veh | Tukey one-way ANOVA | 22.66 | 4.697 to 40.62 | **0.0039** |
| *Rhoa*$^{βMHC-cre}$ Ang/PE vs *Rhoa*$^{βMHC-cre}$;*Rhoc*$^{-/-}$ Ang/PE | Tukey one-way ANOVA | -31.22 | -49.88 to -12.57 | **<0.0001** |
| *Rhoa*$^{fl/fl}$;*Rhoc*$^{-/-}$ Veh vs *Rhoa*$^{fl/fl}$;*Rhoc*$^{-/-}$ Ang/PE | Tukey one-way ANOVA | -33.18 | -53.42 to -12.94 | **<0.0001** |
| *Rhoa*$^{fl/fl}$;*Rhoc*$^{-/-}$ Veh vs *Rhoa*$^{βMHC-cre}$;*Rhoc*$^{-/-}$ Veh | Tukey one-way ANOVA | -10.99 | -30.59 to 8.607 | 0.7102 |
| *Rhoa*$^{fl/fl}$;*Rhoc*$^{-/-}$ Veh vs *Rhoa*$^{βMHC-cre}$;*Rhoc*$^{-/-}$ Ang/PE | Tukey one-way ANOVA | -64.87 | -85.11 to -44.63 | **<0.0001** |

| | | | | |
|---|---|---|---|---|
| *Rhoa*$^{\text{fl/fl}}$;*Rhoc*$^{-/-}$ Ang/PE vs *Rhoa*$^{\beta\text{MHC-cre}}$;*Rhoc*$^{-/-}$ Veh | Tukey one-way ANOVA | 22.19 | 2.597 to 41.79 | **0.0146** |
| *Rhoa*$^{\text{fl/fl}}$;*Rhoc*$^{-/-}$ Ang/PE vs *Rhoa*$^{\beta\text{MHC-cre}}$;*Rhoc*$^{-/-}$ Ang/PE | Tukey one-way ANOVA | -31.69 | -51.93 to -11.45 | **0.0001** |
| *Rhoa*$^{\beta\text{MHC-cre}}$;*Rhoc*$^{-/-}$ Veh vs *Rhoa*$^{\beta\text{MHC-cre}}$;*Rhoc*$^{-/-}$ Ang/PE | Tukey one-way ANOVA | -53.88 | -73.48 to -34.29 | **<0.0001** |
| **Panel D — FS % (Ang/PE)** | | | | |
| **Comparison** | **Test** | **Mean Diff.** | **95% CI** | **Adjusted P** |
| Omnibus one-way ANOVA | One-way ANOVA | $F(9,75)=7.118$ | — | **<0.0001** |
| WT$^{\beta\text{MHC-cre}}$ Veh vs WT$^{\beta\text{MHC-cre}}$ Ang/PE | Tukey one-way ANOVA | -5.195 | -11.45 to 1.062 | 0.1897 |
| WT$^{\beta\text{MHC-cre}}$ Veh vs *Rhoa*$^{\text{fl/fl}}$ Veh | Tukey one-way ANOVA | 0.795 | -5.462 to 7.052 | >0.9999 |
| WT$^{\beta\text{MHC-cre}}$ Veh vs *Rhoa*$^{\text{fl/fl}}$ Ang/PE | Tukey one-way ANOVA | -3.757 | -9.886 to 2.372 | 0.6026 |
| WT$^{\beta\text{MHC-cre}}$ Veh vs *Rhoa*$^{\beta\text{MHC-cre}}$ Veh | Tukey one-way ANOVA | 0.7107 | -6.116 to 7.537 | >0.9999 |
| WT$^{\beta\text{MHC-cre}}$ Veh vs *Rhoa*$^{\beta\text{MHC-cre}}$ Ang/PE | Tukey one-way ANOVA | -2.375 | -8.395 to 3.645 | 0.9537 |
| WT$^{\beta\text{MHC-cre}}$ Veh vs *Rhoa*$^{\text{fl/fl}}$;*Rhoc*$^{-/-}$ Veh | Tukey one-way ANOVA | -1.075 | -7.902 to 5.752 | >0.9999 |
| WT$^{\beta\text{MHC-cre}}$ Veh vs *Rhoa*$^{\text{fl/fl}}$;*Rhoc*$^{-/-}$ Ang/PE | Tukey one-way ANOVA | 4.439 | -2.387 to 11.27 | 0.5194 |
| WT$^{\beta\text{MHC-cre}}$ Veh vs *Rhoa*$^{\beta\text{MHC-cre}}$;*Rhoc*$^{-/-}$ Veh | Tukey one-way ANOVA | -0.7417 | -7.865 to 6.382 | >0.9999 |
| WT$^{\beta\text{MHC-cre}}$ Veh vs *Rhoa*$^{\beta\text{MHC-cre}}$;*Rhoc*$^{-/-}$ Ang/PE | Tukey one-way ANOVA | 7.668 | 0.8413 to 14.49 | **0.0157** |
| WT$^{\beta\text{MHC-cre}}$ Ang/PE vs *Rhoa*$^{\text{fl/fl}}$ Veh | Tukey one-way ANOVA | 5.99 | 0.09121 to 11.89 | **0.0436** |
| WT$^{\beta\text{MHC-cre}}$ Ang/PE vs *Rhoa*$^{\text{fl/fl}}$ Ang/PE | Tukey one-way ANOVA | 1.438 | -4.325 to 7.201 | 0.9982 |

| | | | | |
|---|---|---|---|---|
| WT$^{\beta MHC\text{-}cre}$ Ang/PE vs *Rhoa*$^{\beta MHC\text{-}cre}$ Veh | Tukey one-way ANOVA | 5.906 | -0.5944 to 12.41 | 0.1069 |
| WT$^{\beta MHC\text{-}cre}$ Ang/PE vs *Rhoa*$^{\beta MHC\text{-}cre}$ Ang/PE | Tukey one-way ANOVA | 2.82 | -2.828 to 8.468 | 0.8301 |
| WT$^{\beta MHC\text{-}cre}$ Ang/PE vs *Rhoa*$^{fl/fl}$;*Rhoc*$^{-/-}$ Veh | Tukey one-way ANOVA | 4.12 | -2.380 to 10.62 | 0.5561 |
| WT$^{\beta MHC\text{-}cre}$ Ang/PE vs *Rhoa*$^{fl/fl}$;*Rhoc*$^{-/-}$ Ang/PE | Tukey one-way ANOVA | 9.634 | 3.134 to 16.13 | **0.0003** |
| WT$^{\beta MHC\text{-}cre}$ Ang/PE vs *Rhoa*$^{\beta MHC\text{-}cre}$;*Rhoc*$^{-/-}$ Veh | Tukey one-way ANOVA | 4.453 | -2.358 to 11.26 | 0.5116 |
| WT$^{\beta MHC\text{-}cre}$ Ang/PE vs *Rhoa*$^{\beta MHC\text{-}cre}$;*Rhoc*$^{-/-}$ Ang/PE | Tukey one-way ANOVA | 12.86 | 6.363 to 19.36 | **<0.0001** |
| *Rhoa*$^{fl/fl}$ Veh vs *Rhoa*$^{fl/fl}$ Ang/PE | Tukey one-way ANOVA | -4.552 | -10.31 to 1.211 | 0.2477 |
| *Rhoa*$^{fl/fl}$ Veh vs *Rhoa*$^{\beta MHC\text{-}cre}$ Veh | Tukey one-way ANOVA | -0.08429 | -6.584 to 6.416 | >0.9999 |
| *Rhoa*$^{fl/fl}$ Veh vs *Rhoa*$^{\beta MHC\text{-}cre}$ Ang/PE | Tukey one-way ANOVA | -3.17 | -8.818 to 2.478 | 0.7141 |
| *Rhoa*$^{fl/fl}$ Veh vs *Rhoa*$^{fl/fl}$;*Rhoc*$^{-/-}$ Veh | Tukey one-way ANOVA | -1.87 | -8.370 to 4.630 | 0.9946 |
| *Rhoa*$^{fl/fl}$ Veh vs *Rhoa*$^{fl/fl}$;*Rhoc*$^{-/-}$ Ang/PE | Tukey one-way ANOVA | 3.644 | -2.856 to 10.14 | 0.7155 |
| *Rhoa*$^{fl/fl}$ Veh vs *Rhoa*$^{\beta MHC\text{-}cre}$;*Rhoc*$^{-/-}$ Veh | Tukey one-way ANOVA | -1.537 | -8.348 to 5.275 | 0.9992 |
| *Rhoa*$^{fl/fl}$ Veh vs *Rhoa*$^{\beta MHC\text{-}cre}$;*Rhoc*$^{-/-}$ Ang/PE | Tukey one-way ANOVA | 6.873 | 0.3727 to 13.37 | **0.0297** |
| *Rhoa*$^{fl/fl}$ Ang/PE vs *Rhoa*$^{\beta MHC\text{-}cre}$ Veh | Tukey one-way ANOVA | 4.468 | -1.910 to 10.84 | 0.4112 |
| *Rhoa*$^{fl/fl}$ Ang/PE vs *Rhoa*$^{\beta MHC\text{-}cre}$ Ang/PE | Tukey one-way ANOVA | 1.382 | -4.124 to 6.888 | 0.9981 |

| | | | | |
|---|---|---|---|---|
| *Rhoa*$^{fl/fl}$ Ang/PE vs *Rhoa*$^{fl/fl}$;*Rhoc*$^{-/-}$ Veh | Tukey one-way ANOVA | 2.682 | -3.696 to 9.059 | 0.9321 |
| *Rhoa*$^{fl/fl}$ Ang/PE vs *Rhoa*$^{fl/fl}$;*Rhoc*$^{-/-}$ Ang/PE | Tukey one-way ANOVA | 8.196 | 1.819 to 14.57 | **0.0029** |
| *Rhoa*$^{fl/fl}$ Ang/PE vs *Rhoa*$^{\beta MHC\text{-}cre}$;*Rhoc*$^{-/-}$ Veh | Tukey one-way ANOVA | 3.015 | -3.679 to 9.709 | 0.9 |
| *Rhoa*$^{fl/fl}$ Ang/PE vs *Rhoa*$^{\beta MHC\text{-}cre}$;*Rhoc*$^{-/-}$ Ang/PE | Tukey one-way ANOVA | 11.42 | 5.047 to 17.80 | **<0.0001** |
| *Rhoa*$^{\beta MHC\text{-}cre}$ Veh vs *Rhoa*$^{\beta MHC\text{-}cre}$ Ang/PE | Tukey one-way ANOVA | -3.086 | -9.359 to 3.187 | 0.8421 |
| *Rhoa*$^{\beta MHC\text{-}cre}$ Veh vs *Rhoa*$^{fl/fl}$;*Rhoc*$^{-/-}$ Veh | Tukey one-way ANOVA | -1.786 | -8.836 to 5.265 | 0.9979 |
| *Rhoa*$^{\beta MHC\text{-}cre}$ Veh vs *Rhoa*$^{fl/fl}$;*Rhoc*$^{-/-}$ Ang/PE | Tukey one-way ANOVA | 3.729 | -3.322 to 10.78 | 0.7781 |
| *Rhoa*$^{\beta MHC\text{-}cre}$ Veh vs *Rhoa*$^{\beta MHC\text{-}cre}$;*Rhoc*$^{-/-}$ Veh | Tukey one-way ANOVA | -1.452 | -8.791 to 5.886 | 0.9997 |
| *Rhoa*$^{\beta MHC\text{-}cre}$ Veh vs *Rhoa*$^{\beta MHC\text{-}cre}$;*Rhoc*$^{-/-}$ Ang/PE | Tukey one-way ANOVA | 6.957 | -0.09326 to 14.01 | 0.0561 |
| *Rhoa*$^{\beta MHC\text{-}cre}$ Ang/PE vs *Rhoa*$^{fl/fl}$;*Rhoc*$^{-/-}$ Veh | Tukey one-way ANOVA | 1.3 | -4.973 to 7.573 | 0.9996 |
| *Rhoa*$^{\beta MHC\text{-}cre}$ Ang/PE vs *Rhoa*$^{fl/fl}$;*Rhoc*$^{-/-}$ Ang/PE | Tukey one-way ANOVA | 6.814 | 0.5411 to 13.09 | **0.0226** |
| *Rhoa*$^{\beta MHC\text{-}cre}$ Ang/PE vs *Rhoa*$^{\beta MHC\text{-}cre}$;*Rhoc*$^{-/-}$ Veh | Tukey one-way ANOVA | 1.633 | -4.962 to 8.228 | 0.9983 |
| *Rhoa*$^{\beta MHC\text{-}cre}$ Ang/PE vs *Rhoa*$^{\beta MHC\text{-}cre}$;*Rhoc*$^{-/-}$ Ang/PE | Tukey one-way ANOVA | 10.04 | 3.770 to 16.32 | **<0.0001** |
| *Rhoa*$^{fl/fl}$;*Rhoc*$^{-/-}$ Veh vs *Rhoa*$^{fl/fl}$;*Rhoc*$^{-/-}$ Ang/PE | Tukey one-way ANOVA | 5.514 | -1.536 to 12.56 | 0.2599 |
| *Rhoa*$^{fl/fl}$;*Rhoc*$^{-/-}$ Veh vs *Rhoa*$^{\beta MHC\text{-}cre}$;*Rhoc*$^{-/-}$ Veh | Tukey one-way ANOVA | 0.3333 | -7.005 to 7.672 | >0.9999 |

| | | | | |
|---|---|---|---|---|
| *Rhoa*$^{fl/fl}$;*Rhoc*$^{-/-}$ Veh vs *Rhoa*$^{βMHC-cre}$;*Rhoc*$^{-/-}$ Ang/PE | Tukey one-way ANOVA | 8.743 | 1.692 to 15.79 | **0.0047** |
| *Rhoa*$^{fl/fl}$;*Rhoc*$^{-/-}$ Ang/PE vs *Rhoa*$^{βMHC-cre}$;*Rhoc*$^{-/-}$ Veh | Tukey one-way ANOVA | -5.181 | -12.52 to 2.157 | 0.3999 |
| *Rhoa*$^{fl/fl}$;*Rhoc*$^{-/-}$ Ang/PE vs *Rhoa*$^{βMHC-cre}$;*Rhoc*$^{-/-}$ Ang/PE | Tukey one-way ANOVA | 3.229 | -3.822 to 10.28 | 0.8907 |
| *Rhoa*$^{βMHC-cre}$;*Rhoc*$^{-/-}$ Veh vs *Rhoa*$^{βMHC-cre}$;*Rhoc*$^{-/-}$ Ang/PE | Tukey one-way ANOVA | 8.41 | 1.071 to 15.75 | **0.0125** |
| **Panel E — LW/BW (Ang/PE)** | | | | |
| **Comparison** | **Test** | **Mean Diff.** | **95% CI** | **Adjusted P** |
| Omnibus one-way ANOVA | One-way ANOVA | F(9,72)=31.59 | — | **<0.0001** |
| WT$^{βMHC-cre}$ Veh vs WT$^{βMHC-cre}$ Ang/PE | Tukey one-way ANOVA | -1.073 | -2.897 to 0.7507 | 0.655 |
| WT$^{βMHC-cre}$ Veh vs *Rhoa*$^{fl/fl}$ Veh | Tukey one-way ANOVA | 0.3232 | -1.554 to 2.200 | >0.9999 |
| WT$^{βMHC-cre}$ Veh vs *Rhoa*$^{fl/fl}$ Ang/PE | Tukey one-way ANOVA | -0.9686 | -2.713 to 0.7757 | 0.7249 |
| WT$^{βMHC-cre}$ Veh vs *Rhoa*$^{βMHC-cre}$ Veh | Tukey one-way ANOVA | -0.1328 | -2.076 to 1.810 | >0.9999 |
| WT$^{βMHC-cre}$ Veh vs *Rhoa*$^{βMHC-cre}$ Ang/PE | Tukey one-way ANOVA | -1.847 | -3.534 to -0.1606 | **0.0208** |
| WT$^{βMHC-cre}$ Veh vs *Rhoa*$^{fl/fl}$;*Rhoc*$^{-/-}$ Veh | Tukey one-way ANOVA | 0.3352 | -1.692 to 2.363 | >0.9999 |
| WT$^{βMHC-cre}$ Veh vs *Rhoa*$^{fl/fl}$;*Rhoc*$^{-/-}$ Ang/PE | Tukey one-way ANOVA | -6.162 | -8.105 to -4.219 | **<0.0001** |
| WT$^{βMHC-cre}$ Veh vs *Rhoa*$^{βMHC-cre}$;*Rhoc*$^{-/-}$ Veh | Tukey one-way ANOVA | 0.04027 | -1.987 to 2.068 | >0.9999 |
| WT$^{βMHC-cre}$ Veh vs *Rhoa*$^{βMHC-cre}$;*Rhoc*$^{-/-}$ Ang/PE | Tukey one-way ANOVA | -5.731 | -7.674 to -3.788 | **<0.0001** |
| WT$^{βMHC-cre}$ Ang/PE vs *Rhoa*$^{fl/fl}$ Veh | Tukey one-way ANOVA | 1.397 | -0.4275 to 3.221 | 0.2863 |

| | | | | |
|---|---|---|---|---|
| WT$^{\beta MHC\text{-}cre}$ Ang/PE vs *Rhoa*$^{fl/fl}$ Ang/PE | Tukey one-way ANOVA | 0.1047 | -1.583 to 1.792 | >0.9999 |
| WT$^{\beta MHC\text{-}cre}$ Ang/PE vs *Rhoa*$^{\beta MHC\text{-}cre}$ Veh | Tukey one-way ANOVA | 0.9406 | -0.9512 to 2.832 | 0.8324 |
| WT$^{\beta MHC\text{-}cre}$ Ang/PE vs *Rhoa*$^{\beta MHC\text{-}cre}$ Ang/PE | Tukey one-way ANOVA | -0.7741 | -2.402 to 0.8537 | 0.8657 |
| WT$^{\beta MHC\text{-}cre}$ Ang/PE vs *Rhoa*$^{fl/fl}$;*Rhoc*$^{-/-}$ Veh | Tukey one-way ANOVA | 1.409 | -0.5699 to 3.387 | 0.3866 |
| WT$^{\beta MHC\text{-}cre}$ Ang/PE vs *Rhoa*$^{fl/fl}$;*Rhoc*$^{-/-}$ Ang/PE | Tukey one-way ANOVA | -5.088 | -6.980 to -3.197 | **<0.0001** |
| WT$^{\beta MHC\text{-}cre}$ Ang/PE vs *Rhoa*$^{\beta MHC\text{-}cre}$;*Rhoc*$^{-/-}$ Veh | Tukey one-way ANOVA | 1.114 | -0.8649 to 3.092 | 0.7094 |
| WT$^{\beta MHC\text{-}cre}$ Ang/PE vs *Rhoa*$^{\beta MHC\text{-}cre}$;*Rhoc*$^{-/-}$ Ang/PE | Tukey one-way ANOVA | -4.658 | -6.549 to -2.766 | **<0.0001** |
| *Rhoa*$^{fl/fl}$ Veh vs *Rhoa*$^{fl/fl}$ Ang/PE | Tukey one-way ANOVA | -1.292 | -3.036 to 0.4524 | 0.331 |
| *Rhoa*$^{fl/fl}$ Veh vs *Rhoa*$^{\beta MHC\text{-}cre}$ Veh | Tukey one-way ANOVA | -0.456 | -2.399 to 1.487 | 0.9988 |
| *Rhoa*$^{fl/fl}$ Veh vs *Rhoa*$^{\beta MHC\text{-}cre}$ Ang/PE | Tukey one-way ANOVA | -2.171 | -3.858 to -0.4838 | **0.0028** |
| *Rhoa*$^{fl/fl}$ Veh vs *Rhoa*$^{fl/fl}$;*Rhoc*$^{-/-}$ Veh | Tukey one-way ANOVA | 0.012 | -2.015 to 2.039 | >0.9999 |
| *Rhoa*$^{fl/fl}$ Veh vs *Rhoa*$^{fl/fl}$;*Rhoc*$^{-/-}$ Ang/PE | Tukey one-way ANOVA | -6.485 | -8.428 to -4.542 | **<0.0001** |
| *Rhoa*$^{fl/fl}$ Veh vs *Rhoa*$^{\beta MHC\text{-}cre}$;*Rhoc*$^{-/-}$ Veh | Tukey one-way ANOVA | -0.283 | -2.310 to 1.744 | >0.9999 |
| *Rhoa*$^{fl/fl}$ Veh vs *Rhoa*$^{\beta MHC\text{-}cre}$;*Rhoc*$^{-/-}$ Ang/PE | Tukey one-way ANOVA | -6.054 | -7.997 to -4.111 | **<0.0001** |
| *Rhoa*$^{fl/fl}$ Ang/PE vs *Rhoa*$^{\beta MHC\text{-}cre}$ Veh | Tukey one-way ANOVA | 0.8359 | -0.9792 to 2.651 | 0.8864 |

| | | | | |
|---|---|---|---|---|
| *Rhoa*$^{fl/fl}$ Ang/PE vs *Rhoa*$^{\beta MHC\text{-}cre}$ Ang/PE | Tukey one-way ANOVA | -0.8788 | -2.417 to 0.6591 | 0.6914 |
| *Rhoa*$^{fl/fl}$ Ang/PE vs *Rhoa*$^{fl/fl}$;*Rhoc*$^{-/-}$ Veh | Tukey one-way ANOVA | 1.304 | -0.6013 to 3.209 | 0.4437 |
| *Rhoa*$^{fl/fl}$ Ang/PE vs *Rhoa*$^{fl/fl}$;*Rhoc*$^{-/-}$ Ang/PE | Tukey one-way ANOVA | -5.193 | -7.008 to -3.378 | **<0.0001** |
| *Rhoa*$^{fl/fl}$ Ang/PE vs *Rhoa*$^{\beta MHC\text{-}cre}$;*Rhoc*$^{-/-}$ Veh | Tukey one-way ANOVA | 1.009 | -0.8963 to 2.914 | 0.7754 |
| *Rhoa*$^{fl/fl}$ Ang/PE vs *Rhoa*$^{\beta MHC\text{-}cre}$;*Rhoc*$^{-/-}$ Ang/PE | Tukey one-way ANOVA | -4.762 | -6.577 to -2.947 | **<0.0001** |
| *Rhoa*$^{\beta MHC\text{-}cre}$ Veh vs *Rhoa*$^{\beta MHC\text{-}cre}$ Ang/PE | Tukey one-way ANOVA | -1.715 | -3.475 to 0.04519 | 0.0624 |
| *Rhoa*$^{\beta MHC\text{-}cre}$ Veh vs *Rhoa*$^{fl/fl}$;*Rhoc*$^{-/-}$ Veh | Tukey one-way ANOVA | 0.468 | -1.620 to 2.557 | 0.9992 |
| *Rhoa*$^{\beta MHC\text{-}cre}$ Veh vs *Rhoa*$^{fl/fl}$;*Rhoc*$^{-/-}$ Ang/PE | Tukey one-way ANOVA | -6.029 | -8.036 to -4.022 | **<0.0001** |
| *Rhoa*$^{\beta MHC\text{-}cre}$ Veh vs *Rhoa*$^{\beta MHC\text{-}cre}$;*Rhoc*$^{-/-}$ Veh | Tukey one-way ANOVA | 0.173 | -1.915 to 2.262 | >0.9999 |
| *Rhoa*$^{\beta MHC\text{-}cre}$ Veh vs *Rhoa*$^{\beta MHC\text{-}cre}$;*Rhoc*$^{-/-}$ Ang/PE | Tukey one-way ANOVA | -5.598 | -7.605 to -3.592 | **<0.0001** |
| *Rhoa*$^{\beta MHC\text{-}cre}$ Ang/PE vs *Rhoa*$^{fl/fl}$;*Rhoc*$^{-/-}$ Veh | Tukey one-way ANOVA | 2.183 | 0.3300 to 4.035 | **0.0091** |
| *Rhoa*$^{\beta MHC\text{-}cre}$ Ang/PE vs *Rhoa*$^{fl/fl}$;*Rhoc*$^{-/-}$ Ang/PE | Tukey one-way ANOVA | -4.314 | -6.074 to -2.554 | **<0.0001** |
| *Rhoa*$^{\beta MHC\text{-}cre}$ Ang/PE vs *Rhoa*$^{\beta MHC\text{-}cre}$;*Rhoc*$^{-/-}$ Veh | Tukey one-way ANOVA | 1.888 | 0.03499 to 3.741 | **0.0423** |
| *Rhoa*$^{\beta MHC\text{-}cre}$ Ang/PE vs *Rhoa*$^{\beta MHC\text{-}cre}$;*Rhoc*$^{-/-}$ Ang/PE | Tukey one-way ANOVA | -3.884 | -5.643 to -2.124 | **<0.0001** |
| *Rhoa*$^{fl/fl}$;*Rhoc*$^{-/-}$ Veh vs *Rhoa*$^{fl/fl}$;*Rhoc*$^{-/-}$ Ang/PE | Tukey one-way ANOVA | -6.497 | -8.586 to -4.409 | **<0.0001** |

| | | | | |
|---|---|---|---|---|
| *Rhoa*$^{fl/fl}$;*Rhoc*$^{-/-}$ Veh vs *Rhoa*$^{βMHC-cre}$;*Rhoc*$^{-/-}$ Veh | Tukey one-way ANOVA | -0.295 | -2.462 to 1.872 | >0.9999 |
| *Rhoa*$^{fl/fl}$;*Rhoc*$^{-/-}$ Veh vs *Rhoa*$^{βMHC-cre}$;*Rhoc*$^{-/-}$ Ang/PE | Tukey one-way ANOVA | -6.066 | -8.155 to -3.978 | **<0.0001** |
| *Rhoa*$^{fl/fl}$;*Rhoc*$^{-/-}$ Ang/PE vs *Rhoa*$^{βMHC-cre}$;*Rhoc*$^{-/-}$ Veh | Tukey one-way ANOVA | 6.202 | 4.114 to 8.291 | **<0.0001** |
| *Rhoa*$^{fl/fl}$;*Rhoc*$^{-/-}$ Ang/PE vs *Rhoa*$^{βMHC-cre}$;*Rhoc*$^{-/-}$ Ang/PE | Tukey one-way ANOVA | 0.4308 | -1.576 to 2.437 | 0.9994 |
| *Rhoa*$^{βMHC-cre}$;*Rhoc*$^{-/-}$ Veh vs *Rhoa*$^{βMHC-cre}$;*Rhoc*$^{-/-}$ Ang/PE | Tukey one-way ANOVA | -5.771 | -7.860 to -3.683 | **<0.0001** |
| **Panel G — Fibrosis % (Ang/PE)** | | | | |
| **Comparison** | **Test** | **Mean Diff.** | **95% CI** | **Adjusted P** |
| Omnibus one-way ANOVA | One-way ANOVA | F(9,59)=16.72 | — | **<0.0001** |
| WT$^{βMHC-cre}$ Veh vs WT$^{βMHC-cre}$ Ang/PE | Tukey one-way ANOVA | -2.248 | -5.294 to 0.7987 | 0.3301 |
| WT$^{βMHC-cre}$ Veh vs *Rhoa*$^{fl/fl}$ Veh | Tukey one-way ANOVA | 0.1003 | -2.946 to 3.147 | >0.9999 |
| WT$^{βMHC-cre}$ Veh vs *Rhoa*$^{fl/fl}$ Ang/PE | Tukey one-way ANOVA | -2.231 | -5.462 to 1.000 | 0.4234 |
| WT$^{βMHC-cre}$ Veh vs *Rhoa*$^{βMHC-cre}$ Veh | Tukey one-way ANOVA | -0.2911 | -3.654 to 3.072 | >0.9999 |
| WT$^{βMHC-cre}$ Veh vs *Rhoa*$^{βMHC-cre}$ Ang/PE | Tukey one-way ANOVA | -5.575 | -8.497 to -2.652 | **<0.0001** |
| WT$^{βMHC-cre}$ Veh vs *Rhoa*$^{fl/fl}$;*Rhoc*$^{-/-}$ Veh | Tukey one-way ANOVA | 0.3143 | -3.475 to 4.103 | >0.9999 |
| WT$^{βMHC-cre}$ Veh vs *Rhoa*$^{fl/fl}$;*Rhoc*$^{-/-}$ Ang/PE | Tukey one-way ANOVA | -5.879 | -9.242 to -2.516 | **<0.0001** |
| WT$^{βMHC-cre}$ Veh vs *Rhoa*$^{βMHC-cre}$;*Rhoc*$^{-/-}$ Veh | Tukey one-way ANOVA | 0.6677 | -2.872 to 4.207 | 0.9998 |
| WT$^{βMHC-cre}$ Veh vs *Rhoa*$^{βMHC-cre}$;*Rhoc*$^{-/-}$ Ang/PE | Tukey one-way ANOVA | -7.405 | -10.94 to -3.865 | **<0.0001** |

| | | | | |
|---|---|---|---|---|
| WT$^{\beta\text{MHC-cre}}$ Ang/PE vs *Rhoa*$^{\text{fl/fl}}$ Veh | Tukey one-way ANOVA | 2.348 | -0.5017 to 5.198 | 0.1941 |
| WT$^{\beta\text{MHC-cre}}$ Ang/PE vs *Rhoa*$^{\text{fl/fl}}$ Ang/PE | Tukey one-way ANOVA | 0.0169 | -3.030 to 3.063 | >0.9999 |
| WT$^{\beta\text{MHC-cre}}$ Ang/PE vs *Rhoa*$^{\beta\text{MHC-cre}}$ Veh | Tukey one-way ANOVA | 1.957 | -1.229 to 5.143 | 0.5896 |
| WT$^{\beta\text{MHC-cre}}$ Ang/PE vs *Rhoa*$^{\beta\text{MHC-cre}}$ Ang/PE | Tukey one-way ANOVA | -3.327 | -6.044 to -0.6098 | **0.0059** |
| WT$^{\beta\text{MHC-cre}}$ Ang/PE vs *Rhoa*$^{\text{fl/fl}}$;*Rhoc*$^{-/-}$ Veh | Tukey one-way ANOVA | 2.562 | -1.071 to 6.195 | 0.3929 |
| WT$^{\beta\text{MHC-cre}}$ Ang/PE vs *Rhoa*$^{\text{fl/fl}}$;*Rhoc*$^{-/-}$ Ang/PE | Tukey one-way ANOVA | -3.631 | -6.817 to -0.4450 | **0.0138** |
| WT$^{\beta\text{MHC-cre}}$ Ang/PE vs *Rhoa*$^{\beta\text{MHC-cre}}$;*Rhoc*$^{-/-}$ Veh | Tukey one-way ANOVA | 2.915 | -0.4563 to 6.287 | 0.1462 |
| WT$^{\beta\text{MHC-cre}}$ Ang/PE vs *Rhoa*$^{\beta\text{MHC-cre}}$;*Rhoc*$^{-/-}$ Ang/PE | Tukey one-way ANOVA | -5.157 | -8.529 to -1.785 | **0.0002** |
| *Rhoa*$^{\text{fl/fl}}$ Veh vs *Rhoa*$^{\text{fl/fl}}$ Ang/PE | Tukey one-way ANOVA | -2.331 | -5.378 to 0.7153 | 0.2818 |
| *Rhoa*$^{\text{fl/fl}}$ Veh vs *Rhoa*$^{\beta\text{MHC-cre}}$ Veh | Tukey one-way ANOVA | -0.3913 | -3.577 to 2.795 | >0.9999 |
| *Rhoa*$^{\text{fl/fl}}$ Veh vs *Rhoa*$^{\beta\text{MHC-cre}}$ Ang/PE | Tukey one-way ANOVA | -5.675 | -8.392 to -2.958 | **<0.0001** |
| *Rhoa*$^{\text{fl/fl}}$ Veh vs *Rhoa*$^{\text{fl/fl}}$;*Rhoc*$^{-/-}$ Veh | Tukey one-way ANOVA | 0.2141 | -3.419 to 3.847 | >0.9999 |
| *Rhoa*$^{\text{fl/fl}}$ Veh vs *Rhoa*$^{\text{fl/fl}}$;*Rhoc*$^{-/-}$ Ang/PE | Tukey one-way ANOVA | -5.979 | -9.165 to -2.793 | **<0.0001** |
| *Rhoa*$^{\text{fl/fl}}$ Veh vs *Rhoa*$^{\beta\text{MHC-cre}}$;*Rhoc*$^{-/-}$ Veh | Tukey one-way ANOVA | 0.5674 | -2.804 to 3.939 | >0.9999 |
| *Rhoa*$^{\text{fl/fl}}$ Veh vs *Rhoa*$^{\beta\text{MHC-cre}}$;*Rhoc*$^{-/-}$ Ang/PE | Tukey one-way ANOVA | -7.505 | -10.88 to -4.133 | **<0.0001** |

| | | | | |
|---|---|---|---|---|
| *Rhoa*$^{fl/fl}$ Ang/PE vs *Rhoa*$^{βMHC-cre}$ Veh | Tukey one-way ANOVA | 1.94 | -1.423 to 5.303 | 0.6715 |
| *Rhoa*$^{fl/fl}$ Ang/PE vs *Rhoa*$^{βMHC-cre}$ Ang/PE | Tukey one-way ANOVA | -3.344 | -6.267 to -0.4210 | **0.0133** |
| *Rhoa*$^{fl/fl}$ Ang/PE vs *Rhoa*$^{fl/fl}$;*Rhoc*$^{-/-}$ Veh | Tukey one-way ANOVA | 2.545 | -1.244 to 6.334 | 0.4628 |
| *Rhoa*$^{fl/fl}$ Ang/PE vs *Rhoa*$^{fl/fl}$;*Rhoc*$^{-/-}$ Ang/PE | Tukey one-way ANOVA | -3.648 | -7.011 to -0.2848 | **0.0234** |
| *Rhoa*$^{fl/fl}$ Ang/PE vs *Rhoa*$^{βMHC-cre}$;*Rhoc*$^{-/-}$ Veh | Tukey one-way ANOVA | 2.899 | -0.6411 to 6.438 | 0.2008 |
| *Rhoa*$^{fl/fl}$ Ang/PE vs *Rhoa*$^{βMHC-cre}$;*Rhoc*$^{-/-}$ Ang/PE | Tukey one-way ANOVA | -5.174 | -8.714 to -1.634 | **0.0004** |
| *Rhoa*$^{βMHC-cre}$ Veh vs *Rhoa*$^{βMHC-cre}$ Ang/PE | Tukey one-way ANOVA | -5.284 | -8.352 to -2.216 | **<0.0001** |
| *Rhoa*$^{βMHC-cre}$ Veh vs *Rhoa*$^{fl/fl}$;*Rhoc*$^{-/-}$ Veh | Tukey one-way ANOVA | 0.6054 | -3.297 to 4.508 | >0.9999 |
| *Rhoa*$^{βMHC-cre}$ Veh vs *Rhoa*$^{fl/fl}$;*Rhoc*$^{-/-}$ Ang/PE | Tukey one-way ANOVA | -5.588 | -9.078 to -2.098 | **<0.0001** |
| *Rhoa*$^{βMHC-cre}$ Veh vs *Rhoa*$^{βMHC-cre}$;*Rhoc*$^{-/-}$ Veh | Tukey one-way ANOVA | 0.9588 | -2.702 to 4.619 | 0.9971 |
| *Rhoa*$^{βMHC-cre}$ Veh vs *Rhoa*$^{βMHC-cre}$;*Rhoc*$^{-/-}$ Ang/PE | Tukey one-way ANOVA | -7.114 | -10.77 to -3.453 | **<0.0001** |
| *Rhoa*$^{βMHC-cre}$ Ang/PE vs *Rhoa*$^{fl/fl}$;*Rhoc*$^{-/-}$ Veh | Tukey one-way ANOVA | 5.889 | 2.359 to 9.419 | **<0.0001** |
| *Rhoa*$^{βMHC-cre}$ Ang/PE vs *Rhoa*$^{fl/fl}$;*Rhoc*$^{-/-}$ Ang/PE | Tukey one-way ANOVA | -0.3042 | -3.372 to 2.764 | >0.9999 |
| *Rhoa*$^{βMHC-cre}$ Ang/PE vs *Rhoa*$^{βMHC-cre}$;*Rhoc*$^{-/-}$ Veh | Tukey one-way ANOVA | 6.242 | 2.982 to 9.503 | **<0.0001** |
| *Rhoa*$^{βMHC-cre}$ Ang/PE vs *Rhoa*$^{βMHC-cre}$;*Rhoc*$^{-/-}$ Ang/PE | Tukey one-way ANOVA | -1.83 | -5.091 to 1.430 | 0.7042 |

| | | | | |
|---|---|---|---|---|
| *Rhoa*$^{fl/fl}$;*Rhoc*$^{-/-}$ Veh vs *Rhoa*$^{fl/fl}$;*Rhoc*$^{-/-}$ Ang/PE | Tukey one-way ANOVA | -6.193 | -10.10 to - 2.291 | **0.0001** |
| *Rhoa*$^{fl/fl}$;*Rhoc*$^{-/-}$ Veh vs *Rhoa*$^{\beta MHC\text{-}cre}$;*Rhoc*$^{-/-}$ Veh | Tukey one-way ANOVA | 0.3534 | -3.702 to 4.409 | >0.9999 |
| *Rhoa*$^{fl/fl}$;*Rhoc*$^{-/-}$ Veh vs *Rhoa*$^{\beta MHC\text{-}cre}$;*Rhoc*$^{-/-}$ Ang/PE | Tukey one-way ANOVA | -7.719 | -11.77 to - 3.664 | **<0.0001** |
| *Rhoa*$^{fl/fl}$;*Rhoc*$^{-/-}$ Ang/PE vs *Rhoa*$^{\beta MHC\text{-}cre}$;*Rhoc*$^{-/-}$ Veh | Tukey one-way ANOVA | 6.546 | 2.886 to 10.21 | **<0.0001** |
| *Rhoa*$^{fl/fl}$;*Rhoc*$^{-/-}$ Ang/PE vs *Rhoa*$^{\beta MHC\text{-}cre}$;*Rhoc*$^{-/-}$ Ang/PE | Tukey one-way ANOVA | -1.526 | -5.186 to 2.135 | 0.9313 |
| *Rhoa*$^{\beta MHC\text{-}cre}$;*Rhoc*$^{-/-}$ Veh vs *Rhoa*$^{\beta MHC\text{-}cre}$;*Rhoc*$^{-/-}$ Ang/PE | Tukey one-way ANOVA | -8.072 | -11.90 to - 4.249 | **<0.0001** |
